# Lipopolysaccharide stimulates dynamic changes in B cell metabolism to promote proliferation

**DOI:** 10.1101/2024.12.16.628649

**Authors:** Dana M.S. Cheung, Momchil Razsolkov, Fabrizia Bonacina, Stephen Andrews, Megan Sumoreeah, Linda V. Sinclair, Andrew J.M. Howden, J. Simon C. Arthur

**Affiliations:** Division of Cell Signalling and Immunology, Faculty of Life Sciences, University of Dundee, Dow Street, Dundee, DD1; Department of Pharmacological and Biomolecular Sciences “Rodolfo Paoletti”, University of Milan, via Balzaretti 9, 20133, Milan, Italy

**Keywords:** B cell, cholesterol, transporter, toll-like receptor, statin

## Abstract

Naïve B cells exit quiescence and enter a proliferative state upon activation, ultimately differentiating into antibody-secreting or memory B cells. Toll-like receptor (TLR) ligands, such as lipopolysaccharide (LPS), can serve as physiological stimuli to initiate this transition. Using quantitative proteomics, we show that TLR4 engagement induces metabolic reprogramming in murine B cells, increasing the expression of amino acid transporters and cholesterol biosynthetic enzymes. The amino acid transporter SLC7A5 is markedly upregulated following LPS stimulation, and conditional deletion of *Slc7a5* impairs B cell proliferation, underscoring its essential role in B cell activation. LPS also elevates intracellular cholesterol levels, and inhibition of the rate-limiting enzyme HMG-CoA reductase blocks proliferation. This effect was mediated by a dual requirement for cholesterol metabolism and protein prenylation downstream of HMG-CoA reductase. Notably, this was not unique to TLR4 signalling but is also observed in B cells activated via TLR7, TLR9, CD40, or the B cell receptor. Together, these findings reveal that metabolic rewiring, including amino acid uptake and cholesterol metabolism, is an essential feature of B cell activation and proliferation.

## Introduction

B cell activation can be initiated through either T cell-dependent (TD) or T cell-independent (TI) pathways, with TI activation further subdivided into a type I or type II response depending on the stimuli. During TD activation, following recognition of a protein antigen via the B cell receptor (BCR), the B cell can present peptides from the antigen to CD4 T cells, resulting in T cell activation. Additionally, the B cell receives co-stimulation through the binding of its CD40 receptor to the CD40 ligand found on T cells, which promotes differentiation, proliferation, somatic hypermutation, and class switch recombination (CSR) (Hoffman *et al*, 2016). Type I TI responses are activated by agonists such as lipopolysaccharide (LPS), which bind to toll-like receptors (TLRs) on B cells to induce a strong proliferative signal and increase antibody production. Longer-term exposure to LPS can also promote the differentiation of plasma cells (Genestier *et al*, 2007). Conversely, type II TI antigens consist of long repetitive structures, such as polysaccharides, which at high concentrations crosslink multiple BCRs to stimulate proliferation and antibody production (Obukhanych & Nussenzweig, 2006).

A well-established model to study TI B cell activation is to use LPS, a component of the cell wall of gram-negative bacteria, which stimulates TLR4 (Hoshino *et al*, 1999; Poltorak *et al*, 1998) to induce robust B cell proliferation and antibody secretion, both *ex vivo* (Coutinho *et al*, 1974; Dziarski, 1982) and *in vivo* (Johnson *et al*, 1956). Upon activation, TLR4 propagates downstream signalling through the adaptor proteins MyD88 and TRIF. This in turn activates pathways, including MAPK and NF-κB signalling, to elicit the required physiological response (Arthur & Ley, 2013; Kawai *et al*, 2024). The essential role of TLR4 in LPS-induced activation of B cells has been demonstrated by studies using TLR4-deficient mice. B cells from TLR4 or Myd88 knockout mice did not proliferate in response to LPS or lipid A stimulation *ex vivo* (Hoshino *et al*, 1999; Kawai *et al*, 1999). Similarly, in studies where B cells from MyD88- and TLR4-knockout mice were transferred into µMT mice (that lack mature B cells), a significant reduction in IgG1 production was observed (Pasare & Medzhitov, 2005).

Naïve B cells have been shown to be in a quiescent state with low metabolic demand (Waters *et al*, 2018). Conversely, B cell activation must require a significant increase in biomolecules such as proteins, lipids, and nucleic acids to support their rapid expansion and proliferation. The existing literature has focused on the role of glucose and glucose transporters in B cell function. Previous studies have demonstrated that activation using LPS or anti-IgM to stimulate the BCR increases glucose uptake and leads to an increased oxygen consumption rate (Dufort *et al*, 2007; Jellusova *et al*, 2017). Whether this corresponds to changes in metabolic enzymes at a proteomic level, and whether TI B cell activation utilises other metabolic pathways, remains unknown. This increased glucose uptake has been shown to provide precursors for ribonucleotide and lipid synthesis (Waters *et al*, 2018). Similarly, the deletion of *Slc2a1* (which encodes for glucose transporter GLUT1) impaired germinal centre formation, plasma cell differentiation, and the production of high-affinity antibodies (Bierling *et al*, 2024; Brookens *et al*, 2023). Recent reviews have expressed the need for studies investigating metabolic changes in activated B cells, especially the role of lipid metabolism, which is poorly understood (Johnstone *et al*, 2024; Peeters & Jellusova, 2023).

T cell activation has been shown to stimulate global changes in metabolism at a proteomic level (Procaccini *et al*, 2016; Howden *et al*, 2019; Tan *et al*, 2017). However, there are no comparable proteomic studies that demonstrate how B cells support the change in metabolic requirements and energy demand from quiescence to activation upon LPS stimulation. Therefore, we used Data Independent Acquisition (DIA) based mass spectrometry to profile the B cell response to a combination of LPS + Interleukin 4 (IL-4) stimulation, a well-established *ex vivo* model of TI B cell activation. From this, we identified a substantial upregulation of proteins involved in multiple metabolic pathways, including amino acid uptake and cholesterol biosynthesis. We demonstrate the requirement of L-amino acid transporter SLC7A5 for B cell survival and proliferation *ex vivo*. Similarly, we highlight the necessity of both cholesterol metabolism and protein prenylation downstream of the rate-limiting enzyme HMG-CoA reductase for B cell growth, survival, and proliferation. These results highlight the pronounced metabolic rewiring induced downstream of LPS + IL-4 stimulation.

## Results

### LPS + IL-4 stimulation of murine B cells promotes proliferation and changes the expression of enzymes involved in cellular metabolism

Previous studies have shown that *ex vivo* stimulation of murine B cells with LPS and IL-4 can drive B cell proliferation and class switching to IgG1 (Snapper *et al*, 1988). Consistent with this, we found that lymphocytes isolated from lymph nodes and subsequently stimulated with LPS + IL-4 resulted in the proliferation of CD19+ B cells, as judged by dilution of Cell Trace Violet (CTV) labelling. LPS + IL-4 stimulation led to extensive proliferation, with up to 6 generations of B cells present after 72 hours (Figure 1A). Proliferation itself is thought to be a prerequisite for cells to undergo class-switching (Hodgkin *et al*, 1996). In line with this, only LPS + IL-4 stimulated B cells found in later generations were positive for IgG1 (Figure 1A). On a similar note, cell size is considered a key regulator of the G1/S transition, as progression through the cell cycle can only continue once a certain threshold in size has been reached (Barberis *et al*, 2007). Accordingly, we found that 72 hours of LPS + IL-4 stimulation led to a marked increase in cell size and granularity as indicated by a higher forward/side scatter compared to naïve B cells (Figure 1B-D). Stimulation with IL-4 alone resulted in a smaller increase in cell size than a combination of LPS + IL-4, but did not give rise to a strong proliferative response or class switching (Figure 1A-D). Naïve B cells were used for comparison in ongoing experiments, as surviving unstimulated B cells exhibited a similar phenotype in the flow cytometry analysis; however, the lack of stimulation led to significant cell death from 24 hours onwards (Figure S1A-B).

**Figure 1.**
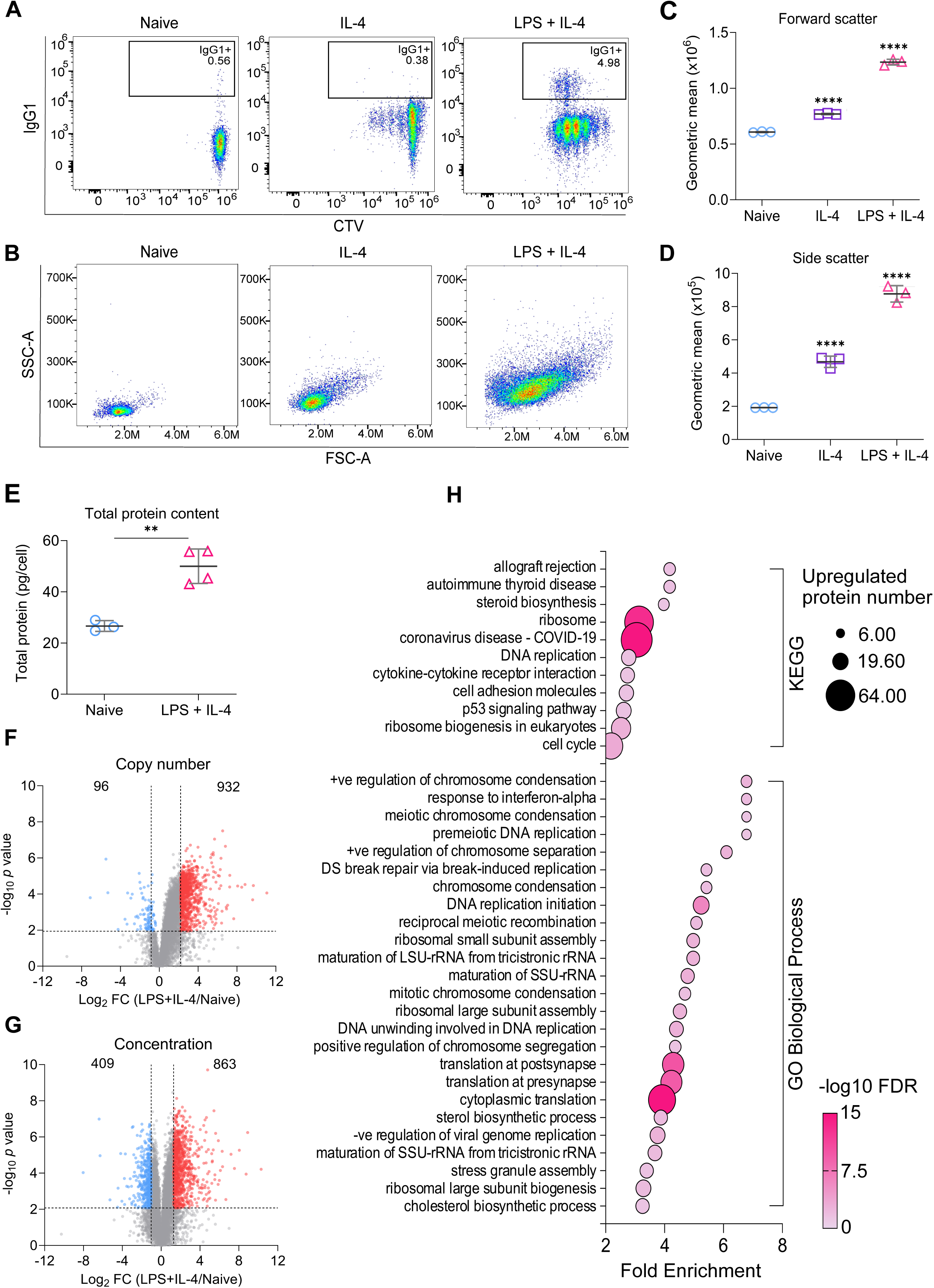
Stimulation with LPS + IL-4 promotes B cell proliferation and class switch recombination. **(A-D)** Cells from the lymph nodes of C57BL6/J mice were stained with Cell Trace Violet (CTV) and stimulated with IL-4 (10ng/ml) +/- LPS (20μg/ml) for 72 hours and analysed by flow cytometry. Data for naive B cells (stained on day 0) is also shown. Live CD19+ B cells were identified using the gating strategy described in Figure S14A. **(A)** Representative flow cytometry plots comparing IgG1 expression and CTV staining after 72 hours of IL-4 or LPS + IL-4 stimulation. **(B)** Representative plots for FSC and SSC at 72 hours of IL-4 or LPS + IL-4 stimulation with geometric means for the **(C)** forward scatter and **(D)** side scatter. Data shows the results of three biological replicates. Data was analysed by one-way ANOVA followed by multiple comparison testing via Sidak’s analysis. For comparisons to naïve B cells, *p*<0.0001 is indicated by ****. Full ANOVA results are given in Supplementary Table S5. **(E-H)** Cells from the lymph nodes of C57BL6/J mice, stimulated with LPS (20μg/ml) and IL-4 (10ng/ml) for 24 hours, and CD19+ B cells were isolated by FACS. Alternatively, naïve CD19+ B cells were sorted directly from *ex vivo* lymph node cells. Cells were lysed and analysed by proteomics as described in the methods. Samples from four mice for LPS + IL-4 and three for naive were generated. **(E)** Total protein content (pg/cell) was estimated from the proteomic data. Statistical power was determined using an unpaired two-tailed Student’s t-test, where *p*<0.01 is indicated by **. **(F)** Volcano plot depicting changes in estimated protein copy number in naïve vs LPS + IL-4 stimulated B cells. **(G)** Volcano plot showing the estimated cellular protein concentration (μM) in naïve vs LPS + IL-4 stimulated B cells. Horizontal dashed lines indicate *q*<0.05. Vertical dashed lines indicate log2 fold change of one standard deviation away from the median. **(H)** Enrichment analysis of the upregulated proteins in **(G)** against GO-term and KEGG databases.

These functional differences following activation suggest a significant change may be occurring at the protein level. To investigate this further, we analysed changes in the proteome after 24 hours of stimulation, as this time point was sufficient for B cells to increase in size but retain their synchrony in terms of the number of divisions they had undergone *ex vivo*. DIA-based mass spectrometry was used to compare the proteomes of naive B cells to activated B cells after 24 hours of LPS + IL-4 stimulation. The proteomic ruler method (Wiśniewski *et al*, 2014) was then used to estimate protein copy numbers and concentration. Using this approach, a total of 8030 protein species were identified, of which 7187 were quantified in both naïve and LPS + IL-4 stimulated B cells. Based on the mass spectrometry data, the predicted total protein mass in the B cells increased by approximately 2-fold following stimulation with a combination of LPS + IL-4 (Figure 1E). This change in protein mass was due to an increased copy number of most proteins in the cell (Figure 1F), with a median log2 fold increase of 1.04. To adjust for the increase in cell size, the fold change in protein concentration was analysed, as this provides an indication of what proteins are selectively upregulated or downregulated as opposed to those that scale with cell size.

For the data using estimated cellular concentrations, using cutoffs of *q* < 0.05 and a log2 fold change more than one standard deviation away from the median, 863 proteins were found to be upregulated following activation, while a further 301 were detected in all of the stimulated B cells but none of the naïve ones. 409 proteins were downregulated in the LPS + IL-4 stimulated B cells in comparison to naïve B cells, while a further 57 were detected in the naïve B cells, but none of the activated B cells (Figure 1G).

To identify the biological pathways that were affected by LPS + IL-4 stimulation, proteins that were classified as regulated using the criteria above were searched against Gene Ontology (GO) Biological Processes terms and *Kyoto Encyclopaedia of Genes and Genomes* (KEGG) databases. Enrichment analysis on the downregulated proteins did not highlight any specific processes. However, upregulated proteins were associated with terms related to the cell cycle, ribosome biogenesis, protein translation, and steroid biosynthesis (Figure 1H, Table S1).

While changes in the above processes would make sense in terms of cell proliferation, it was notable that changes in the glycolytic and oxidative phosphorylation pathways were not highlighted in this analysis. Previous studies have indicated that there is an increased demand for glucose in LPS-activated B cells (Dufort *et al*, 2007; Johnstone *et al*, 2024). Isotope tracing experiments following CD40L and IL-4 activation to stimulate proliferation have suggested that this increased glucose uptake is used in the pentose phosphate pathway, potentially to provide NADPH and precursors for nucleotide biosynthesis, while alternative carbon sources may be used to drive an increase in the TCA cycle and oxidative phosphorylation (Waters *et al*, 2018). The levels of enzymes involved in glucose metabolism were therefore examined in the proteomic data. In general, changes in the estimated concentrations of proteins involved in glycolysis, pentose phosphate pathway, TCA cycle, or oxidative phosphorylation were not observed (Figure S2A, B), although their copy numbers did increase in proportion to the increase in cell size (Figure S2C, D). B cells express the SLC2A1 (Glut1) glucose transporter, and deletion of this gene has been shown to reduce B cell activation *in vivo* (Bierling *et al*, 2024; Brookens *et al*, 2023). There was an increase in the concentration of SLC2A1, which would be consistent with an elevated ability to take up glucose (Figure S2E). In addition, there was a 13-fold increase in hexokinase 2 (HK2), one of the enzymes involved in catalysing the 1st committed step for glucose to enter the glycolytic or pentose phosphate pathways (Tanner *et al*, 2018) (Figure S2F) and has recently been shown to be required for maximal B cell responses to *ex vivo* stimulation with LPS (Paradoski *et al*, 2024). To investigate if the increased copy number of Electron Transport Chain proteins correlated with changes in mitochondrial volume, B cells were stained with MitoTracker Red, a fluorescent dye that binds to mitochondria. We found that MitoTracker staining was significantly increased following 24 hours of LPS + IL-4 stimulation (Figure S2G, H).

Notably, one of the enriched processes was identified as the ‘response to interferon alpha’. To take this further, we compared our data with a previously published RNAseq dataset that looked at the effect of IFNα on murine B cells (Mostafavi *et al*, 2016). We found that the majority of the IFNα regulated genes were not significantly upregulated at a protein level following LPS + IL-4 stimulation compared to naïve B cells (Figure S3A). IFNα and IFNβ signalling requires phosphorylation of the transcription factor STAT1 to induce transcription of type I IFN-dependent genes (Platanias, 2005). We found that LPS stimulation did not lead to detectable phosphorylation of STAT1 compared to stimulation with IFNβ (Figure S3B-C). Together, this suggests that there is not a strong interferon response in this system. It is possible, however, that as we stimulated lymph node cells before sorting for B cells for the proteomic experiment, a low level of type I interferon production from myeloid cells may have occurred, which could have contributed to a weak IFN gene signature in the B cells.

### LPS + IL-4 stimulation upregulates proteins involved in the cell cycle

Stimulation with LPS + IL-4 causes B cells to exit from a quiescent state and start to proliferate (Figure 1). This was confirmed using DAPI staining, which demonstrated that while naïve B cells were predominantly in a G0/1 state, after 24 hours of LPS + IL-4 stimulation, the proportion of B cells in the S and G2 stages of the cell cycle were increased (Figure 2A, B). Progression through G1 is associated with an increase in Cyclin D expression, and activation of CDK4 and 6 (Rubin *et al*, 2020). At the same time, inhibitory cell cycle proteins such as p27Kip1 (Cdkn1b) are downregulated (Wagner *et al*, 1998). Therefore, we analysed the proteomic data for evidence of these changes. In naïve B cells, Cyclin D was not detected, while levels of CDK4 and 6 were low (Figure 2C-E). The levels of Cyclin D, CDK4, and CDK6 were all increased following stimulation (Figure 2C-E). Naïve B cells also expressed high levels of p27Kip1, and this was reduced by LPS + IL-4 stimulation (Figure 2F).

**Figure 2.**
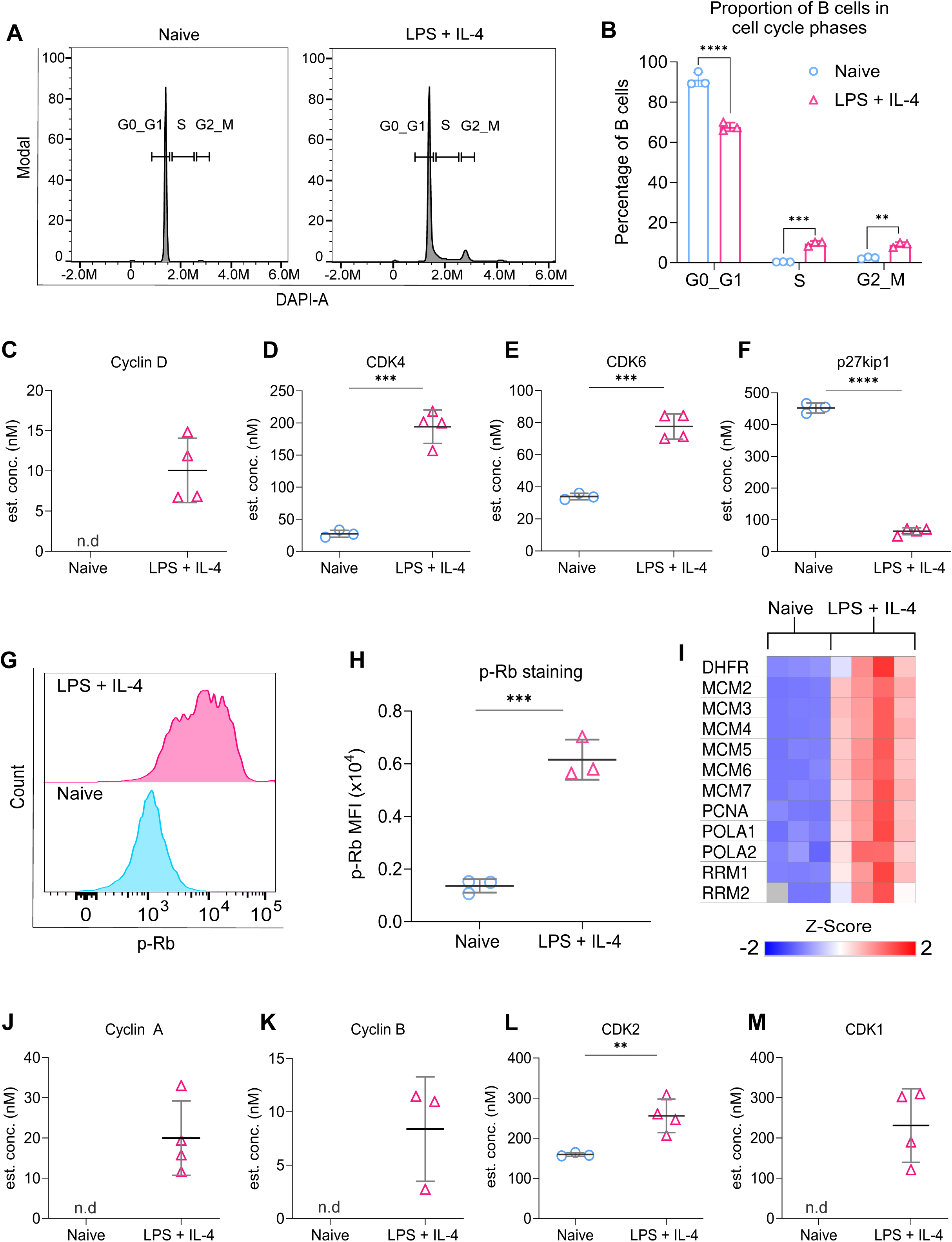
Proteins involved in cell cycle progression are upregulated in LPS + IL-4 activated B cells. **(A-B)** B cells were purified from the spleens of C57BL6/J mice and either fixed on isolation (naive) or stimulated with LPS (20μg/ml) and IL-4 (10ng/ml) for 24 hours before fixation. Cells were then stained with DAPI and CD19. The cell cycle stages were analysed using the gating strategy shown in Figure S14B. **(A)** Representative histograms showing the proportion of B cells in different phases of the cell cycle. **(B)** Quantification shows three technical replicates from cells isolated from one mouse and is representative of three independent experiments. Statistical power was determined using two-way ANOVA followed by multiple comparison testing via Sidak’s analysis, where *p*<0.01 is indicated by **, *p*<0.001 by *** and *p*<0.0001 by **** for comparisons between the naive and LPS + IL-4 conditions. **(C-F)** Graphs depicting changes in the estimated cellular concentration (nM) of proteins implicated in entry into the cell cycle, determined from the proteomic dataset described in Figure 1. **(C)** Cyclin D**, (D)** CDK4**, (E)** CDK6**, (F)** p27Kip1. **(G-H)** B cells were purified from the spleens of C57BL6/J mice and stimulated with LPS (20μg/ml) and IL-4 (10ng/ml) for 24 hours before fixing and staining for phospho-retinoblastoma (p-Rb). Gating strategy for p-Rb staining is shown in Figure S14B. **(G)** Representative histogram comparing p-Rb staining in naïve and LPS + IL-4 stimulated B cells. **(H)** Quantification shows three technical replicates from cells isolated from one mouse and is representative of three independent experiments. **(I)** Heat map showing the expression of proteins encoded by E2F target genes, derived from the proteomic data. **(J-M)** Graphs depicting changes in the estimated cellular concentration (nM) of proteins implicated in cell cycle progression, determined from the proteomic dataset described in Figure 1. **(J)** Cyclin A, **(K)** Cyclin B, **(L)** CDK2, **(M)** CDK1. p values were determined using an unpaired two-tailed Student’s t-test for **(H)** or represent adjusted p values from the FDR calculations applied to the proteomic dataset **(D, E, F, L)**, where *p*<0.01 is indicated by **, *p*<0.001 by *** and *p*<0.0001 by ****.

During the cell cycle, unphosphorylated RB1 binds to the E2F1/2/3 transcription factor, preventing the transcription of genes required for cell cycle progression. Increased CDK4/6 activity initiates RB1 phosphorylation, reducing its interaction with E2F, and resulting in the induction of E2F target genes, allowing progression through G1 to the S phase of the cell cycle (Engeland, 2022; Rubin *et al*, 2020). Increased phosphorylation of RB1 was observed in B cells following LPS + IL-4 stimulation (Figure 2G, H). In line with the increased RB1 phosphorylation, the levels of protein expressed from known E2F target genes, such as *Mcm2-7, Pcna, Dhfr, Rrm1/2,* and *Pola*3 (Chicas *et al*, 2010; Engeland, 2022; Leone *et al*, 1998; Wagner *et al*, 1998) were increased following LPS + IL-4 stimulation (Figure 2I). Further progression through the cell cycle requires Cyclins E, A, and B, as well as CDK2 and CDK1. While Cyclin E was not detected in the proteomic data, the levels of Cyclin A, Cyclin B, CDK2, and CDK1 were all increased by LPS + IL-4 stimulation (Figure 2J-M).

### Protein synthesis in LPS + IL-4 stimulated B cells is dependent on the uptake of amino acids

The cell growth and proliferation promoted by LPS + IL-4 stimulation would be expected to require an increased rate of protein synthesis. This is consistent with the proteomic analysis, which demonstrated an increase in total protein content within activated B cells (Figure 1E) and an enrichment of proteins associated with protein translation, including proteins that regulate ribosome biogenesis and the ribosomes themselves (Figure 1H). The ribosome is composed of 40S and 60S subunits, each of which is composed of multiple proteins (Doudna & Rath, 2002). LPS + IL-4 stimulation increased the total concentration of ribosomal proteins that make up both the 40S and 60S subunits (Figure 3A, B). This overall increase was due to higher expression of most of the individual proteins that make up the subunits, rather than a large increase in one or two of the proteins (Figure 3C, D). In line with the increase in ribosomal proteins, there was also an upregulation of proteins associated with ribosome biogenesis (Figure 3E-G). To determine if this correlated with an increased rate of protein synthesis, we measured the incorporation of the puromycin analogue o-propargyl-puromycin (OPP) into newly synthesised proteins. Consistent with the upregulation of ribosomal proteins, OPP incorporation was significantly higher in B cells stimulated with LPS + IL-4 for 24 hours compared to naive B cells (Figure 3H-I). Since LPS + IL-4 stimulation increased OPP incorporation, we tested whether this was a stimulus-specific effect or whether other stimuli known to activate B cell proliferation could increase protein synthesis. Murine B cells also respond to agonists of TLR7 (Weeratna *et al*, 2005) and TLR9 (Arunkumar *et al*, 2013; Yi *et al*, 1998). Both the TLR7 agonist Resiquimod and the TLR9 agonist CpG increased OPP incorporation in B cells (Figure S4A-B). Naive B cells can also be activated *ex vivo* by ligation of their BCR with anti-IgM antibodies or by stimulation with CD40 ligand (CD40L). Similar to TLR agonists, both stimuli also increased OPP incorporation in B cells (Figure S4A-B).

**Figure 3.**
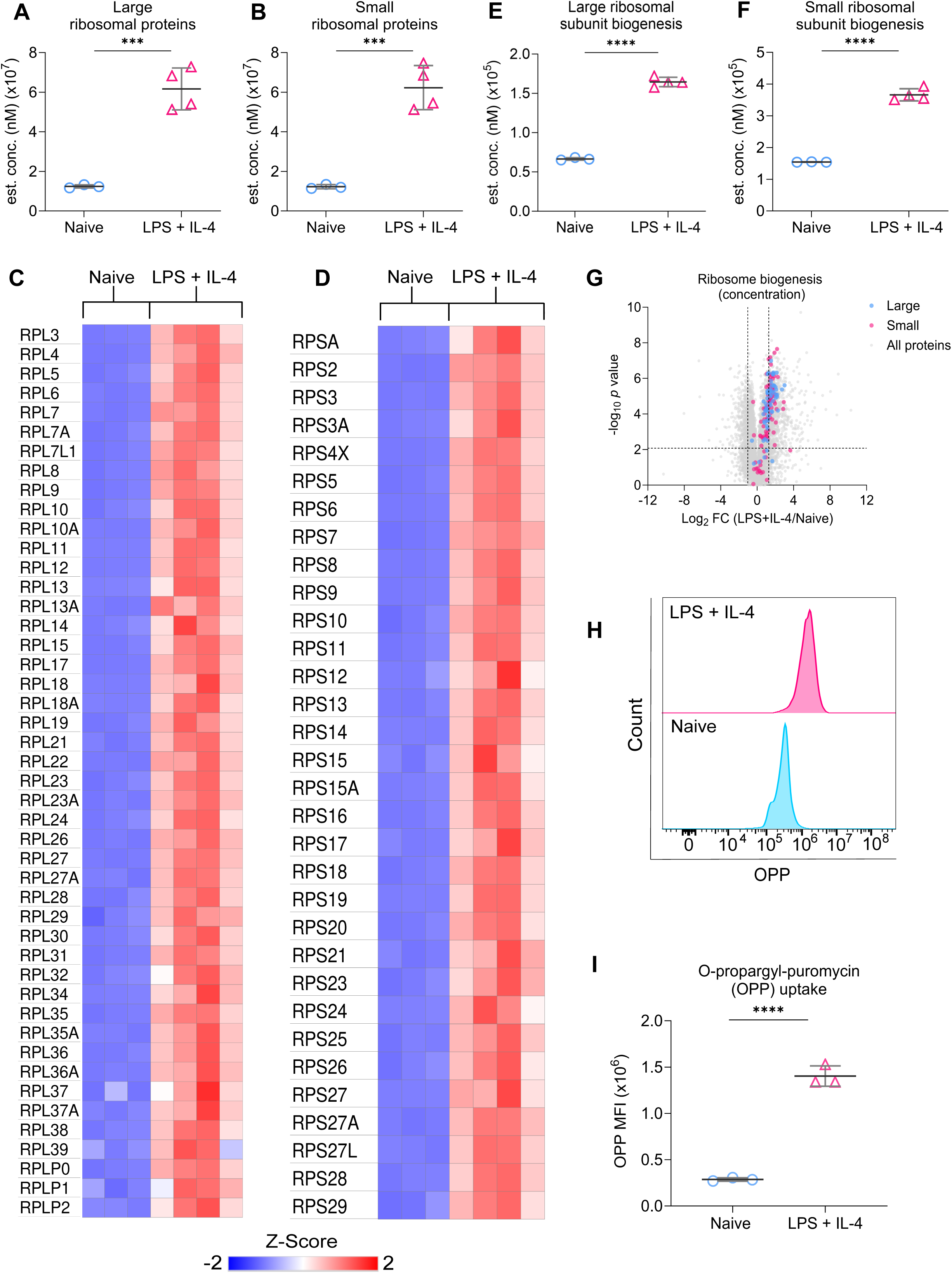
LPS + IL-4 stimulation promotes protein synthesis. **(A, B)** Graphs show changes in cellular concentration (nM) of the sum of proteins that make up the large **(A)** and small **(B)** ribosomal subunits, with heat maps showing the expression of the individual proteins making up the large **(C)** and small **(D)** subunits. **(E-G)** The proteomic dataset was mined for proteins involved in the biogenesis of the large and small ribosomal subunits based on the GO terms: GO:0000027, GO:0042273, GO:0000028, GO:0042274. **(E)** shows the sum of the proteins involved in the biogenesis of the large subunit and **(F)** shows the small subunit, with individual proteins represented on the volcano plot **(G)** Horizontal dashed lines on (G) indicate *q* < 0.05 while vertical dashed lines indicate log2 fold change more than one standard deviation away from the median. **(H-I)** Splenocytes from C57BL6/J mice were stimulated with LPS (20μg/ml) and IL-4 (10ng/ml) for 24 hours before fixing and staining for the uptake of puromycin analog O-propargyl-puromycin (OPP) to measure protein synthesis. Gating strategy for OPP staining is shown in Figure S14C. **(H)** Representative histogram comparing OPP uptake between naïve and LPS + IL-4 stimulated B cells. **(I)** Quantification shows three technical replicates from cells isolated from one mouse. Statistical power was determined using an unpaired two-tailed Student’s t-test, where *p*<0.001 is indicated by *** and *p*<0.0001 by ****.

Elevated protein synthesis during rapid proliferation would require an increased supply of amino acids. B cells have also been reported to increase amino acid biosynthesis upon stimulation. scRNAseq has shown that cycling B cells express higher levels of mRNA for enzymes in the serine biosynthetic pathway (D’Avola *et al*, 2022). Furthermore, the level of these proteins, as well as rates of serine biosynthesis, were found to be increased by stimuli such as anti-IgM, CD40L, and CpG, which are known to induce B cell proliferation (D’Avola *et al*, 2022). While this was not further addressed in the current study, the proteomic data reported here did show that serine biosynthetic enzymes, including the rate-limiting enzyme PHDGH, were increased in response to LPS + IL-4 stimulation (Figure S5).

Increased synthesis alone cannot satisfy the increased need for amino acids in proliferating cells, as the cell is unable to make essential amino acids. Several plasma membrane amino acid transporters were detected in the proteomic analysis, and these were all found to be upregulated in response to stimulation with LPS + IL-4 (Figure 4A). To look at the requirement for increased amino acid uptake during B cell proliferation more closely, we focused on Lat1, a member of the System L transporters. Lat1 is a heterodimer composed of an amino acid transporting light chain subunit, SLC7A5, and a chaperone heavy chain subunit, SLC3A2 (CD98) (Napolitano *et al*, 2015). Both of these subunits were upregulated in the proteomic data following LPS + IL-4 stimulation (Figure 4B, C). Additionally, the upregulation of SLC3A2 was confirmed by flow cytometry (Figure 4D, E). To determine whether this increased protein expression resulted from increased transcription of the relevant genes, we analysed RNA-Seq data generated from short-term LPS stimulation of murine B cells (Tesi et al, 2019). Both SLC7A5 and SLC3A2 mRNA expression was significantly upregulated following LPS stimulation (Figure S6A-B). A similar study using anti-IgM and anti-CD40 to activate murine B cells has found an upregulation of amino acid transporters, including SLC7A5, in their proteomic data, suggesting that this is not a stimulus-specific effect (James *et al*, 2024).

**Figure 4.**
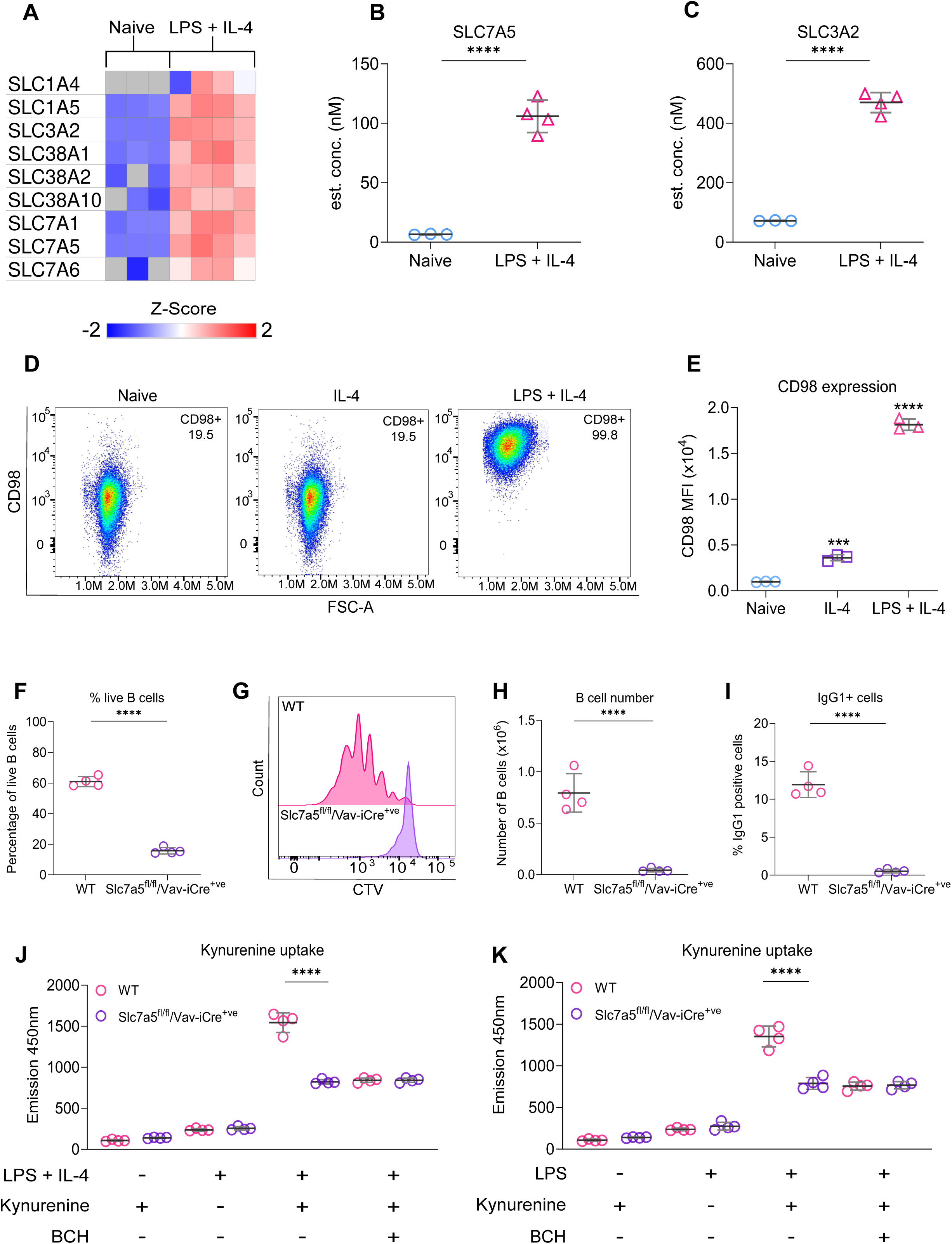
Amino acid transporter SLC7A5 is required for key B cell functions. **(A)** Heat map showing the expression of genes encoding for proteins involved in plasma membrane amino acid transport determined from the proteomic dataset described in Figure 1. **(B, C)** Graphs depicting changes in cellular concentration (nM) of **(B)** SLC7A5 and **(C)** SLC3A2 derived from the proteomic data. A*p(adj)* of <0.0001, based on the proteomic FDR calculations, is indicated by ****. **(D-E)** Lymph node cells from C57BL6/J mice were stimulated with LPS (20μg/ml) and IL-4 (10ng/ml) for 24 hours before staining for CD98. **(D)** Representative FACS plot comparing CD98 expression. **(E)** Quantification shows three technical replicates from cells isolated from one mouse and is representative of two independent experiments. Statistical power was determined using one-way ANOVA followed by multiple comparison testing via Dunnett’s analysis. For comparison to naïve B cells, where *p*<0.001 is indicated by *** and *p*<0.0001 by ****. **(F-I)** B cells were purified from the spleens of wild type (WT) and *Slc7a5* ^fl/fl^/Vav-iCre^+ve^ mice and stained with CTV before stimulation with LPS (20μg/ml) and IL-4 (10ng/ml) for 72 hours. **(F)** Percentage of live B cells (7AAD-ve) in WT and *Slc7a5*^fl/fl^/Vav-iCre^+ve^mice. **(G)** Histogram representing CTV staining of WT and *Slc7a5*^fl/fl^/Vav-iCre^+ve^ B cells. **(H)** Live B cell number of WT and *Slc7a5*^fl/fl^/Vav-iCre^+ve^mice. **(I)** Percentage of B cells that are IgG1+ve in WT and *Slc7a5*^fl/fl^/Vav-iCre^+ve^mice. Data shows the results of four biological replicates per genotype. *p*<0.0001 is indicated by **** (unpaired two-tailed Student’s t-test). **(J-K)** B cells were purified from the spleens of WT and *Slc7a5*^fl/fl^/Vav-iCre^+ve^mice and stimulated with LPS (20μg/ml) +/- IL-4 (10ng/ml) for 24 hours before fixing and staining for the uptake of kynurenine to measure amino acid uptake. Gating strategy for kynurenine uptake in Figure S14D. **(J)** Quantification of kynurenine MFI between B cells from WT and SLC7A5 KO mice with (+) or without (-) LPS + IL-4, kynurenine or aminobicyclo-(2,2,1)-heptane-2-carboxylic acid (BCH). **(K)** Quantification of kynurenine MFI between B cells from WT and *Slc7a5*^fl/fl^/Vav-iCre^+ve^mice with (+) or without (-) LPS, kynurenine or BCH. Data shows the results of four biological replicates per genotype. Statistical power was determined for using two-way ANOVA followed by multiple comparison testing via Sidak’s analysis, where *p*<0.0001 is indicated by **** for comparisons between genotypes.

*Slc7a5*^fl/fl^/Vav-iCre^+ve^ mice, which have a deletion of SLC7A5 in all hematopoietic cells, have a comparable number of naive B cells to wild-type (WT) controls, indicating that SLC7A5 is not critical for initial B cell development (Sinclair *et al*, 2013). To examine how the loss of SLC7A5 affects B cell proliferation in response to LPS + IL-4 stimulation, B cells were purified from WT and *Slc7a5*^fl/fl^/Vav-iCre^+ve^ mice, labelled with CTV, and stimulated for 72 hours with LPS + IL-4 *ex vivo*. The percentage of viable B cells after 72 hours was much lower in *Slc7a5*^fl/fl^/Vav-iCre^+ve^ mice compared to WT B cells (Figure 4F). Similarly, while WT B cells proliferated, the majority of the *Slc7a5*^fl/fl^/Vav-iCre^+ve^ B cells failed to divide, and the number of live B cells after 72 hours was much lower compared to WT B cells (Figure 4G, H). In line with the failure to proliferate, there was no evidence of class switching in the remaining *Slc7a5*^fl/fl^/Vav-iCre^+ve^ B cells (Figure 4I). SLC7A5 transports large neutral amino acids, including phenylalanine, histidine, methionine, tryptophan, and isoleucine, as well as the tryptophan metabolite kynurenine (Fotiadis *et al*, 2013). As kynurenine is fluorescent, and its uptake can be measured by flow cytometry, the effect of LPS + IL-4 treatment on kynurenine uptake in B cells was determined. In line with the increased expression of SLC7A5 in LPS + IL-4 treated B cells, kynurenine uptake was also higher in B cells stimulated with LPS + IL-4 for 24 hours compared to naive B cells (Figure 4J). Kynurenine uptake in the LPS + IL-4 stimulated B cells was reduced by the competitive substrate 2-aminobicyclo-(2,2,1)-heptane-2-carboxylic acid (BCH), which inhibits uptake through System L transporters. Kynurenine uptake in *Slc7a5*^fl/fl^/Vav-iCre^+ve^ B cells was similar to the uptake in BCH-treated WT cells, suggesting that SLC7A5 was the major system L transporter in LPS + IL-4 stimulated B cells (Figure 4J). BCH or SLC7A5 knockout did not reduce kynurenine uptake to baseline levels, indicating that other transport mechanisms for kynurenine uptake exist in B cells. Similar results were obtained for B cells stimulated with LPS alone, suggesting that LPS rather than IL-4 is the main driver of kynurenine uptake (Figure 4K).

### Cholesterol metabolism is upregulated in activated B cells

The enrichment analysis (Figure 1H) suggested that sterol biosynthesis may be upregulated by LPS + IL-4 stimulation. Sterol biosynthesis allows the generation of cholesterol from acetyl-CoA within the cell via a complex multistep process (Figure S7). The 1st half of this pathway, often referred to as the mevalonate pathway, results in the production of farnesyl pyrophosphate (Buhaescu & Izzedine, 2007). This can be used for several purposes in the cell, including the synthesis of cholesterol via the Bloch or Kandutsch-Russell pathways (Mitsche *et al*, 2015), prompting us to investigate this pathway further. Analysis of the proteomic data showed that the majority of enzymes involved in cholesterol biosynthesis were increased in LPS + IL-4 stimulated B cells relative to naive B cells (Figure 5A). Cholesterol biosynthesis is controlled by 2 rate-limiting steps: the conversion of HMG-CoA to mevalonic acid, which is catalysed by HMG-CoA-reductase (HMGCR), and the conversion of squalene to 2,3(*S*)-oxidosqualene, catalysed by squalene monooxygenase (SQLE) (Buhaescu & Izzedine, 2007). While neither of these enzymes were detected in the proteomic dataset for naive B cells, they were both consistently expressed after LPS + IL-4 stimulation (Figure 5B, C). Cells can also obtain cholesterol via the uptake of low-density lipoprotein (LDL) through the low-density lipoprotein receptor (LDLR) (Luo *et al*, 2020). Similar to HMGCR and SQLE, the LDLR was not found in naive B cells but was detected in B cells following LPS + IL-4 stimulation (Figure 5D). Analysis of published transcriptomic data (Tesi et al, 2019) on LPS stimulated B cells showed that *Hmgcr*, *Sqle* and *Ldlr* mRNA expression was significantly increased by LPS stimulation, and peaked after 4 hours of stimulation (Figures S6C-E).

**Figure 5.**
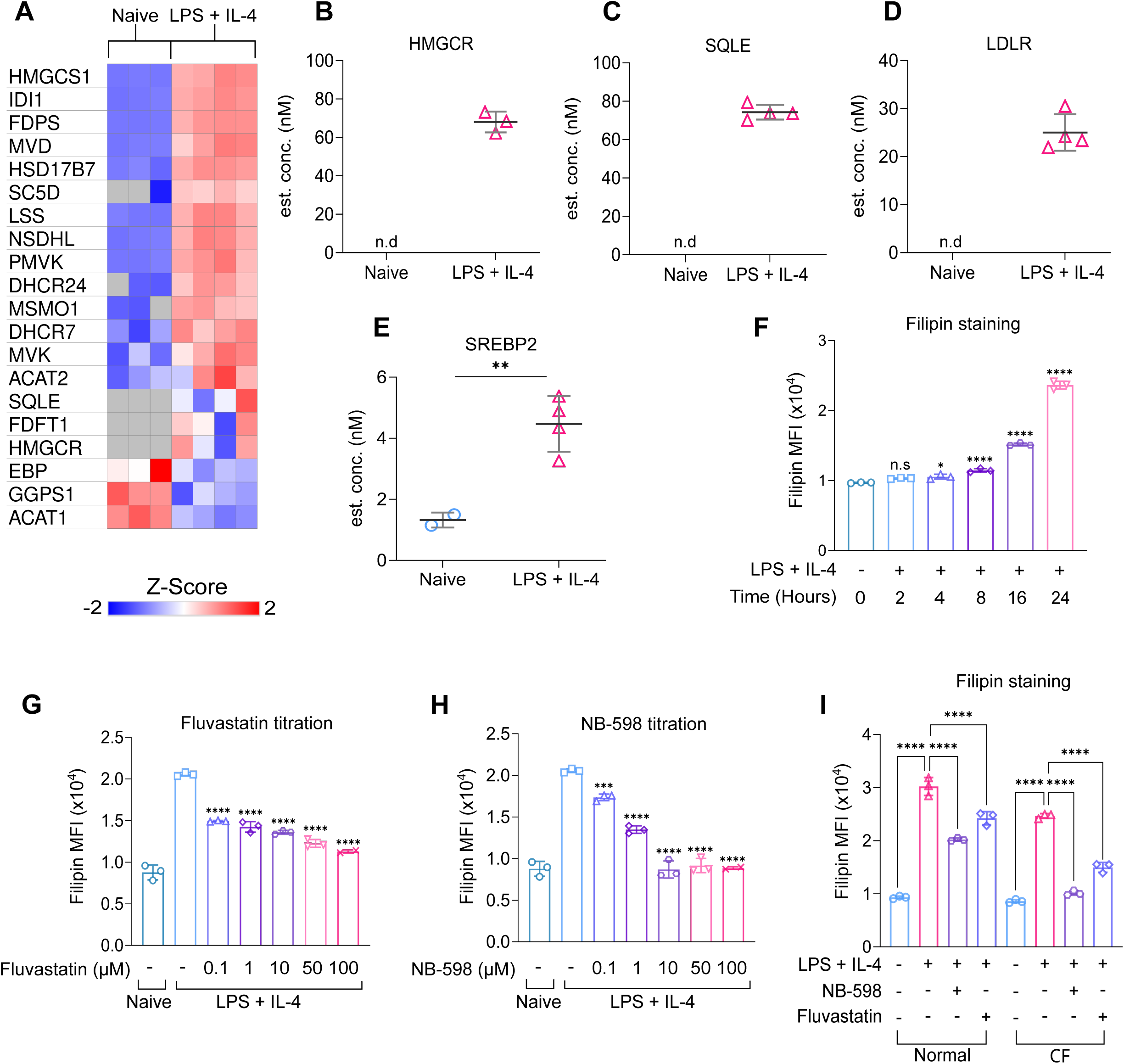
LPS + IL-4 stimulation upregulates cholesterol metabolism in B cells. **(A-E)** Analysis of the genes involved in cholesterol metabolism in the proteomic dataset. **(A)** Heat map of all the enzymes involved in cholesterol biosynthesis. Cellular concentration (nM) of **(B)** HMGCR, **(C)** SQLE, **(D)** LDLR, and **(E)**. A *p(adj)* of <0.01 is indicated by **. **(F)** Splenocytes from C57BL6/J mice were plated in cholesterol-free (CF) media and stimulated with LPS (20μg/ml) and IL-4 (10ng/ml). The cells were fixed at the stated time points and stained with filipin. The gating strategy for filipin staining in Figure S15A. **(F)** Filipin timecourse. Data shows three technical replicates from cells isolated from one mouse. Statistical power was determined by one-way ANOVA followed by multiple comparison testing via Dunnett’s analysis. For comparisons to naïve B cells, *p*<0.05 is indicated by * and *p*<0.0001 by ****. **(G-H)** Splenocytes from C57BL6/J mice were plated in cholesterol-free media and pre-treated with DMSO as a vehicle control or varying concentrations of Fluvastatin or NB-598 for 45 minutes before stimulation with LPS (20μg/ml) and IL-4 (10ng/ml). The cells were fixed after 24 hours and stained with filipin before acquisition. **(G)** Filipin staining of Fluvastatin titration. **(H)** Filipin staining of NB-598 titration. Data shows three technical replicates from cells isolated from one mouse and is representative of two independent experiments, each with one biological replicate. Statistical power was determined using a one-way ANOVA followed by multiple comparison testing via Dunnett’s analysis. For comparison to LPS + IL-4 stimulated B cells, *p*<0.001 is indicated by *** and *p*<0.0001 by ****. **(I)** Spleenocytes from C57BL6/J mice were plated in normal or cholesterol-free (CF) media and pre-treated with DMSO as a vehicle control, Fluvastatin (10μM) or NB-598 (10μM) before stimulation with LPS + IL-4. The cells were fixed after 24 hours and stained with filipin before acquisition. Data shows the results of three biological replicates. Statistical power was determined using two-way ANOVA followed by multiple comparison testing via Tukey’s analysis. For comparisons to the LPS + IL-4 condition, *p*<0.0001 is indicated by ****.

Cholesterol homeostasis is regulated by the transcription factor SREBP2, which is found as an immature precursor protein embedded in the endoplasmic reticulum (ER) with chaperones SCAP and INSIG (Luo *et al*, 2020; Brown *et al*, 2018). Low sterol levels are detected by the sterol-sensing domain of SCAP, leading to the dissociation and degradation of INSIG. Subsequently, the SCAP-SREBP2 complex is transported from the ER to the Golgi apparatus, where it interacts with site 1 and site 2 proteases, allowing cleavage of SREBP2. The mature form of SREBP2 then translocates to the nucleus, resulting in the upregulation of genes involved in cholesterol metabolism, including *hmgcr*, *sqle*, and *ldlr* (Luo *et al*, 2020; Brown *et al*, 2018). Low levels of SREBP2 expression were detectable in naïve B cells and became upregulated following LPS + IL-4 stimulation (Figure 5E). Conversely, there was no significant difference in the mRNA expression of *Srebp2* (Figure S6F), suggesting that the regulation of SREBP2 levels may occur post-transcriptionally.

Together, the above results suggest that LPS + IL-4 stimulation would drive an increase in cellular cholesterol. To test this, B cells were stained with the fluorescent dye filipin, which binds to cellular cholesterol. LPS + IL-4 stimulation resulted in a gradual increase in cholesterol levels from 4 hours to 24 hours of stimulation (Figure 5F). As cells can obtain cholesterol via *de novo* synthesis or uptake of LDL from serum, we compared the effects of culturing B cells in normal serum or serum treated with fumed silica to remove LDL (Brovkovych *et al*, 2019). Analysis of the serum indicated that this treatment successfully removed LDL and HDL but did not affect the levels of triglycerides (Table S2). Culture in media free of LDL (cholesterol-free (CF) media) did not prevent the increase in filipin staining stimulated by LPS + IL-4 (Figure 5G-I). Inhibitors of cholesterol biosynthesis were titrated to determine what concentration was required to reduce filipin staining in B cells stimulated in CF media. The increase in cholesterol following LPS + IL-4 stimulation could be reduced by the HMGCR inhibitor Fluvastatin or by NB-598, an inhibitor of SQLE (Figure 5G, H). Cells remained viable after 24 hours of inhibitor treatment with up to 100µM of either inhibitor (Figure S8A-F). Furthermore, the ability of HMGCR/SQLE inhibition to reduce cholesterol levels did not correlate with effects on cell size, as judged by FSC measurements by flow cytometry after 24 hours (Figure S8B, E). From this, a concentration of 10µM was chosen for both NB-598 and Fluvastatin, as this was sufficient to reduce cholesterol as close to naïve B cell levels as possible, but did not cause significant cell death after 24 hours. Different classes of statins have been described to exert different pleiotropic effects *in vivo* (Greenwood *et al*, 2006; Oesterle *et al*, 2017). Depending on their solubility, statins can be categorised as hydrophilic or lipophilic. Lipophilic statins, such as Fluvastatin, passively diffuse into cells, whereas hydrophilic statins may enter the cell through active transport. In the case of LPS + IL-4 stimulated B cells, the use of a more hydrophilic statin, Rosuvastatin, gave similar results to Fluvastatin (Figure S8G-J).

The increase in filipin staining following LPS + IL-4 stimulation for 24 hours in normal media was reduced, but not abolished, by either NB-598 or Fluvastatin treatment (Figure 5I). The ability of NB-598 and Fluvastatin treatment to reduce LPS + IL-4 induced filipin staining was greater in CF media relative to normal media that contains cholesterol (Figure 5I). Together, these results suggest that B cells may obtain cholesterol via a combination of biosynthesis and uptake.

### The mevalonate pathway is critical for the survival and proliferation of LPS + IL-4 stimulated B cells

As the above data indicated that B cells increased their cholesterol levels following LPS + IL-4 stimulation, we assessed the importance of this process for B cell proliferation *ex vivo*.

Stimulation with LPS + IL-4 for 48 hours in normal media promoted proliferation, and this was reduced by treatment with either Fluvastatin or NB-598 (as shown by CTV staining, Figure 6A). In line with this decreased proliferation, both Fluvastatin and NB-598 greatly reduced the number of B cells present at 48 hours compared to cells treated with LPS + IL-4 alone (Figure 6B). The remaining cells in the Fluvastatin or NB-598 treated conditions also showed significantly lower levels of survival (based on 7-AAD staining, Figure 6C) and failed to increase in size (as judged by forward scatter, Figure 6D). Similarly, B cells grown in CF media were still able to proliferate at a normal rate and experienced similar inhibitory effects after treatment with Fluvastatin or NB-598 (Figure 6A-D).

**Figure 6.**
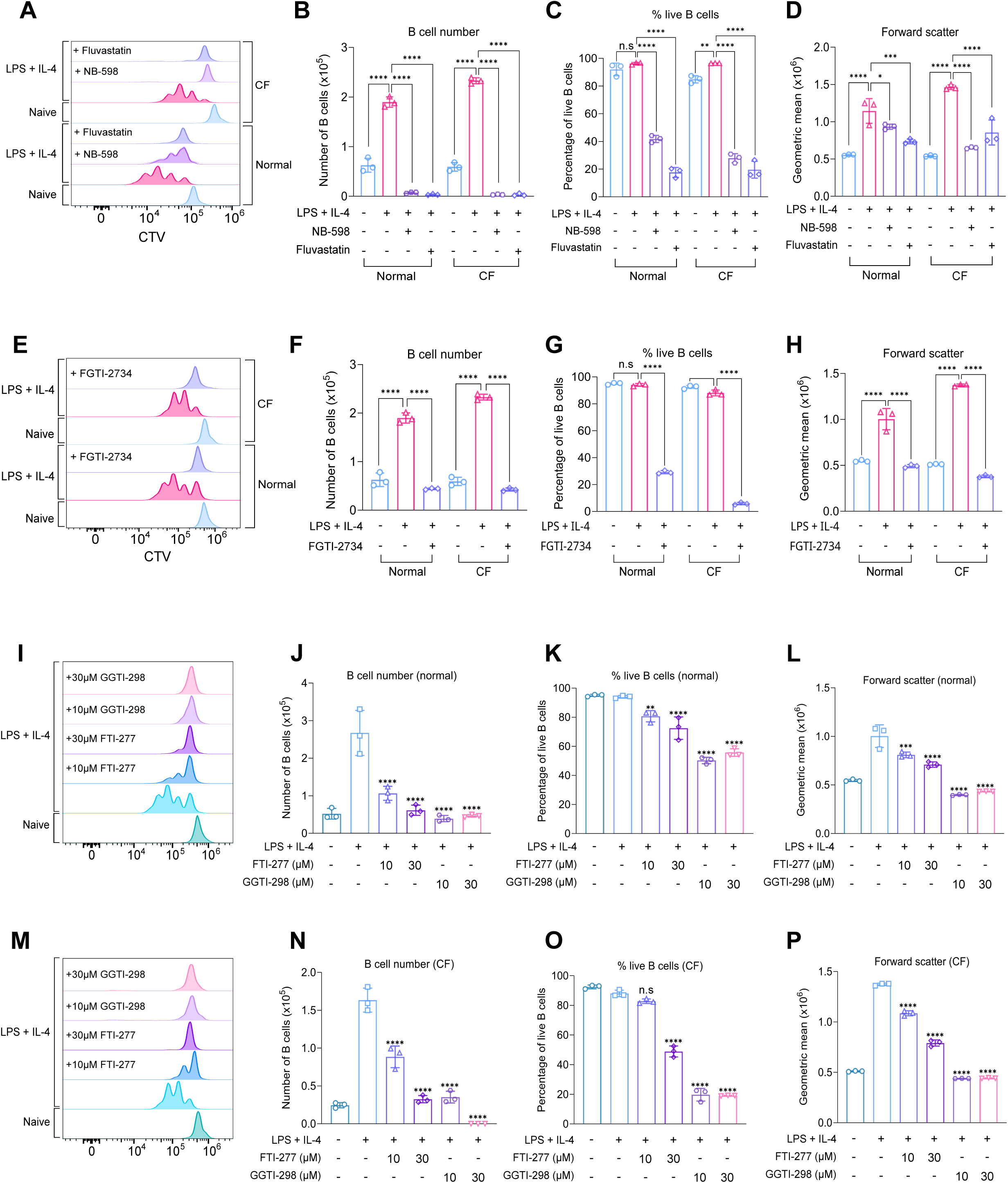
Blocking rate-limiting enzymes in the cholesterol biosynthesis pathway reduces B cell growth, survival and proliferation. **(A-D)** B cells were purified from the spleens of C57BL6/J mice and cultured in normal or cholesterol-free (CF) media. The cells were stained with CTV, then pre-treated with DMSO as a vehicle control or Fluvastatin (10µM) or NB-598 (10µM), where indicated, for 45 minutes prior to stimulation with LPS (20μg/ml) and IL-4 (10μg/ml) for 48 hours. Gating strategy for CTV staining is shown in Figure S15B. **(A)** shows representative CTV staining, **(B)** live B cell number, **(C)** percentage of live B cells (7AAD-ve) and **(D)** forward scatter. Data shows the results of three biological replicates. **(E-H)** Same as **(A-D)** but with pretreatment using FGTI-2734 (10µM). **(E)** shows representative CTV staining, **(F)** live B cell number, **(G)** percentage of live B cells (7AAD-ve) and **(H)** forward scatter of B cells. Data shows the results of three biological replicates. **(I-P)** Same as **(A-D)** but with pretreatment using FTI-277 or GGTI-298 (10µM-30µM). **(I)** shows representative CTV staining in normal media, **(J)** live B cell number, **(K)** percentage of live B cells (7AAD-ve) and **(L)** forward scatter of B cells. **(M)** shows representative CTV staining in CF media, **(N)** live B cell number, **(O)** percentage of live B cells (7AAD-ve) and **(P)** forward scatter of B cells. Data shows the results of three biological replicates. **(B-D) (F-H)** Statistical power was determined using two-way ANOVA followed by multiple comparison testing via Tukey’s analysis. For comparisons to the LPS + IL-4 condition, *p*<0.01 is indicated by **, *p*<0.001 by *** and *p*<0.0001 by ****. ns indicated by *p*>0.05. **(J-L) (N-P)** Statistical power was determined using one-way ANOVA followed by multiple comparison testing via Dunnett’s analysis. For comparisons to the LPS + IL-4 condition, *p*<0.01 is indicated by **, *p*<0.001 by *** and *p*<0.0001 by ****. ns indicated by *p*>0.05. For all panels, cells in the absence of LPS + IL-4 were naïve B cells analysed on the day of isolation.

To determine if blocking cholesterol biosynthesis or cholesterol uptake induced any major changes in the proteome, we analysed splenic B cells after 24 hours of LPS + IL-4 stimulation in normal and CF media in the presence or absence of Fluvastatin, using DIA-based mass spectrometry.

Using cutoffs of a log2 fold change more than one standard deviation away from the median and *q* < 0.05, no significant changes in protein expression were found when comparing LPS + IL-4 stimulated B cells cultured in normal media to those in CF media (Figure S9A). There was an upregulation of 31 proteins and a downregulation of 12 proteins when comparing LPS + IL-4 stimulated B cells grown in normal media with and without Fluvastatin (Figure S9B). The enrichment analysis (Table S1) did not highlight any specific processes in the downregulated proteins, but the upregulated proteins were enriched in terms associated with steroid biosynthesis. This suggests Fluvastatin treatment does not cause any major off-target effects in the B cells. Comparing LPS + IL-4 stimulated B cells grown in normal media to those in CF media + Fluvastatin, there was an upregulation of 32 proteins and downregulation of 35 proteins (Figure S9C). Again, the enrichment analysis (Table S1) did not highlight any specific processes in the downregulated proteins, but there was an enrichment for proteins involved in steroid biosynthesis in the upregulated proteins. Comparing upregulated proteins from Figure SF9B and C across all the conditions indicated that for most changes, the combination of CF media and Fluvastatin had the greatest effect (Figure SF9D). A similar pattern was observed for the downregulated proteins (Figure SF9E). Looking specifically at proteins involved in cholesterol metabolism, the majority of these were upregulated by treatment with Fluvastatin or growth in CF media (Figure S9F), including the rate-limiting enzymes HMGCR and SQLE (Figure S9G, H). In addition, there was upregulation of the LDLR (Figure S9I). Again, the biggest effect on these proteins was seen with the combination of CF media and Fluvastatin.

In addition to cholesterol biosynthesis, the mevalonate pathway is also required for protein prenylation, a process that involves the addition of a farnesyl or geranylgeranyl motif to cysteine residues near the C-terminus of target proteins

(Figure S7) (Wang & Casey, 2016). As SQLE acts downstream of this branch point, the effects of NB-598 would argue for an important role of cholesterol biosynthesis in LPS + IL-4 induced proliferation; however, they would not exclude a parallel role for protein prenylation. To determine if the inhibitory effects of Fluvastatin in stimulated B cells may be in part due to changes in prenylation, B cells were treated with a dual prenylation inhibitor, FGTI-2734 (Kazi *et al*, 2019), which blocks the enzymes responsible for catalysing prenylation; farnesyl transferase (FTase) and geranylgeranyl transferase (GGTase). Treatment of B cells with FGTI-2734 for 48 hours decreased proliferation in response to LPS + IL-4 stimulation in normal media (Figure 6E), and in line with this, the number of live B cells also decreased (Figure 6F). Cell survival was significantly reduced to the same level as seen with Fluvastatin treatment (compare Figure 6G to 6C). FGTI-2734 also had a major impact on cell size in normal media (Figure 6H). Similar effects were observed in CF media (Figure 6E-6H). To determine whether inhibiting farnesylation or geranylgeranylation was responsible for the inhibitory effect of FGTI-2734, B cells were treated with the FTase inhibitor FTI-277 (Lerner *et al*, 1995) or the GGTase inhibitor GGTI-298 (McGuire *et al*, 1996). In normal media, treatment with 10µM of FTI-277 reduced proliferation to around two generations of B cells (Figure 6I), whereas 30µM of FTI-277 had a more significant effect on proliferation (Figure 6I).

Inhibition of GGTase with either 10µM or 30µM of GGTI-298 blocked proliferation entirely (Figure 6I). In line with this, the number of live B cells was significantly reduced by treatment with either inhibitor (Figure 6J). In terms of survival, FTI-277 reduced the percentage of live B cells compared to the LPS + IL-4 stimulation alone, whereas GGTI-298 had a more profound impact on survival (Figure 6K). Similarly, treatment with FTI-277 reduced cell size, but GGTI-298 reduced cell size to a level comparable to naïve B cells (Figure 6L). Similar effects were observed in CF media (Figure 6M-6P); however, treatment with GGTI-298 reduced survival to an even greater extent compared to normal media (Figure 6O).

In theory, the effects of HMGCR inhibition should be reduced if cells were supplemented with a metabolite downstream of HMGCR in the mevalonate pathway. To test this, the effects of supplementing growth medium with mevalonate, the product generated in the reaction catalysed by HMGCR, were tested. For this, a concentration of 2mM mevalonate was used, as initial experiments showed that higher concentrations resulted in cell death and reduced filipin staining (Figure S10A-B). Initially, the ability of mevalonate to rescue the effects of Fluvastatin on filipin staining was examined. In cells treated for 48 hours with LPS + IL-4, the reduction in filipin caused by Fluvastatin treatment in normal media was reversible by the addition of mevalonate (Figure 7A). Mevalonate had less effect on cells stimulated for 24 hours with LPS + IL-4, although at this time point, the ability of Fluvastatin to reduce filipin staining was also less pronounced than at 48 hours (Figure 7A). Surprisingly, in CF media, mevalonate was unable to rescue the effects of Fluvastatin at either time point (Figure 7B).

**Figure 7.**
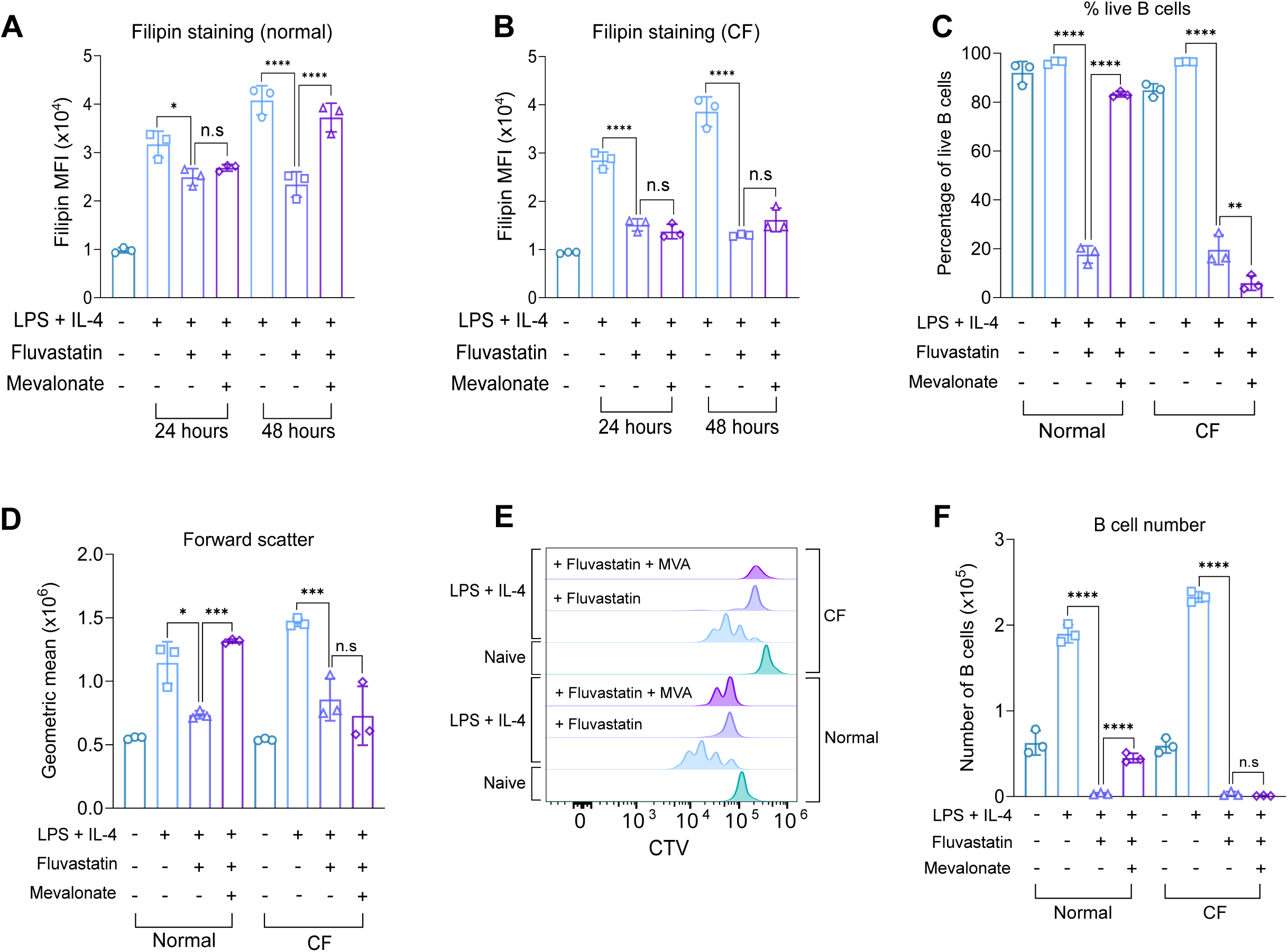
Mevalonate supplementation rescues the effect of Fluvastatin treatment in B cells. **(A-B)** Splenocytes from C57BL6/J mice were plated in normal or cholesterol-free (CF) media and pre-treated with HEPES as a vehicle control or mevalonate (2mM) for 1 hour prior to treatment with DMSO or Fluvastatin (10µM), where indicated, for 45 minutes before stimulation with LPS (20μg/ml) and IL-4 (10ng/ml). The cells were fixed after 24 hours or 48 hours of LPS + IL-4 stimulation and stained with filipin. **(A)** Filipin staining comparing B cells +/- Fluvastatin or mevalonate after 24 or 48 hours of LPS + IL-4 stimulation in normal media. **(B)** Filipin staining comparing B cells +/- Fluvastatin or mevalonate after 24 or 48 hours of LPS + IL-4 stimulation in CF media. Data shows the results of three biological replicates. **(C-F)** B cells were purified from the spleens of C57BL6/J mice and cultured in normal or cholesterol-free media. The cells were stained with CTV, then pre-treated with HEPES as a vehicle control or mevalonate (2mM) for 1 hour. The cells were treated with DMSO or Fluvastatin (10µM), where indicated, for 45 minutes prior to stimulation with LPS (20μg/ml) and IL-4 (10μg/ml) for 48 hours. **(C)** Shows percentage of live B cells (7AAD-ve), **(D)** forward scatter of B cells, **(E)** representative CTV staining and **(F)** live B cell number. Data shows the results of three biological replicates. Where shown, statistical power was determined using two-way ANOVA followed by multiple comparison testing via Sidak’s analysis. For comparison to Fluvastatin treated B cells, *p*<0.05 is indicated by *, *p*<0.01 by **, *p*<0.001 by ***, *p*<0.0001 by ****. ns by >0.05. For all panels, cells in the absence of LPS + IL-4 were naïve B cells analysed on the day of isolation.

The effect of Fluvastatin on cell survival was partially reversible by the addition of mevalonate in normal media, which restored cell survival to 80% (Figure 7C) and fully restored cell size relative to cells stimulated with LPS and IL-4 in the absence of Fluvastatin (Figure 7D). However, mevalonate supplementation was not sufficient to completely rescue proliferation, as CTV staining demonstrated that 2 generations of B cells were present rather than 4 generations after LPS + IL-4 stimulation alone (Figure 7E). The number of live B cells following mevalonate addition was significantly higher compared to the Fluvastatin only condition, but did not reach the same level as LPS + IL-4 stimulation alone (Figure 7F). Similar to what was seen for filipin staining, mevalonate was unable to rescue any of the inhibitory effects of Fluvastatin treatment in CF media (Figure 7C-F).

Since inhibiting geranylgeranyl transferase had such a profound effect on B cell function, the role of prenylation was also tested by supplementing cells with geranylgeranyl pyrophosphate (GGPP), which can be used for prenylation but not cholesterol biosynthesis. Surprisingly, GGPP was more effective than mevalonate in rescuing the effect of Fluvastatin on LPS + IL-4 induced proliferation (Figure 8A), and in line with this, there was a greater number of live B cells in the GGPP treated condition relative to the mevalonate treated condition (Figure 8B). A combination of GGPP and mevalonate did not have a synergistic effect compared to GGPP alone (Figure 8A, B). GGPP was also able to rescue the effects of Fluvastatin on both cell survival and cell size (Figure 8C, D). Conversely, in CF media, the addition of GGPP was unable to rescue any of the inhibitory effects of Fluvastatin on cell proliferation or survival (Figure 8E-H).

**Figure 8.**
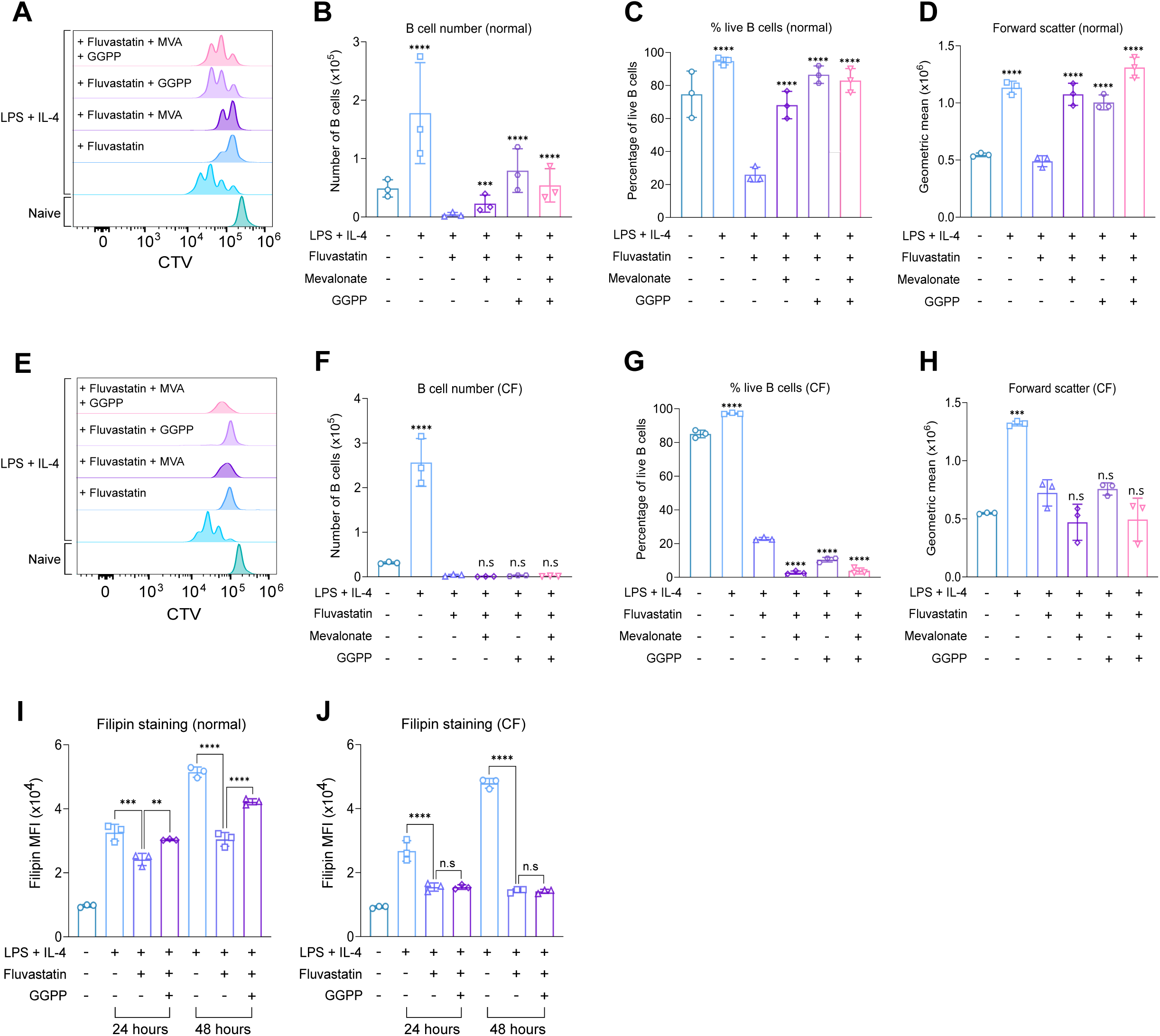
GGPP supplementation rescues the effect of Fluvastatin treatment in B cells. **(A-H)** B cells were purified from the spleens of C57BL6/J mice and cultured in normal or cholesterol-free media. The cells were stained with CTV, pre-treated with methanol:ammonium hydroxide solution (CH3OH:NH4OH) as a vehicle control or geranylgeranyl pyrophosphate (GGPP) (10μM) for 1 hour before treatment with Fluvastatin (10μM), where indicated. The cells were stimulated with LPS (20μg/ml) and IL-4 (10μg/ml) and cultured for 48 hours. **(A)** shows representative CTV staining for B cells cultured in normal media, **(B)** live B cell number, **(C)** percentage of live B cells and **(D)** forward scatter of B cells. **(E)** shows representative CTV staining for B cells cultured in CF media, **(F)** live B cell number, **(G)** percentage of live B cells and **(H)** forward scatter of B cells. Data shows the results of three biological replicates. Statistical power was determined using one-way ANOVA followed by multiple comparison testing via Dunnett’s analysis. For comparison to Fluvastatin treated B cells, *p*<0.001 is indicated by *** and *p*<0.0001 by ****. ns indicated *p*>0.05. For **(B)** and **(F)** this data was log-transformed then statistically analysed due to unequal variance. **(I-J)** Splenocytes were plated in normal or cholesterol-free media and pre-treated with methanol:ammonium hydroxide solution (CH3OH:NH4OH) as a vehicle control or geranylgeranyl pyrophosphate (GGPP) (10μM), where indicated, for 1 hour before treatment with DMSO or Fluvastatin (10μM). The cells were then stimulated with LPS (20μg/ml) and IL-4 (10ng/ml). The cells were fixed after 24 hours and stained with filipin before acquisition. **(I)** Filipin staining comparing B cells +/- Fluvastatin or GGPP after 24 or 48 hours of LPS + IL-4 stimulation in normal media. **(J)** Filipin staining comparing B cells +/- Fluvastatin or GGPP after 24 or 48 hours of LPS + IL-4 stimulation in CF media. Data shows the results of three biological replicates. Statistical power was determined using two-way ANOVA followed by multiple comparison testing via Sidak’s analysis. For comparison to Fluvastatin treated B cells, *p*<0.01 is indicated by **, *p*<0.001 by *** and *p*<0.0001 by ****. ns indicated *p*>0.05. For all panels, cells in the absence of LPS + IL-4 were naïve B cells analysed on the day of isolation.

Given that GGPP should rescue prenylation but not cholesterol metabolism, its effects on proliferation were unexpected if B cells need cholesterol to proliferate. We therefore tested if GGPP affected cholesterol levels in Fluvastatin treated B cells. As seen in previous experiments, Fluvastatin reduced the ability of LPS + IL-4 stimulation to increase cholesterol levels at both 24 hours and 48 hours of treatment. GGPP was able to partially rescue this at both time points (Figure 8I). This was in contrast to mevalonate, which could only rescue at 48 hours (Figure 7A), an observation that may, in part, explain why GGPP was more effective at restoring proliferation in the presence of Fluvastatin than mevalonate. The effects of GGPP on cholesterol levels were, however, limited to normal media, as similar to mevalonate, GGPP did not rescue the effect of Fluvastatin on filipin staining in CF media (Figure 8J).

### MAPK and mTOR signalling are required for B cell proliferation and cholesterol metabolism

To investigate how LPS induces these metabolic changes, we examined signalling pathways activated downstream of TLR4. MyD88 is a key adaptor protein which links most TLRs, including TLR4, to downstream signalling proteins (Kawai *et al*, 2024).

To confirm the necessity of this protein for LPS signalling, we stimulated B cells from WT and MyD88 knockout (KO) mice with LPS for 30 minutes before immunoblotting for the phosphorylation of downstream proteins. Knockout of MyD88 prevented the phosphorylation of p38α and ERK1/2. LPS also activates PI3K/mTOR pathways in B cells, and loss of Myd88 reduced the phosphorylation of AKT, p70S6K and S6 in these pathways (Figure S11).

To determine if these signalling pathways were important for LPS-induced B cell proliferation, B cells were treated with inhibitors of MAPK and mTOR signalling before stimulation with LPS + IL-4 for 48 hours. B cells treated with PD184352, an inhibitor of MEK1/2 (Bain *et al*, 2007), the upstream activators of ERK1/2, were still able to proliferate, with 4 generations of B cells present (Figure 9A). However, looking at the proportion of B cells per generation revealed that a higher percentage of B cells treated with PD184352 remained in generation 1 with a smaller percentage in generation 4, compared to the LPS + IL-4 stimulation alone (Figure 9B).

**Figure 9.**
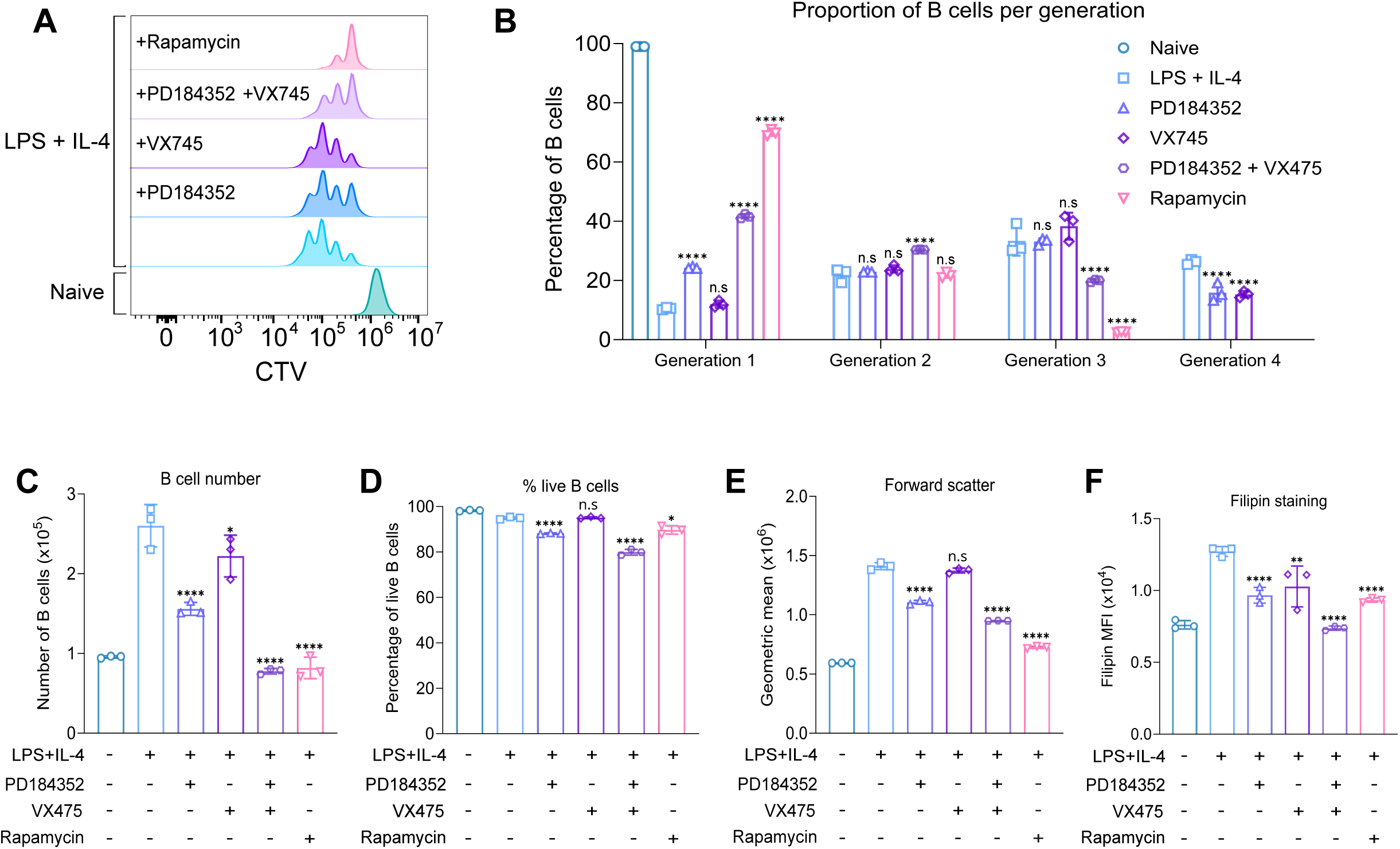
MAPK and mTOR signalling regulate B cell proliferation and cholesterol levels. **(A-E)** B cells were purified from the spleens of C57BL6/J mice and stained with CTV, then pre-treated with DMSO as a vehicle control, PD18352 (2μM), VX745 (1μM) or Rapamycin (20nM), where indicated, for 45 minutes prior to stimulation with LPS (20μg/ml) and IL-4 (10μg/ml) for 48 hours. **(A)** shows representative CTV staining, **(B)** percentage of B cells per generation quantified from **(A)**, **(C)** live B cell number **(D)** percentage of live B cells (7AAD-ve) and **(E)** forward scatter. Data shows three technical replicates from cells isolated from one mouse and is representative of three independent experiments. Statistical power for **(B)** was determined using two-way ANOVA followed by multiple comparison testing via Sidak’s analysis. For comparison to LPS + IL-4 stimulated B cells, *p*<0.0001 by **** and ns by >0.05. **(F)** Splenocytes from C57BL6/J mice were pre-treated with DMSO as a vehicle control, PD18352 (2μM), VX745 (1μM) or Rapamycin (20nM), where indicated, for 45 minutes prior to stimulation with LPS (20μg/ml) and IL-4 (10μg/ml). The cells were fixed after 24 hours and stained with filipin before acquisition. **(F)** Filipin staining after inhibitor treatment. Statistical power was determined by one-way ANOVA followed by multiple comparison testing via Dunnett’s analysis, where p<0.05 is indicated by *, p<0.01 by **, p<0.001 by ***, p<0.0001 by **** and ns by >0.05. For all panels, cells in the absence of LPS + IL-4 were naïve B cells analysed on the day of isolation.

Treatment with VX745, a selective inhibitor of p38α and β (McGuire *et al*, 2013), had a lesser effect on proliferation, with the only difference being a lower percentage in generation 4 (Figure 9A-B). To determine the impact of inhibiting both p38/MEK1/2 signalling, we treated B cells with a combination of both PD184352 and VX745, which reduced proliferation to 3 generations of B cells with a higher proportion of B cells in generations 1 and 2 (Figure 9A, B). Inhibition of the mTOR pathway using rapamycin had the greatest effect on proliferation out of all the compounds tested, resulting in 70% of B cells remaining in generation 1, with few cells progressing past generation 2 (Figure 9A-B). This correlated with the number of live B cells, with combined MAPK inhibition and mTOR inhibition resulting in the biggest impact on B cell number (Figure 9C). Interestingly, most of the inhibitor treatments had minimal impact on the percentage of live cells at 48 hours, although treatment with PD184352 reduced the percentage of live cells by 7% and the combination of PD184352 and VX745 by 15% (Figure 9D). In terms of cell size, treatment with PD184352, the combination of PD184352 and VX745, and rapamycin reduced cell size (Figure 9E). Treatment with all of the inhibitors reduced the levels of filipin staining, with combined PD184352 and VX745 treatment having the most significant effect on cholesterol levels (Figure 9F). Together, this would suggest multiple signalling pathways feed into the regulation of cholesterol in LPS activated B cells.

Interestingly, while the mTOR pathway is strongly linked to the regulation of protein synthesis (Ma & Blenis, 2009), rapamycin reduced but did not block protein synthesis and amino acid uptake in B cells stimulated with LPS + IL-4 (Figure S12A, B).

### The mevalonate pathway is required for B cell proliferation in response to multiple agonists

While the above results indicate that cholesterol is required for B cell proliferation in response to LPS + IL-4 stimulation, they do not demonstrate whether the primary signal for this is LPS or IL-4. To test this, B cells were stimulated with either IL-4, LPS, or a combination of both, and changes in cholesterol were measured by filipin staining. Stimulation with either LPS or IL-4 for 24 hours was able to increase cholesterol levels, although LPS had a greater effect (Figure 10A).

**Figure 10.**
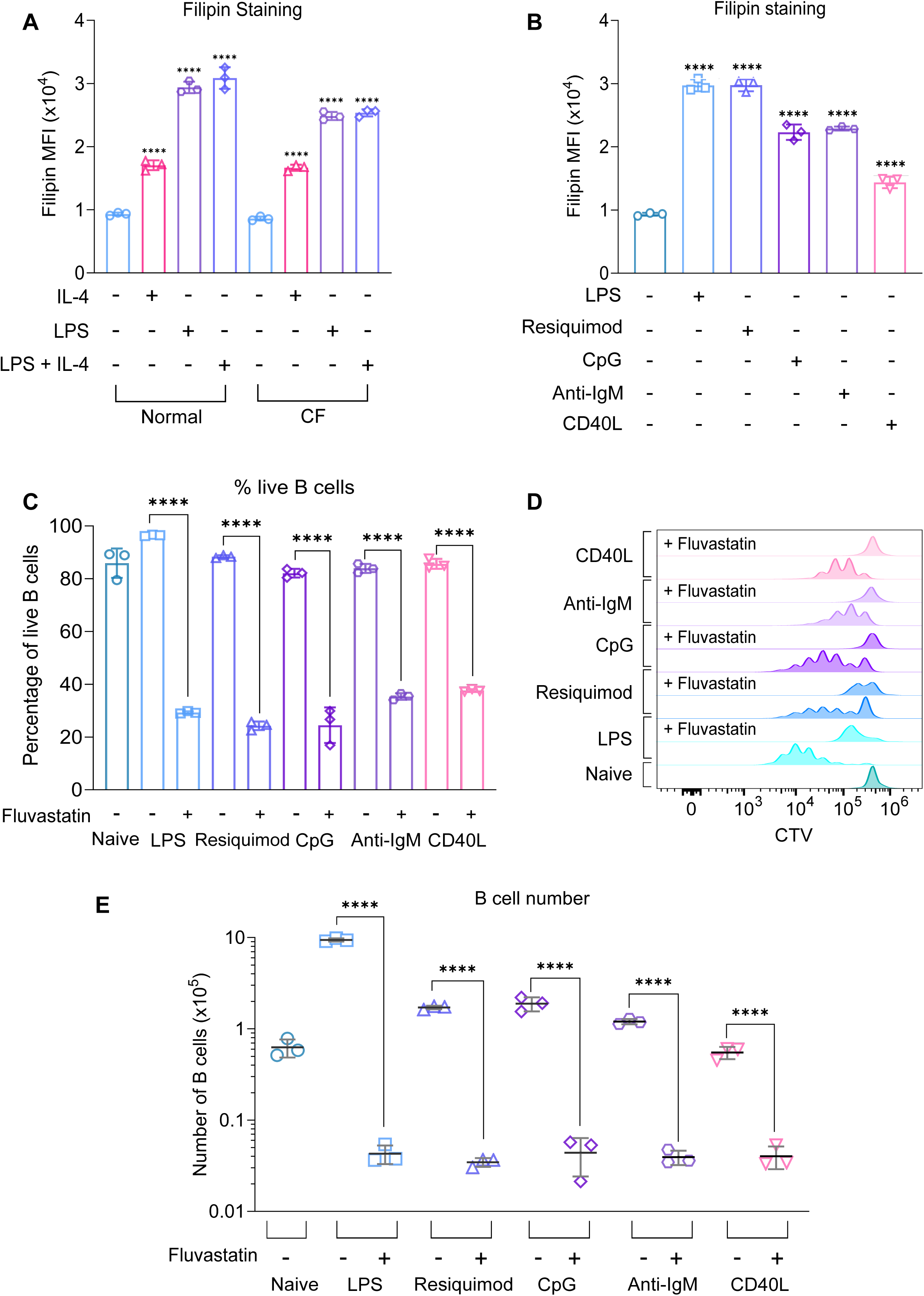
Cholesterol is required for the growth and proliferation of B cells by multiple stimuli. **(A)** Splenocytes from C57BL6/J mice were plated in normal or CF media, before stimulation with LPS (20μg/ml), IL-4 (10μg/ml) or a combination of LPS and IL-4. stimuli. The cells were fixed after 24 hours and stained with filipin. **(A)** Filipin staining comparing cholesterol content between different stimuli. Data shows the results of three biological replicates. Statistical power was determined using two-way ANOVA followed by multiple comparison testing via Tukey’s analysis. For comparisons to naïve B cells, *p*<0.0001 is indicated by ****. **(B)** Splenocytes from C57BL6/J mice were plated in normal media before stimulation with LPS (20μg/ml), Resiquimod (1μg/ml), ODN 1826 (1μg/ml), Anti-IgM (10μg/ml), or CD40L (500ng/ml). The cells were fixed after 24 hours and stained with filipin. **(B)** Filipin staining comparing cholesterol content between different stimuli. Data shows the results of three biological replicates. Statistical power was determined using one-way ANOVA followed by multiple comparison testing via Dunnett’s analysis. For comparisons to naïve B cells, *p*<0.0001 is indicated by ****. **(C-E)** B cells were purified from the spleens of C57BL6/J mice and cultured in normal media. The cells were stained with CTV, then pre-treated with DMSO as a vehicle control or Fluvastatin (10µM) for 45 minutes before stimulation with LPS (20μg/ml), Resiquimod (1μg/ml), ODN 1826 (1μg/ml), Anti-IgM (10μg/ml), or CD40L (500ng/ml) as indicated for 72 hours. **(C)** shows the percentage of live B cells (7AAD-ve), **(D)** representative histogram for CTV staining and **(E)** live B cell number. Data shows the results of three biological replicates. Statistical power was determined using two-way ANOVA followed by multiple comparison testing via Tukey’s analysis, where *p*<0.0001 is indicated by ****. For **(E),** this data was log-transformed and then statistically analysed due to unequal variance. Statistical power was determined using two-way ANOVA followed by multiple comparison testing via Sidak’s analysis, where *p*<0.0001 is indicated by ****.

Considering the effects of HMGCR inhibition on cholesterol levels following LPS stimulation, we also tested whether other stimuli known to promote B cell proliferation could elevate cholesterol levels. Both the TLR7 agonist Resiquimod and the TLR9 agonist CpG increased cholesterol levels in B cells (Figure 10B). Similar to TLR agonists, both anti-IgM and CD40L stimulation increased cholesterol levels in B cells (Figure 10B). Likewise, Resiquimod, CpG, anti-IgM, and CD40L were all able to stimulate proliferation, as evidenced by CTV labelling and live B cell number after 72 hours of treatment (Figure 10D, E). While none of these stimuli were as effective at inducing proliferation as LPS, in each case, treatment with Fluvastatin reduced survival and blocked proliferation of the B cells (Figure 10C-E).

Since stimulation with these agonists increased cholesterol levels in B cells, we wondered whether that correlated with an increased expression of key proteins involved in cholesterol metabolism at a proteomic level. Based on proteomic data from (James *et al*, 2024) stimulation of B cells with either IL-4, anti-IgM or anti-CD40 could increase levels of HMGCR, SQLE and LDLR, with the largest increase in expression seen with a combination of these stimuli (Figure S13A-D).

## Discussion

In this study, we have used proteomic analysis to highlight the switch in metabolic demand from naïve B cells in a resting state to LPS + IL-4 activated B cells, which significantly increase their energy and biosynthetic requirements to sustain growth, proliferation, and class switching.

Upon activation, B cells upregulated several amino acid transporters, including the system L transporters SLC7A5 and 6 (Figure 4A), which can import large neutral amino acids, including the essential amino acids histidine, isoleucine, leucine, methionine, phenylalanine, threonine, tryptophan, and valine (Fotiadis *et al*, 2013). Of these 2 transporters, SLC7A5 was expressed at a higher level and was responsible for the majority of amino acid uptake via system 1 transporters (Figure 4J). SLC7A5 forms a dimer with CD98 (SLC3A2) to form an active transporter. SLC7A5 is not, however, the only binding partner of CD98, as it can also bind to the SLC7A11 and SLC7A6 transporters or interact with integrins (Kahya *et al*, 2021). Previous work has shown that mice with B cell-specific deletions of CD98 presented normal development of B cells in the bone marrow, but significantly lower levels of class-switched IgG due to the inhibition of B cell proliferation and plasma cell formation following the activation of naive B cells in the periphery (Cantor *et al*, 2009). The defect in proliferation was attributed to defective integrin signalling rather than an impairment in amino acid transport, based on the deletion of the integrin or amino acid transporter regions in CD98. Knockout of SLC7A5 in either hematopoietic cells or B cells has been previously shown not to affect the number of splenic B cells in mice (Sinclair *et al*, 2013), although a separate study found a decrease in the number of peritoneal B1 cells (Sun *et al*, 2023). While not critical for initial B cell development, SLC7A5 was critical for the survival and expansion of B cells in response to *ex vivo* stimulation with LPS + IL-4 (Figure 4F-I). Of note, recent research in human B cells has shown that stimulation with CpG increases SLC7A5 expression, which in turn transports L-leucine into the cell and facilitates mTORC activation, which was required for IgG production and inflammatory cytokine secretion (Torigoe *et al*, 2019).

Similar observations have been made in T cells, where SLC7A5 is not required for initial T cell development but is required for the clonal expansion and effector functions of T cells following activation via the T cell receptor in the periphery (Sinclair *et al*, 2013). This suggests that B cells, similar to T cells, may rely on different amino acid transporters at different points in their development. The expression of SLC7A5 following T cell activation has been shown to require the transcription factor c-Myc (Marchingo *et al*, 2020). While the role of c-Myc was not directly addressed in this study, analysis of the proteomic data presented here does show that c-Myc was increased by LPS stimulation (Table S3). This is consistent with previous reports showing an upregulation of c-Myc in response to several B cell activating stimuli, including LPS, CD40, and the BCR (Luo *et al*, 2018; Dominguez-Sola *et al*, 2012; Calado *et al*, 2012; Tesi *et al*, 2019). While in T cells, c-Myc regulates metabolic changes, including the expression of SLC7A5 (Sinclair & Cantrell, 2025), this has not been studied to the same degree in B cells. Of note, transcriptional profiling following LPS stimulation of B cells has suggested that SLCA5 mRNA induction in response to LPS can occur in the absence of c-Myc (Tesi *et al*, 2019), suggesting that the regulation of SLC7A5 may not be identical between T and B cells.

As B cells expand and divide, they will require an increase in lipids to generate new membrane. Cholesterol is an important lipid component of the plasma membrane, which affects membrane fluidity (Luo *et al*, 2020) and has also been implicated in the regulation of several receptors, including the BCR (Bléry *et al*, 2006). The regulation of cholesterol levels in cells is therefore closely controlled at the level of both cholesterol biosynthesis and its uptake via the endocytosis of LDL particles. Low levels of cholesterol in the endoplasmic reticulum are detected by the transcription factor SREBP2 (Luo *et al*, 2020; Brown *et al*, 2018). The importance of this system in B cells was recently demonstrated by the conditional knockout of SCAP in B cells, a key regulator of SREBP1 and 2. While this did not prevent the development of B cells or their levels at steady state in mice, it did prevent the activation and proliferation of B cells in response to antigens and thus compromised the humoral immune response. SCAP deletion will, however, affect both SREBP1 and SREBP2 function and therefore have wider effects on lipid metabolism beyond its effects on cholesterol. We show here that in response to several activating stimuli *ex vivo*, including LPS, B cells upregulate cholesterol, and that blocking HMGCR, a key rate-limiting enzyme in the cholesterol biosynthesis pathway, inhibits B cell proliferation (Figure 10B-E). In the case of LPS stimulation, B cells seem to be able to obtain cholesterol by both uptake and biosynthesis. For example, B cells can still increase their cholesterol levels in the absence of LDL in the cell culture medium (Figure 5I). In addition, NB-598, an inhibitor of the 2nd rate-limiting enzyme in the cholesterol biosynthesis pathway, can reduce the ability of LPS to increase cholesterol levels (Figure 5I). In contrast, the greater effect of NB-598 on cholesterol levels of B cells grown in CF media relative to normal media (Figure 5I), and the inability of mevalonate to rescue the effects of statin treatment in CF media (Figure 7B-F), would indicate that B cells also use LDL uptake as a source of cholesterol. In line with this, B cells upregulated the LDLR following activation (Figure 5D). Additionally, proteomic analysis of LPS + IL-4 stimulated B cells grown in normal vs CF media, or +/- Fluvastatin treatment (normal) reveals an upregulation of proteins involved in cholesterol metabolism, including HMGCR, SQLE and LDLR (Figure S9). This suggests that a feedback mechanism exists whereby the detection of low cholesterol levels (either due to blocking biosynthesis or uptake) by SREBP2 increases the expression of cholesterol metabolic proteins. A similar study in activated CD8+ T cells has demonstrated that knockout of the LDLR leads to an upregulation of proteins involved in cholesterol biosynthesis, including HMGCR, and a reduction in proteins involved in cholesterol efflux compared to WT CD8+ T cells (Bonacina *et al*, 2022). The precise balance between these two mechanisms in B cells may depend on the duration of stimulation and the availability of LDL in the extracellular environment.

In line with our results, following CD40 mediated activation of human B cells, Atorvastatin was reported to inhibit human B cell proliferation and the expression of the activation markers CD80 and CD86 (Shimabukuro-Vornhagen *et al*, 2014). Interpretation of the effects of statins on B cells is, however, complex. Statins inhibit HMGCR, the rate-limiting enzyme in the mevalonate pathway that is required for the initial steps in cholesterol biosynthesis. This pathway generates farnesyl pyrophosphate, an intermediate that can feed into not only cholesterol biosynthesis, but also protein prenylation and ubiquinone pathways (Mullen *et al*, 2016; Wang & Casey, 2016). Thus, the effects of statins reported here could be due not only to their effects on cholesterol, but also via these other pathways. Prenylation has been linked to proliferation in several cell types (Su *et al*, 2020), including T cells, where Simvastatin inhibits activation via two mechanisms, one of which involves blocking the prenylation of Ras and Rac (Ghittoni *et al*, 2005). We therefore investigated whether statin-mediated effects on B cell survival and proliferation were in part due to prenylation. We found that FGTI-2734, a dual inhibitor of FTase and GGTase, blocked LPS + IL-4 induced B cell proliferation and survival (Figure 6E-H). Similar GGTI-298, an inhibitor of GGTase, had a stronger inhibitory effect on B cell function compared to FTI-277, which blocks FTase activity, suggesting an important role for prenylation in B cells (Figure 6I-P). In agreement with this, prenylation has also been reported to be important in B cells for the expression of CD80 and CD86 following stimulation via CD40, although its role in proliferation was not directly addressed (Shimabukuro-Vornhagen *et al*, 2014). Given that prenylation can target a large number of proteins, including many members of the Ras GTPase superfamily, to regulate their cellular localisation, the effects of blocking prenylation in B cells are likely to be complex. Interestingly, GGPP, which is needed for protein geranylgeranylation, was able to rescue the effects of Fluvastatin on LPS + IL-4 stimulated B cells in normal media

(Figure 8A-D). Notably, it was more efficient than mevalonate at restoring proliferation in the Fluvastatin treated B cells. This ability to rescue proliferation was unexpected, as GGPP would not directly feed into the cholesterol biosynthesis pathway, and other experiments described here using CF media and squalene monooxygenase inhibitors argue for a requirement of cholesterol for B cell proliferation. We therefore looked at its effect on cholesterol levels and found that GGPP was able to rescue the effects of Fluvastatin in normal but not CF media (Figure 8I,J). This could be explained by prenylation being required for cholesterol uptake but not biosynthesis. Cholesterol uptake via the LDLR requires the clathrin-mediated endocytosis of the LDLR and its trafficking to endosomes, where cholesterol is released from LDL (Nguyen *et al*, 2025). Vesicle trafficking is regulated via the action of several Rab GTPases, including Rab5, which is recruited to early endosomes and regulates their maturation (Zhao *et al*, 2025; Zeigerer *et al*, 2012). More significantly, Rab5 prenylation has been shown to be important for its correct targeting to endosomes, and therefore its biological function (Gomes *et al*, 2003). Treatment of cells with statins to inhibit prenylation has previously been shown to affect Rab subcellular localisation and vesicle trafficking (Ali *et al*, 2010; Ronzier *et al*, 2019; Xia *et al*, 2018). Loss of the prenylation of Rab proteins involved in LDLR endocytosis could therefore negatively impact this process, leading to a failure of LDL uptake into the B cell. Prenylation is also likely to impact other processes in B cells, for example, TLR9-induced IL-10 production by regulatory B cells has been found to require geranylgeranylation and not farnesylation (Bibby *et al*, 2020). Further work will be necessary to identify the proteins prenylated upon B cell activation and to determine the effects of prenylation on their function.

In terms of the pathways that control B cell function downstream of TLR4 signalling, we found that inhibition of p38α/β and MEK1/2 signalling impacted B cell growth and proliferation, as well as cholesterol levels (Figure 9A-F). Related to this, inhibition of MEK1/2 has been found to abolish *LDLR* promoter activity (Kotzka *et al*, 2000), and depletion of ERK2, but not ERK1, significantly reduced the expression levels of *SREBP2*, *HMGCR* and *LDLR* (Yu *et al*, 2019). Studies in primary mouse hepatocytes have demonstrated that treatment with the p38α/β inhibitor SB203580 also significantly reduced *LDLR* promoter activity (Pham *et al*, 2016).

Additionally, we found that blocking mTOR had a major inhibitory effect on B cell proliferation (Figure 9A-B), as well as reducing cholesterol levels (Figure 9F), and to a lesser extent, the rate of protein synthesis and amino acid uptake through System L transporters (Figure 12A-B). In response to environmental cues, including energy and nutrient availability, the mTORC1 complex regulates processes such as cell growth and proliferation (Saxton & Sabatini, 2017). Previous studies have shown that the addition of amino acids to the growth medium increases the transcription of genes involved in cholesterol biosynthesis, including HMGCR and SREBP2 (Eid *et al*, 2017). Treatment with the mTORC1 inhibitor Torin1 blocked this effect, suggesting that cholesterol metabolism may be dependent on mTORC1 activity (Eid *et al*, 2017).

Overall, the proteomic analysis described here supports that B cells undergo a major reorganisation of metabolic pathways following TLR activation. Naïve B cells exist in a quiescent state where they do not grow or proliferate and express low levels of metabolic enzymes. Conversely, B cells stimulated with LPS + IL-4 massively upregulate metabolic enzymes involved in processes including protein synthesis, amino acid uptake, and cholesterol metabolism, which is required to support an increase in cell size, proliferation, and antibody class switching. Most significantly, we found that LPS + IL-4 stimulated B cells depended on cholesterol biosynthesis, cholesterol uptake, and prenylation for expansion and proliferation.

## Methods

### Mice

Wildtype C57BL/6 mice (Charles Rivers) aged between 8 and 12 weeks were used for the majority of experiments. Conditional *SLC7A5* knockout mice bred to Vav-iCre mice have been described previously and were maintained on a C57Bl6/J background (de Boer *et al*, 2003; Poncet *et al*, 2014; Sinclair *et al*, 2013). MyD88 knockout mice have been described previously and were maintained on a C57Bl6/J background (Adachi et al, 1998). The mice were kept in individually ventilated cages under specific pathogen-free conditions. Mice were given free access to food (RM3 irradiated pelleted diet, Special Diet Services) and water. Animal rooms were maintained at 21°C and a humidity of 45 to 55% with a 12/12 h light cycle. Mice were sacrificed using a rising concentration of CO2, and death was confirmed by either cervical dislocation or exsanguination. All animal work was approved by the University of Dundee’s Welfare and Ethical Use of Animals Committee and in accordance with UK and EU law.

### Cell isolation

Lymph nodes were homogenised through a 40µM cell strainer in DPBS buffer (Gibco), and after centrifugation at 450g for 4 minutes, the pellet was resuspended in DPBS. Lymphocytes were counted on a Novocyte (Agilent Technologies) and seeded at 1×10^6^ cells/ml in B cell medium (RPMI-1640 supplemented with 10% FBS, 50 U/ml penicillin-streptomycin, 5mM L-glutamine, 10mM HEPES buffer, 1mM sodium pyruvate, 50 µM 2-mercaptoethanol and non-essential amino acids) for stimulation.

Spleens were dissociated through a 40µM filter in DPBS buffer, and after centrifugation at 450g for 4 minutes, the pellet was resuspended in 1ml of Red Blood Cell Lysing Buffer (Hybri-Max™, Sigma) for 4 minutes at room temperature to lyse red blood cells. The reaction was quenched by the addition of DPBS, and the cells were pelleted by centrifugation at 450g for 4 minutes. The pellet was resuspended in 10ml DPBS before counting the splenocytes on a Novocyte. Splenocytes were seeded at 1×10^6^ cells/ml in B cell medium for stimulation or used for further purification of B cells.

### B cell isolation

To obtain an enriched population of B cells, magnetic sorting was used to remove non-B cells. Briefly, splenocytes were incubated with Fc receptor-blocking anti-CD16/CD32 antibody (BD Pharmingen) for 5 minutes on ice. The cells were incubated with a mixture of biotin-labelled anti-mouse antibodies (10µg/ml) (Table 1) for 20 minutes on ice. Unbound antibody was removed by washing with 10ml Magnetic Activated Cell Sorting (MACS) buffer (DPBS + 0.5% BSA + 2mM EDTA), followed by centrifugation. The cells were then resuspended in MACS buffer with streptavidin microbeads (1:10) (Miltenyi Biotech) and incubated at 4⁰C for 15 minutes. Miltenyi LD columns were calibrated by passing through 3ml of MACS buffer. Cells were washed with 10ml MACS buffer and centrifuged at 450g for 4 minutes to remove excess beads. The cell pellet was then resuspended in MACS buffer and separated using magnetic-activated cell sorting columns (LD) according to the manufacturer’s instructions (Miltenyi Biotec). Single-cell suspensions were collected, and B cell numbers and purity (CD19+) were measured on a NovoCyte. Typical purity levels were 97% CD19+ B cells. B cells were seeded at 2×10⁵ cells/ml of B cell medium for stimulation for flow cytometry experiments unless otherwise stated. When stated, purified B cells were stained with 2.5µM CellTrace™ Violet (ThermoFisher) at 37⁰C for 20 minutes, then quenched in 10ml B cell medium before seeding at the required density.

**Table 1:** MACS antibodies.

| Table 1: MACS antibodies |  |  |  |  |
| --- | --- | --- | --- | --- |
| Antibody | Clone | Concentration | Catalogue number | Supplier |
| Anti-CD11b | M1/70 | 10µg/ml | 101204 | BioLegend |
| Anti-CD11c | N418 | 10µg/ml | 117304 | BioLegend |
| Anti-CD3ε | 145-2C11 | 10µg/ml | 100304 | BioLegend |
| Anti-TER119 | TER-119 | 10µg/ml | 116204 | BioLegend |

### Stimulations and inhibitors

Cells were rested at 37⁰C for 1 hour. When required, the cells were pre-treated with the appropriate inhibitor (Table 2) for 45 minutes before stimulation, at the concentrations indicated in the figure legend. Inhibitors were dissolved in DMSO (Thermo Fisher) and used at a dilution of 1:1000 of culture medium. As indicated in the figure legends, cells were treated with 1μl DMSO as a vehicle control.

**Table 2:** Inhibitors.

| Table 2: Inhibitors |  |  |  |  |
| --- | --- | --- | --- | --- |
| Name | Target | Concentration | Catalogue number | Supplier |
| Fluvastatin sodium | HMG-CoA reductase | 0.1µM -100µM | 3309 | Tocris |
| Rosuvastatin calcium | HMG-CoA reductase | 0.1µM -100µM | 6343 | Tocris |
| NB-598 | Squalene monooxygenase | 0.1µM -100µM | HY-16343 | MedChem Express |
| FGTI-2734 | Farnesyl and geranylgeranyl transferase-1 | 10µM | HY-128350 | MedChem Express |
| FTI-277 hydrochloride | Farnesyl transferase | 10µM-30µM | HY-15872A | MedChem Express |
| GGTI-289 trifluoroacetate | Geranylgeranyl transferase-1 | 10µM-30µM | HY-15871 | MedChem Express |
| PD18352 | MSK1/2 | 2µM | 4237 | Tocris |
| VX745 | P38 | 1µM | 3915 | Tocris |
| Rapamycin | mTOR | 20nM | R8781 | Merck |

Cells were pre-incubated with the following metabolites at 37⁰C for 1 hour before inhibitor treatment/stimulation. Metabolites were used at the concentrations indicated in the figure legend: Mevalonic acid 5-phosphate lithium salt hydrate (Sigma Aldrich), which was dissolved in HEPES (Thermo Fisher). As indicated in the figure legends, cells were treated with 20µl HEPES as a vehicle control. Geranylgeranyl pyrophosphate ammonium salt (Sigma Aldrich), which comes in a 7:3 methanol: ammonium hydroxide solution. As indicated in the figure legends, cells were treated with 4.55µl 7:3 methanol: ammonium hydroxide solution as a vehicle control.

A combination of 20µg/ml LPS (L2654, Sigma) and 10ng/ml IL-4 (214-14, Peprotech) was used to stimulate the cells unless otherwise stated in the figure legend. For some experiments, cells were treated with LPS or IL-4 alone, or with 1µg/ml Resiquimod/R848 (tlrl-r848, Invivogen), 1µg/ml ODN 2006 (tlrl-2006, Invivogen), 10µg/ml Anti-Mouse IgM μ chain (115-006-075, Stratech) or 500ng/ml CD40 ligand (8230-CL-050/CF, R&D).

### Cholesterol free serum

Silica-depleted FBS was purchased from Biowest and used in place of normal FBS when making B cell culture medium. Samples of normal and cholesterol-free FBS were sent to London Health Company, which carried out tests to measure changes in triglycerides, cholesterol, high-density lipoprotein (HDL) and low-density lipoprotein (LDL) (Table S2).

### Cell survival, proliferation and class switching assays

Following stimulation, cells were washed in Fluorescence Activated Cell Sorting (FACS) buffer (1% BSA, DPBS) and then incubated with Fc receptor-blocking anti-CD16/CD32 antibody in FACS buffer for 15 minutes at 4⁰C before staining with the appropriate fluorochrome-conjugated antibodies (Table 3) in FACS buffer for 20 minutes at 4⁰C. The samples were washed with FACS buffer prior to resuspension in 0.25μg of 7AAD. Flow cytometry data was collected on a Novocyte and analysed using FlowJo V.10 software (BD Biosciences). A representative gating strategy is shown in Figure S14A for lymph node/splenocyte isolation, and Figure S15B for B cell isolation.

**Table 3:**
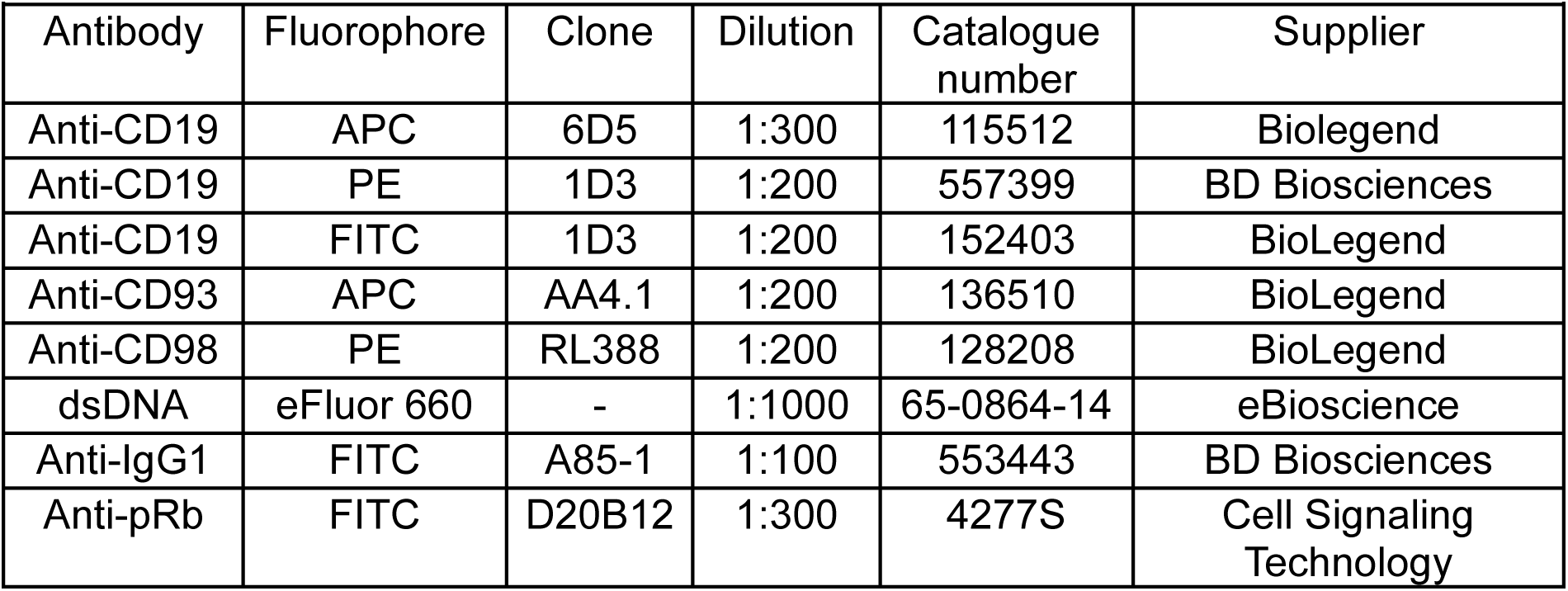
Flow cytometry antibodies.

| Table 3: Flow cytometry antibodies |  |  |  |  |  |
| --- | --- | --- | --- | --- | --- |
| Antibody | Fluorophore | Clone | Dilution | Catalogue number | Supplier |
| Anti-CD19 | APC | 6D5 | 1:300 | 115512 | Biolegend |
| Anti-CD19 | PE | 1D3 | 1:200 | 557399 | BD Biosciences |
| Anti-CD19 | FITC | 1D3 | 1:200 | 152403 | BioLegend |
| Anti-CD93 | APC | AA4.1 | 1:200 | 136510 | BioLegend |
| Anti-CD98 | PE | RL388 | 1:200 | 128208 | BioLegend |
| dsDNA | eFluor 660 | - | 1:1000 | 65-0864-14 | eBioscience |
| Anti-IgG1 | FITC | A85-1 | 1:100 | 553443 | BD Biosciences |
| Anti-pRb | FITC | D20B12 | 1:300 | 4277S | Cell Signaling Technology |

### p-STAT1 staining

Splenocytes (1×10^6^ cells/ml) were stimulated with LPS (20μg/ml) and/or IFN-β for 15 minutes before washing in FACS buffer. The cells were fixed using 1:1 IC fixation buffer (catalogue number: 00-8222-49, eBiosciences) in FACS buffer and incubated at room temperature for 20 minutes. Following another wash in FACS buffer, cells were resuspended in 100µl of cold ethanol and stored at -20°C for 30 minutes. The samples were washed with FACS buffer twice before incubation with Fc receptor-blocking anti-CD16/CD32 antibody in FACS buffer for 15 minutes at 4⁰C. The cells were incubated with p-STAT1 (Table 4) for 1 hour at room temperature before washing with FACS buffer and incubation with (1:1000) anti-rabbit IgG (H+L), F(ab’)2 Fragment (Alexa Fluor^®^ 647 Conjugate) (Cell Signalling Technology, 4414S) and CD19 PE (Table 3) for 1 hour at room temperature. The cells were washed with FACS buffer before resuspension in 300μl FACS buffer. Flow cytometry data was collected on a Novocyte and analysed using FlowJo V.10 software.

**Table 4:**
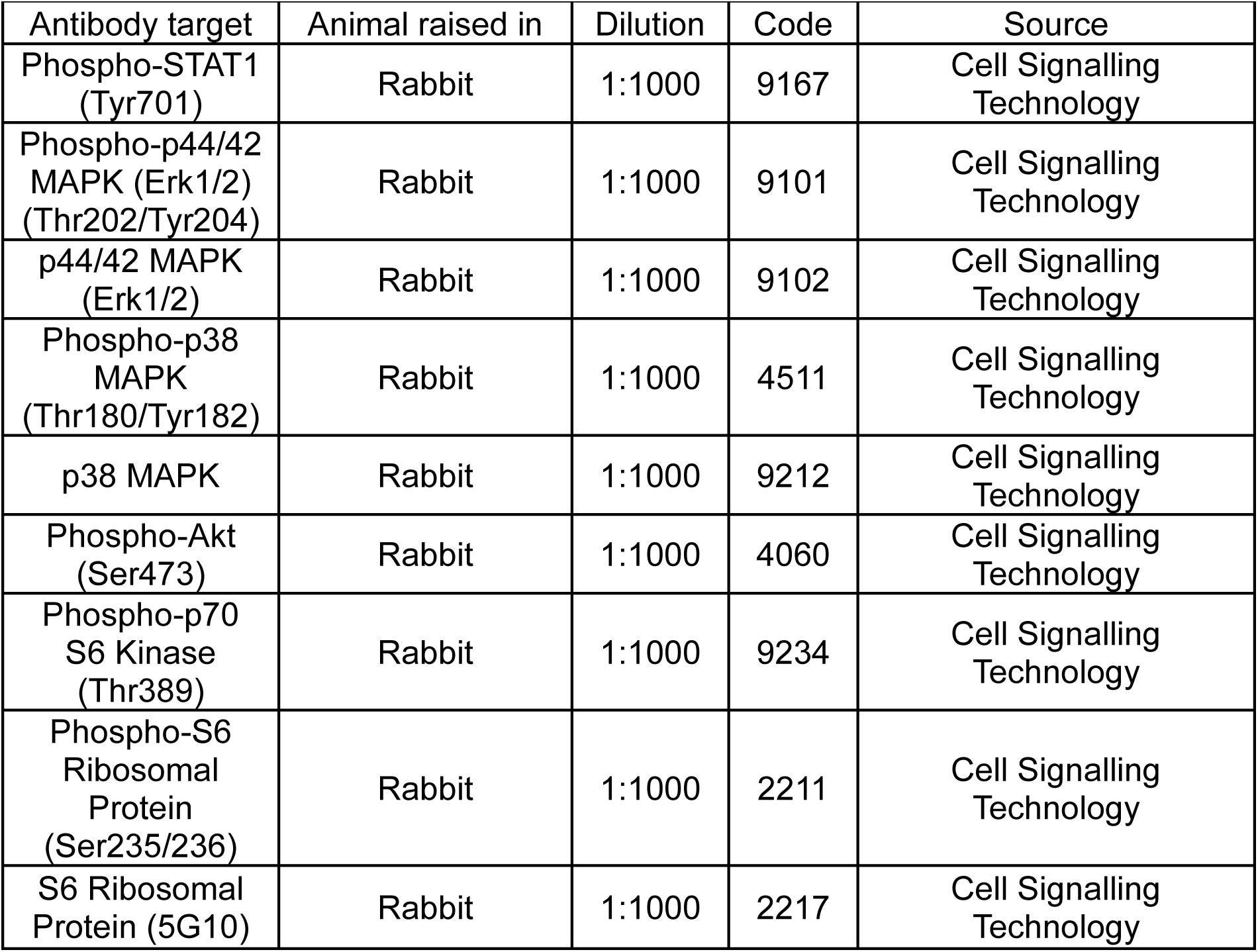
Western blot primary antibodies.

| Table 4: Western blot primary antibodies |  |  |  |  |
| --- | --- | --- | --- | --- |
| Antibody target | Animal raised in | Dilution | Code | Source |
| Phospho-STAT1 (Tyr701) | Rabbit | 1:1000 | 9167 | Cell Signalling Technology |
| Phospho-p44/42 MAPK (Erk1/2) (Thr202/Tyr204) | Rabbit | 1:1000 | 9101 | Cell Signalling Technology |
| p44/42 MAPK (Erk1/2) | Rabbit | 1:1000 | 9102 | Cell Signalling Technology |
| Phospho-p38 MAPK (Thr180/Tyr182) | Rabbit | 1:1000 | 4511 | Cell Signalling Technology |
| p38 MAPK | Rabbit | 1:1000 | 9212 | Cell Signalling Technology |
| Phospho-Akt (Ser473) | Rabbit | 1:1000 | 4060 | Cell Signalling Technology |
| Phospho-p70 S6 Kinase (Thr389) | Rabbit | 1:1000 | 9234 | Cell Signalling Technology |
| Phospho-S6 Ribosomal Protein (Ser235/236) | Rabbit | 1:1000 | 2211 | Cell Signalling Technology |
| S6 Ribosomal Protein (5G10) | Rabbit | 1:1000 | 2217 | Cell Signalling Technology |

### Cell cycle analysis and p-Rb staining

Cells were washed in FACS buffer and then fixed using 1:1 IC fixation buffer and FACS buffer, and incubated at room temperature for 20 minutes. Following another wash in FACS buffer, cells were resuspended in 100µl of cold ethanol and stored at -20°C for 30 minutes. The samples were washed with FACS buffer twice before incubation with Fc receptor-blocking anti-CD16/CD32 antibody in FACS buffer for 15 minutes at 4⁰C. Surface stains (Table 3) were incubated for 1 hour at room temperature before washing with FACS buffer and resuspension in 0.5μg/ml of DAPI. Flow cytometry data was collected on a Novocyte and analysed using FlowJo V.10 software. A representative gating strategy is shown in Figure S14B.

### O-propargyl-puromycin uptake assay

Uptake of the puromycin analogue O-propargyl-puromycin was used to estimate protein synthesis rates. Splenocytes (1×10^5^ cells/100µl) were stimulated with 20μg/ml LPS and 10ng/ml IL-4. After 24 hours, the cells were treated with 20µM of O-propargyl-puromycin for 30 minutes at 37°C. 100µl of 2% paraformaldehyde in B cell medium was added to the cells before incubation in complete darkness at room temperature for 30 minutes. The samples were pelleted by centrifugation at 450g for 4 minutes before resuspension in 100µl of DPBS. Following this, the samples were washed with FACS buffer and then resuspended in 100µl of 0.01% Saponin (Sigma) in DPBS, before incubation at room temperature for 20 minutes. The samples were pelleted by centrifugation at 450g for 4 minutes before resuspension in 30µl of Click mixture: 1mM Copper sulphate (209198, Sigma), 10mM Sodium ascorbate (A7631. Sigma), 1mM BTTAA (906328, Sigma), 10mM Aminoguanidine (81530, Cayman Chemicals), DPBS (14190144, Gibco), 5µM Alexafluor 647 azide (A10277, Invitrogen) and incubation in complete darkness at room temperature for 1 hour. The samples were pelleted by centrifugation at 450g for 4 minutes before resuspension in 200µl of FACS buffer and incubation in complete darkness at room temperature for 30 minutes. Again, the samples were pelleted by centrifugation at 450g for 4 minutes before washing again in FACS buffer. The cells were incubated with Fc receptor-blocking anti-CD16/CD32 antibody in FACS buffer for 15 minutes at 4⁰C. Surface stains (Table 3) were incubated for 1 hour at room temperature. The samples were washed with FACS buffer prior to resuspension in 300µl FACS buffer. Flow cytometry data was collected on a Novocyte and analysed using FlowJo V.10 software. A representative gating strategy is shown in Figure S14C.

### Kynurenine uptake assay

Uptake of the tryptophan metabolite kynurenine was used to estimate the uptake rates via neutral amino acid transporters (Sinclair *et al*, 2018). Purified B cells (1×10^6^ cells/ml) were stimulated with 20μg/ml LPS and 10ng/ml IL-4. For the assay, pre-warm HBSS, BCH (40mM in HBSS) and Kynurenine (800μM in HBSS) to 37°C. After 24 hours, the cells were washed in FACS buffer and then incubated with Fc receptor-blocking anti-CD16/CD32 antibody in FACS buffer for 15 minutes at 37°C before staining with CD19 APC (Table 3) in FACS buffer for 20 minutes at 37°C. The samples were washed with FACS buffer and then resuspended in 200μl HBSS. Keep the samples at 37°C. Either 100μl of HBSS or 100μl of BCH was then added to each sample. After this, 100μl of Kynurenine was added, and the samples were incubated for 5 minutes at 37°C. After 5 minutes, the samples were fixed by adding 125μl of 4% paraformaldehyde made in DPBS for 30 minutes at room temperature. The samples were then washed 2 times with FACS buffer and re-suspended in 400μl of FACS buffer. Flow cytometry data was collected on an LSR Fortessa (BD biosciences) and analysed using FlowJo V.10 software. A representative gating strategy is shown in Figure S14D.

### Cholesterol staining assay

Changes in cellular cholesterol content were determined using the cholesterol-binding agent filipin (Miller, 1984). Splenocytes were stimulated with 20μg/ml LPS and 10ng/ml IL-4. After 24 hours, the cells were washed twice in DPBS before staining for viability using eFluor 660 fixable viability dye (Table 3) for 30 minutes at 4⁰C. After two washes in FACS buffer, the cells were fixed using 1:1 IC fixation buffer and FACS buffer, before incubation at 4⁰C for 20 minutes. Following another wash in FACS buffer, cells were incubated with Fc receptor-blocking anti-CD16/CD32 antibody in FACS buffer for 15 minutes at 4⁰C. Surface stains (Table 3) were incubated for 20 minutes at 4⁰C. The samples were washed with FACS buffer prior to resuspension in 270µl FACS buffer and 30µl of 1µg/µl filipin (Merck) before incubation in complete darkness at room temperature for 30 minutes. Flow cytometry data was collected on an LSR Fortessa and analysed using FlowJo V.10 software. A representative gating strategy is shown in Figure S15A.

### Mitochondria staining assay

Splenocytes were stimulated with 20μg/ml LPS and 10ng/ml IL-4. After 24 hours, the cells were washed twice in FACS buffer. 50µl of prewarmed (37°C) staining solution containing 100nM MitoTracker Red probe (Invitrogen) in FACS buffer was added to the cells before incubation at 37°C for 20 minutes. The samples were washed with FACS buffer before incubation with Fc receptor-blocking anti-CD16/CD32 antibody (BD Pharmingen) in FACS buffer for 15 minutes at 4⁰C before staining with the appropriate fluorochrome-conjugated antibodies (Table 3) in FACS buffer for 20 minutes at 4⁰C. The samples were washed with FACS buffer prior to resuspension in 0.25μg of 7AAD. Flow cytometry data was collected on a Novocyte and analysed using FlowJo V.10 software. A representative gating strategy is shown in Figure S15C.

### Immunoblotting

B cells were seeded at 2×10^6^ cells/ml of B cell medium for immunoblotting. To generate lysates, the cells were stimulated with LPS (20µg/ml) for 60 minutes and centrifuged at 100g for 2 minutes. The pellet was resuspended in 300µl of SDS Triton Lysis buffer (50mM Tris-HCL pH 7.5, 1% (v/v) Triton-X-100, 1mM EGTA, 1mM EDTA, 1mM Sodium Orthovanadate, 50mM Sodium fluoride, 1mM Sodium Pyrophosphate, 10mM Sodium B-glycerophosphate, 0.27M Sucrose, 0.1% (v/v) 2-β-mercatoethanol, cOmplete mini EDTA-Free Protease Inhibitor, 1% Sodium Dodecyl sulfate, 5% Glycerol) and incubated for 5 minutes at 100⁰C to denature proteins. Following a brief period of cooling, the DNA was sheared using a 25g syringe and the samples were stored at -20⁰C. Cell lysates were resolved by SDS-PAGE using 10% gels in running buffer (250mM Tris, 1% (w/v) 10% Sodium Dodecyl Sulfate, 192mM glycine) and transferred to nitrocellulose membranes in the presence of transfer buffer (48 mM Tris, 39 mM glycine, 20% (v/v) methanol). The membranes were blocked using 5% (w/v) skimmed milk powder (Marvel) in TBST (0.5M Tris-HCl, pH 7.6, 1.5M NaCl and 0.1%(v/v) Tween 20). Primary antibodies (Table 4) were used at a dilution of 1/1000 overnight in TBST with 5% (w/v) BSA. The membranes were incubated with Anti-rabbit HRP (horseradish peroxidase) secondary antibodies (Sigma) in 5% (w/v) milk powder in TBST. The blots were developed with Clarity™ Western ECL substrate (Bio-Rad) and scanned using Li-Cor Odyssey Fc Imager (Licor) (Chemi, 600nm and 700nm channels). Image Studio software was utilised to compile Western blot data.

### Proteomics

For the naïve vs LPS + IL-4 dataset, wildtype C57BL/6 mice (Charles Rivers) aged between 8 and 12 weeks were used for all proteomic experiments. Lymph nodes were extracted from mice, mashed in RPMI media before filtering through a 70μm cell strainer. Naïve B cells were purified by fluorescence-activated cell sorting. Lymphocytes were incubated with Fc receptor-blocking anti-CD16/CD32 antibody (BD Pharmingen) in FACS buffer for 15 minutes at 4⁰C before staining with fluorochrome-conjugated antibodies (Table 3) in FACS buffer for 20 minutes at 4⁰C. The samples were washed with FACS buffer prior to resuspension in 0.5μg/ml of DAPI. CD19+, CD93-, DAPI- cells were sorted using a Sony SH800 cell sorter. Sorted cells were washed twice with HBSS and snap frozen in liquid nitrogen to be stored at -80 °C until processing for mass spectrometry. For B cell activation, lymphocytes were suspended at a final density of 1.5 million cells/ml in medium (RPMI-1640 containing glutamine supplemented with 10% FBS, 50 U/ml penicillin-streptomycin, 50 µM 2-mercaptoethanol) and activated for 24 hours in the presence of 20µg/ml LPS and 10ng/ml IL-4. After 24 hours, the cells were incubated with Fc receptor-blocking anti-CD16/CD32 antibody (BD Pharmingen) in FACS buffer for 15 minutes at 4⁰C before staining with fluorochrome-conjugated antibodies (Table 3) in FACS buffer for 20 minutes at 4⁰C. The samples were washed with FACS buffer prior to resuspension in 0.5μg/ml of DAPI. CD19+ and DAPI- were sorted and collected as described above. Cells were then washed 3 times in PBS and cell pellets frozen at -80°C.

For the dataset in Figure S9, B cells were isolated from murine spleens as described above and seeded at 2×10⁵ cells/ml of B cell medium, with 1×10^6^ cells/5ml per condition. Cells were rested at 37⁰C for 1 hour. Cells were washed twice with PBS and snap frozen in liquid nitrogen to be stored at -80°C until processing for mass spectrometry. For activation, B cells were pre-treated with Fluvastatin (10µM) for 45 minutes before stimulation. The cells were stimulated with LPS (20µg/ml) and IL-4 (10ng/ml) for 24 hours. Cells were washed twice with PBS and snap frozen in liquid nitrogen to be stored at -80 °C until processing for mass spectrometry.

Cell pellets were lysed, and proteins were digested following the protocol described by (Baker *et al*, 2022). In brief, 400 ml of lysis buffer (5% sodium dodecyl sulfate, 50 mM triethylammonium bicarbonate (pH 8.5) and 10 mM tris(2-carboxyethyl) phosphine-hydrochloride) was added to each sample and the lysates were shaken at room temperature at 1000 rpm for 5 minutes before being incubated at 95 °C at 500 rpm for 5 minutes. Samples were allowed to cool and were then sonicated using a BioRuptor (15 cycles: 30 sec on and 30 sec off) and alkylated with 20mM iodoacetamide for 1 h at 22 °C in the dark. To determine protein concentration, the EZQ protein quantitation kit (Thermo) was used, and protein cleanup and digestion were performed using S-TRAP mini columns (Protifi). Proteins were digested with trypsin at 1:20 ratio (enzyme:protein) for 2 hours at 47°C. Digested peptides were eluted from S-TRAP columns using 50mM ammonium bicarbonate, followed by 0.2% aqueous formic acid and 50% aqueous acetonitrile containing 0.2% formic acid. Peptides were dried by speedvac before resuspending in 1% formic acid. Peptide quantity was measured using the CBQCA kit (Thermo).

### LC-MS/MS analysis

#### Naïve vs LPS + IL-4 dataset

Peptides were analysed by single-shot Data Independent Acquisition (DIA) mass spectrometry as previously described (Molina-Gonzalez *et al*, 2023; Sollberger *et al*, 2024). 1.5 µg of peptide from each sample was injected onto a nanoscale C18 reverse-phase chromatography system (UltiMate 3000 RSLC nano, Thermo Scientific) and electrosprayed into an Orbitrap Exploris 480 Mass Spectrometer (Thermo Fisher). The following buffers were used for liquid chromatography: buffer A (0.1% formic acid in Milli-Q water (v/v)) and buffer B (80% acetonitrile and 0.1% formic acid in Milli-Q water (v/v). Samples were loaded at 10μl/min onto a trap column (100μm × 2 cm, PepMap nanoViper C18 column, 5μm, 100 Å, Thermo Scientific) equilibrated in 0.1% trifluoroacetic acid (TFA). The trap column was washed for 3 min at the same flow rate with 0.1% TFA, then switched in-line with a Thermo Scientific, resolving C18 column (75μm × 50 cm, PepMap RSLC C18 column, 2μm, 100 Å). Peptides were eluted from the column at a constant flow rate of 300nl/min with a linear gradient from 3% buffer B to 6% buffer B in 5 min, then from 6% buffer B to 35% buffer B in 115 min, and finally to 80% buffer B within 7 min. The column was then washed with 80% buffer B for 4 min and re-equilibrated in 3% buffer B for 15 min. Two blanks were run between each sample to reduce carryover. The column was kept at a constant temperature of 50°C.

The data was acquired using an easy spray source operated in positive mode with spray voltage at 2.445 kV, and the ion transfer tube temperature at 250°C. The MS was operated in DIA mode. A scan cycle comprised a full MS scan (m/z range from 350 to 1650), with RF lens at 40%, AGC target set to custom, normalised AGC target at 300%, maximum injection time mode set to custom, maximum injection time at 20 ms, microscan set to 1 and source fragmentation disabled. MS survey scan was followed by MS/MS DIA scan events using the following parameters: multiplex ions set to false, collision energy mode set to stepped, collision energy type set to normalized, HCD collision energies set to 25.5, 27 and 30%, orbitrap resolution 30,000, first mass 200, RF lens 40%, AGC target set to custom, normalized AGC target 3000%, microscan set to 1 and maximum injection time 55ms. Data for both MS scan and MS/MS DIA scan events were acquired in profile mode.

#### Naïve vs LPS + IL-4 (Normal/CF +/- Fluvastatin)

Peptides were analysed using single-shot Data Independent Acquisition (DIA). For each sample 200ng of peptide was injected onto a C18 reverse-phase chromatography system (Vanquish, Thermo Scientific) and electrosprayed into an Astral Orbitrap Mass Spectrometer (Thermo Fisher). The following buffers were used for liquid chromatography: buffer A (0.1% formic acid in Milli-Q water (v/v)) and buffer B (80% acetonitrile and 0.1% formic acid in Milli-Q water (v/v). Samples were loaded onto a trap column (Pep Map Neo C18, 5µm, 300µm x5 mm) Thermo Scientific) equilibrated in buffer A. The column (Pep MAP RSLC C18, 2µm, 150 µm x15 cm) was equilibrated with 4% buffer and peptides were eluted at a variable flow rate, starting at 1.3 µl/min to 0.8 µl/min from 4% buffer B to 8% buffer B in 0.1 min, then from 8% buffer B to 22.5% buffer B in 13 min, and from 22.5% b to 35% B in 6.90 min. The low rate is increased from 0.8 µl/min to 2 µl/min in 0.4 min with buffer B increasing from 35% to 55%. The column is finally washed at a flow of 2µl/min with 99% buffer B. The column was kept at a constant temperature of 50°C. The data was acquired using an easy spray source operated in positive mode with spray voltage at 2.0 kV, and the ion transfer tube temperature at 280°C. The MS was operated in DIA mode. A scan cycle comprised a full MS scan (m/z range from 380-980), with orbitrap resolution at 2400000, RF lens at 40%, AGC target set to custom, normalised AGC target at 500%, absolute AGC value set to 5.00e6, maximum injection time 5ms and micros-can set to 1. MS survey scan was followed by MS/MS DIA scan events using the following parameters: DIA window type set to auto, isolation window 2, window overlap set to 0, window placement optimisation on, number of scan events 299, collision energy type normalized, HCD collision energy 25%, detector type Astral, scan range 150-2000, normalized AGC target 500%, absolute AGC target 5.000e4, maximum injection time 5 ms, microscan set to 1, loop control time, time 0.6s. Data for both MS scan and MS/MS DIA scan events were acquired in profile mode.

### Proteomic data analysis

For comparing naïve and stimulated B cells, raw mass spec data files were searched using Spectronaut version 19 (Biognosys). Data was analysed by DirectDIA. Raw mass spec data files were searched against a mouse database (Swissprot Trembl November 2023) with the following parameters: directDIA, false discovery rate set to 1%, protein N-terminal acetylation and methionine oxidation were set as variable modifications and carbamidomethylation of cysteine residues was selected as a fixed modification.

The following Spectronaut settings were used (Baker *et al*, 2024): all identification settings were set to 0.01 (precursor q-value cut-off, precursor PEP cut-off, protein FDR strategy (accurate), protein q-value cut-off (experiment), protein q-value cut-off (run) and protein PEP cut-off). The following quantification settings were used: ‘Quant 2.0’, the MS-Level quantity was set to ‘MS2’, imputation was disabled, major group Top N and minor group Top N were set as ‘False’ and cross run normalisation was set as ‘False’.

Intensity values from Spectronaut were further analysed in Perseus (Tyanova *et al*, 2016) and the histone ruler method (Wiśniewski *et al*, 2014) used to estimate copy number and concentrations of the identified proteins. Fold change was calculated relative to levels in naïve B cells and significance determined using unpaired Student’s t tests on log10-transformed data (Table S3 and Table S4). The False Discovery Rate approach using the 2-stage step up method, Benjamini, Krieger, and Yekutieli (Benjamini *et al*, 2006) with a Q value of 0.01 (Table S3). Enrichment against the Gene Ontology (GO) Biological Processes, Uniprot Keyword Biological Process and *Kyoto Encyclopaedia of Genes and Genomes* (KEGG) databases. Biological Process GO terms (Gene Ontology Consortium *et al*, 2023) and KEGG pathways (Kanehisa *et al*, 2016) was carried out using the DAVID gene enrichment site (Sherman *et al*, 2022) (Table S1).

### Statistical analysis

Unless otherwise stated, graphs show mean values + SD with individual replicates shown by symbols. Unpaired two-tailed Student’s *t* test, one-way ANOVA, two-way ANOVA with post-hoc testing were performed in Prism, and the tests used are indicated in the figure legends. Full results of the ANOVA calculations are given in Supplemental Table S5.

## Supporting information

Figures S1-16, Table S2 - serum tests, Table S4 - statistics

## Acknowledgments

The authors thank A. Rennie and R. Clarke from the Flow Cytometry Facility for cell sorting and advice on flow cytometry. We also thank the FingerPrints Proteomics facility for running our samples for mass spectrometry and for advice on proteomics. This research was supported by a Wellcome Trust (DMSC, UNS148487) and MRC (MR/N013735/1, MR) PhD studentship awards.

## Author contributions

D.M.S.C., M.R., S.A., performed the experiments. A.J.M. performed the liquid chromatography–mass spectrometry. D.M.S.C and J.S.C.A analysed the proteomics data. L.V.S designed the kynurenine uptake and protein synthesis experiments. M.S optimised the silica depletion and filipin staining experiments. D.M.S.C and J.S.C.A wrote the manuscript.

## Resource availability Lead contact

Further information and requests for resources should be directed to and will be fulfilled by the lead contact, Simon Arthur.

## Materials availability

This study did not generate any new unique reagents.

## Data availability

The mass spectrometry proteomics data (Naïve vs LPS + IL-4) has been deposited to the ProteomeXchange Consortium (http://proteomecentral.proteomexchange.org) via the PRIDE partner repository (Perez-Riverol *et al*, 2024) with the dataset identifier PXD057739.

The mass spectrometry proteomics data (Naïve vs LPS + IL-4 (Normal/CF) +/-Fluvastatin has been deposited to PRIDE with the dataset identifier PXD074568

Flow cytometry data that support the findings of this study are available from the corresponding author upon request. Any additional information required to reanalyse the data reported in this paper is available from the lead contact upon request.

## Supplemental information

Document S1. Figures S1–S16, Table S2 - Serum tests related to Figure 5 and Table S5 - Statistics table related to all figures.

Supplemental Table S1. Excel file containing terms for protein enrichment too large to fit in a pdf. Related to Figure 1H.

Supplemental Table S3. Excel file containing proteomic data set (Naïve vs LPS + IL-4)

Supplemental Table S4. Excel file containing proteomic data set (Naïve vs LPS + IL-4 (Normal/CF +/- Fluvastatin)

