## Supplementary material for "Lipopolysaccharide stimulates dynamic changes in B cell metabolism to promote proliferation": Figures S1-16, Table S2 - serum tests, Table S4 - statistics

<u>Supplementary information</u>

Supplementary figures 1 – 16

Supplementary tables 2 and 5

*Supplementary tables 1, 3 and 4 are provided as separate Excel files*

#### Supplementary Figure 1

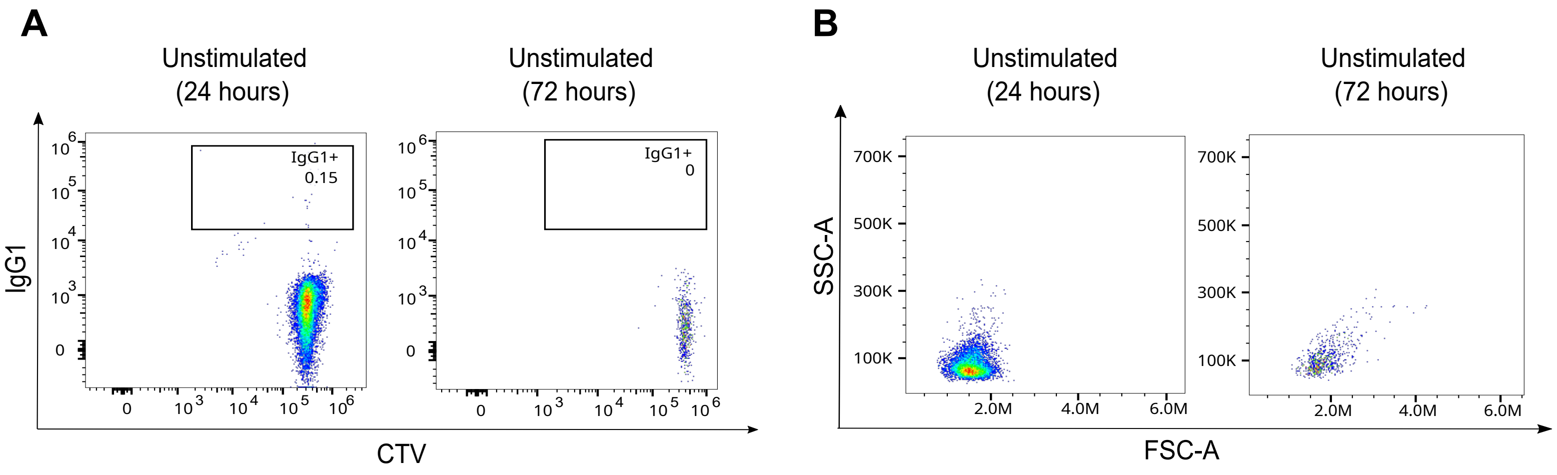

**Figure S1. Unstimulated B cells do not proliferate or undergo class switching. Related to Figure 1.**

**(A-B)** Lymph node cells from WT mice were stained with Cell Trace Violet (CTV), cultured for 24 or 72 hours and analysed by flow cytometry. Live CD19+ B cells were identified using the gating strategy described in Figure S14A. **(A)** Representative flow cytometry plots comparing IgG1 expression and CTV staining after 24 or 72 hours. **(B)** Representative plots for FSC and SSC at 24 or 72 hours.

### Supplementary Figure 2

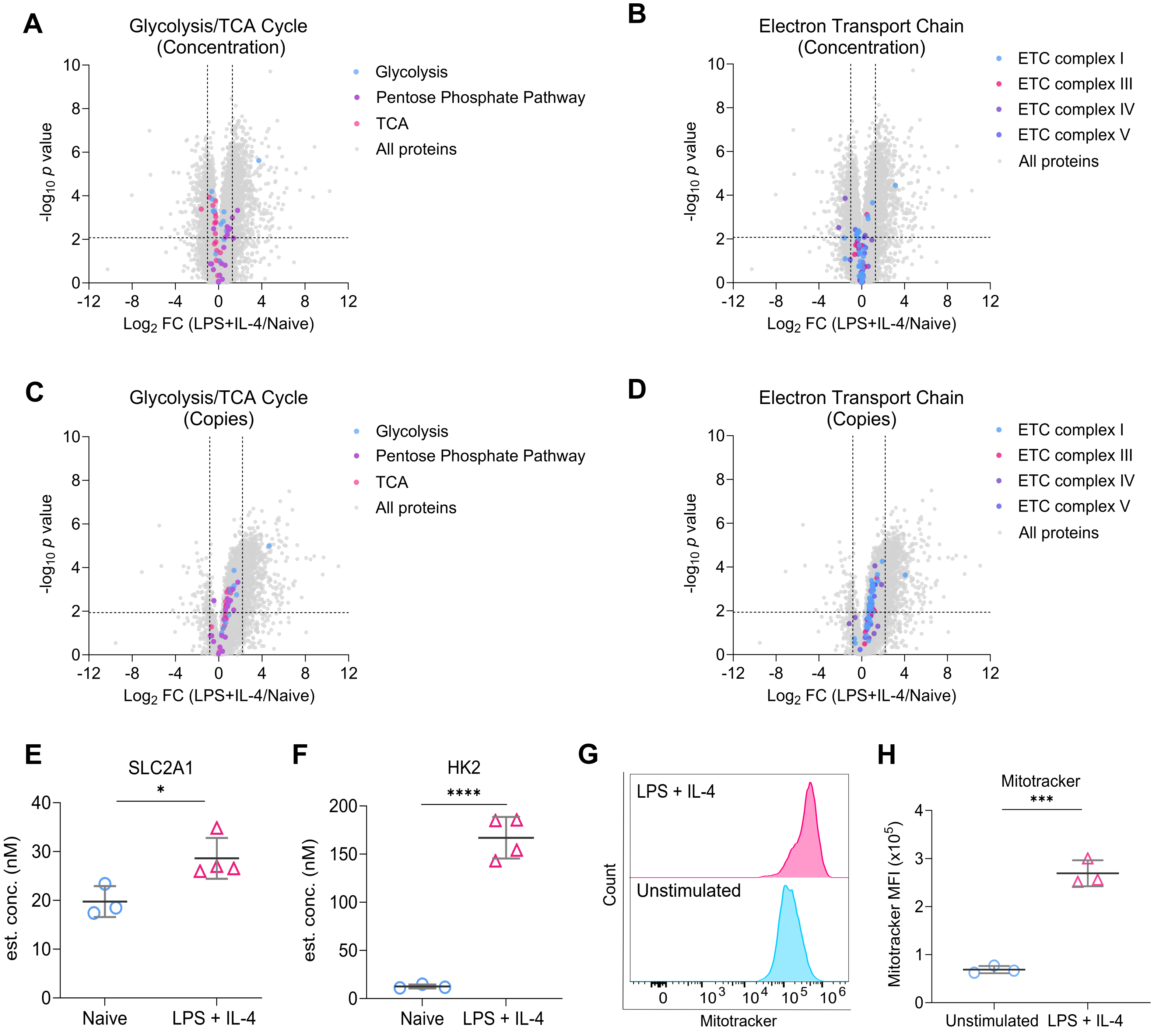

**Figure S2. LPS + IL-4 stimulation upregulates glycolysis in B cells. Related to Figure 1.**

**(A)** Volcano plot showing the cellular protein concentration (μM) of proteins involved in glycolysis, the citric acid cycle (TCA) and pentose phosphate pathway (PPP) between naïve vs LPS + IL-4 stimulated B cells derived from the proteomic dataset described in figure 1.
**(B)** Volcano plot showing the cellular protein concentration (μM) of proteins involved in the electron transport chain (ECT) between naïve vs LPS + IL-4 stimulated B cells.
**(C)** Volcano plot depicting changes in protein copy number of proteins involved in glycolysis, the TCA cycle and PPP between naïve vs LPS + IL-4 stimulated B cells.
**(D)** Volcano plot depicting changes in protein copy number of proteins involved in the ECT between naïve vs LPS + IL-4 stimulated B cells. For **(A-D)** horizontal dashed lines indicate $q<0.05$. Vertical dashed lines indicate log2 fold change of one standard deviation away from the median.
**(E, F)** Graphs depicting changes in cellular concentration (nM) of **(E)** SLC2A1 and **(F)** HK2. *p*(adj)<0.05 is indicated by * and *p(adj)*<0.0001 by ****, based of the FDR calculations on the proteomic dataset described in figure 1.
**(G, H)** Splenocytes from WT mice were stimulated with LPS (20μg/ml) and IL-4 (10ng/ml) for 24 hours before staining with MitoTracker to measure mitochondrial volume. Gating strategy described in Figure S15C. Representative histograms are shown in **(G)** and quantification of MFI of MitoTracker staining between naïve and LPS + IL-4 stimulated B cells in **(H)**. The graph shows three technical replicates from cells isolated from one mouse and is representative of three independent experiments. Statistical power was determined using an unpaired Student's t-test (two-tailed), where $p<0.001$ is indicated by ***.

### Supplementary Figure 3

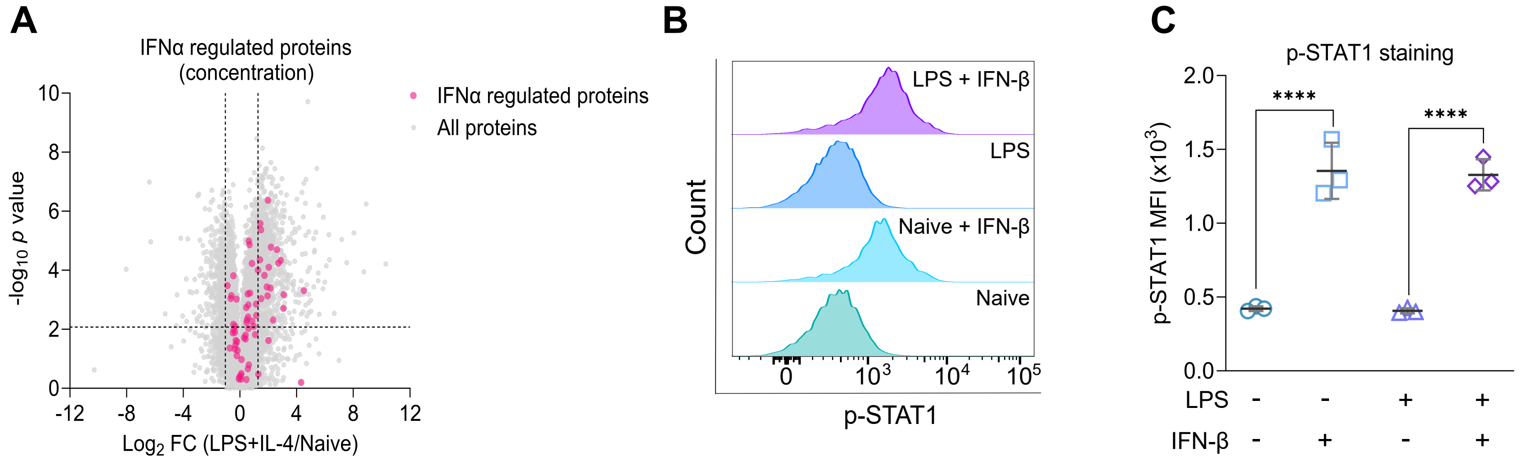

**Figure S3. LPS + IL-4 stimulation does not increase STAT1 phosphorylation. Related to Figure 1.**

**(A)** Volcano plot showing the cellular protein concentration (μM) of proteins regulated by IFNα between naïve vs LPS + IL-4 stimulated B cells derived from the proteomic dataset described in Figure 1. A list of B cell specific genes upregulated by IFNα stimulation from (Mostafavi et al, 2016) (Table S1. Transcripts induced by systemic IFNa in different mouse immunocyte lineages).
**(B, C)** Splenocytes from C57BL6/J mice were stimulated with LPS (20μg/ml) and/or IFN-β (10ng/ml) for 15 minutes before fixing and staining for p-STAT1. Representative histograms are shown in **(B)** and quantification of the MFI of p-STAT1 staining between naïve, IFN-β, LPS and LPS + IFN-β stimulated B cells in **(C)**.
The graph shows three technical replicates from cells isolated from one mouse and is representative of three independent experiments. Statistical power was determined using one-way ANOVA followed by multiple comparison testing via Dunnett's analysis, where $p$<0.0001 is indicated by ****.

#### Supplementary Figure 4

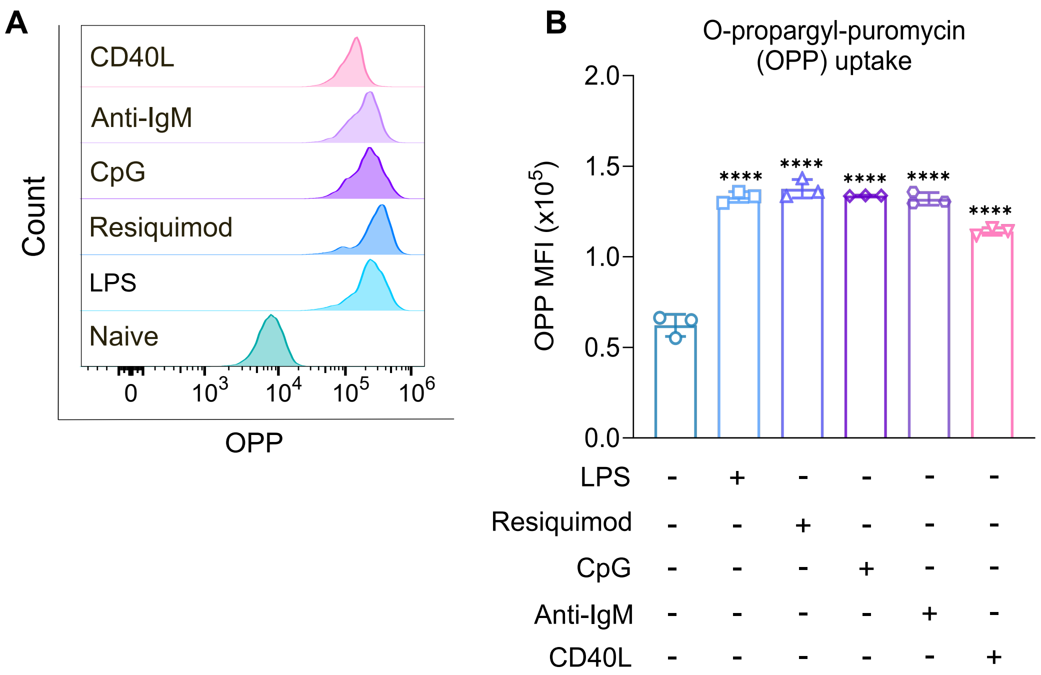

**Figure S4. Protein synthesis is increased in B cells regardless of stimuli. Related to Figure 3.**

Splenocytes from C57BL6/J mice were plated in normal media before stimulation with LPS (20µg/ml), Resiquimod (1µg/ml), ODN 1826 (1µg/ml), Anti-IgM (10µg/ml), or CD40L (500ng/ml) for 24 hours before fixing and staining for the uptake of puromycin analog O-propargyl-puromycin (OPP) to measure protein synthesis. Gating strategy for OPP staining is shown in Figure S14C.

**(A)** Representative histogram comparing OPP uptake between different stimuli. **(B)** Quantification shows the results of three biological replicates. Statistical power was determined using one-way ANOVA followed by multiple comparison testing via Dunnett's analysis. For comparisons to naïve B cells, $p<0.0001$ is indicated by ****.

**Supplementary Figure 5**

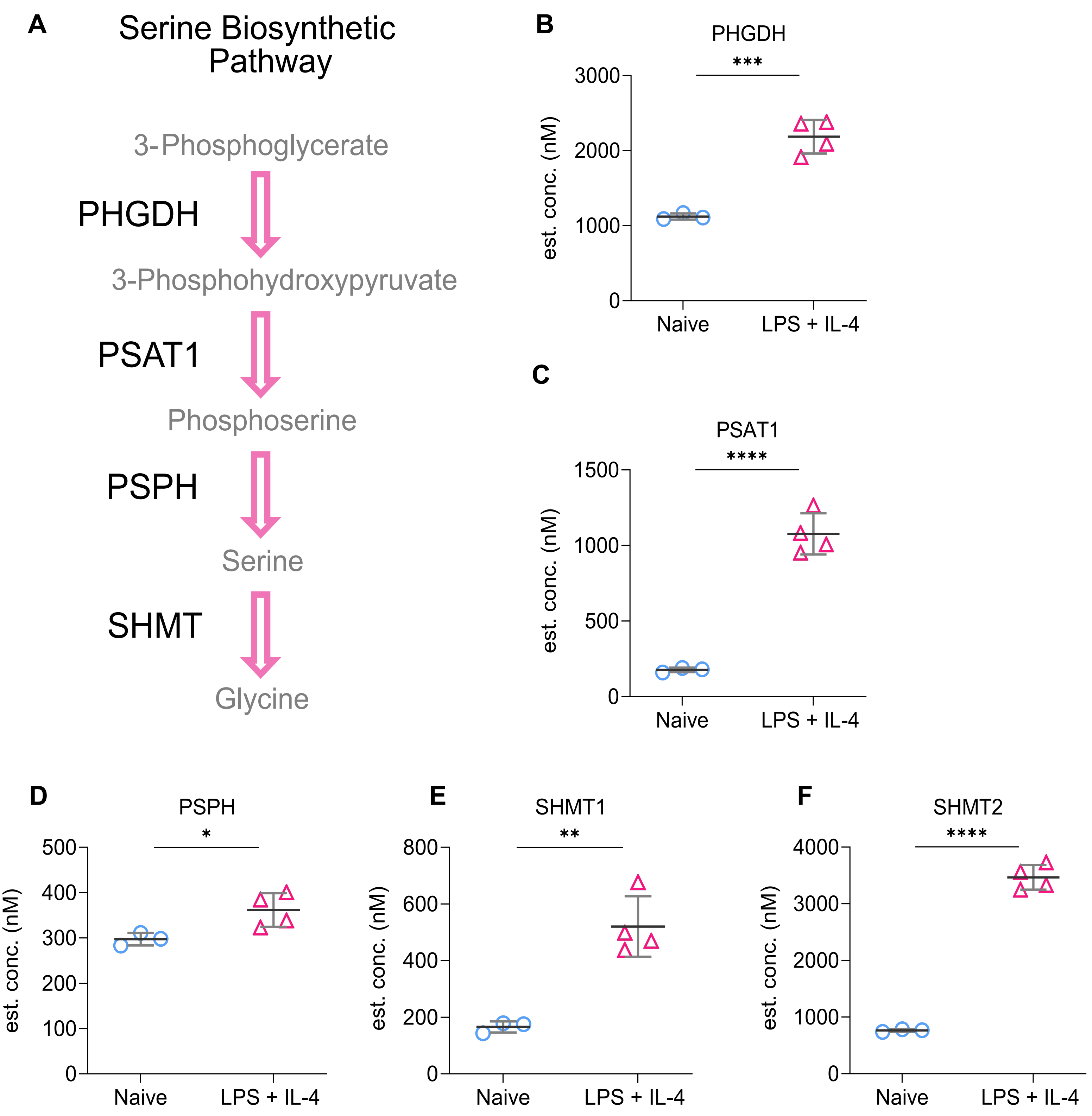

**Figure S5. LPS + IL-4 stimulation upregulates serine biosynthesis. Related to Figure 4.**

**(A)** Schematic depicting the serine biosynthetic pathway. **(B-F)** Graphs depicting changes in cellular concentration (nM) of **(B)** PHGDH, **(C)** PSAT1, **(D)** PSPH, **(E)** SHMT1 and **(F)** SHMT2 determined from the proteomic dataset described in Figure 1. *p(adj)*<0.05 is indicated by *, *p(adj)*<0.01 by **, *p(adj)*<0.001 by *** and *p(adj)*<0.0001 by **** as determined from the FDR calculations on the proteomic dataset.

### Supplementary Figure 6

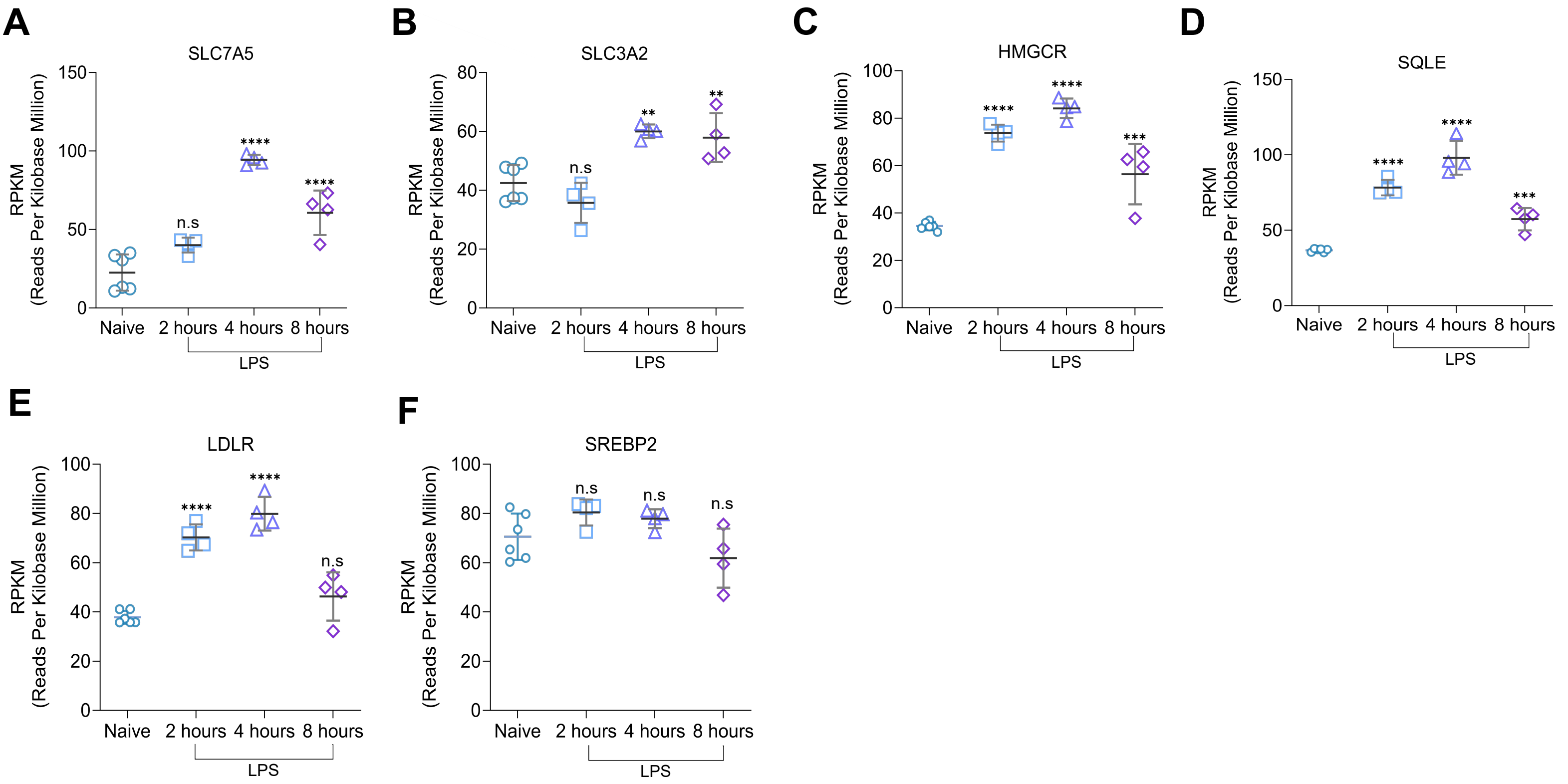

**Figure S6. Short term LPS stimulation increases the transcription of genes involved in amino acid uptake and cholesterol metabolism in B cells. Related to Figues 4 and 5.**

RNA-Seq data comparing reads per kilobase million (RPKM) between naive and LPS stimulated B cells at different time points from wild-type C57BL/6 mice derived from dataset EV1 (Tesi et al, 2019). Reads per kilobase million (RPKM) of **(A)** SLC7A5, **(B)** SLC3A2, **(C)** HMGCR, **(D)** SQLE, **(E)** LDLR, and **(F)** SREBP2. Statistical power for the remaining figures was determined using one-way ANOVA followed by multiple comparison testing via Dunnett's analysis. For comparisons to naïve B cells, $p$<0.01 is indicated by **, $p$<0.001 by *** and $p$<0.0001 by ****. ns indicated by $p$>0.05.

**Supplementary Figure 7**

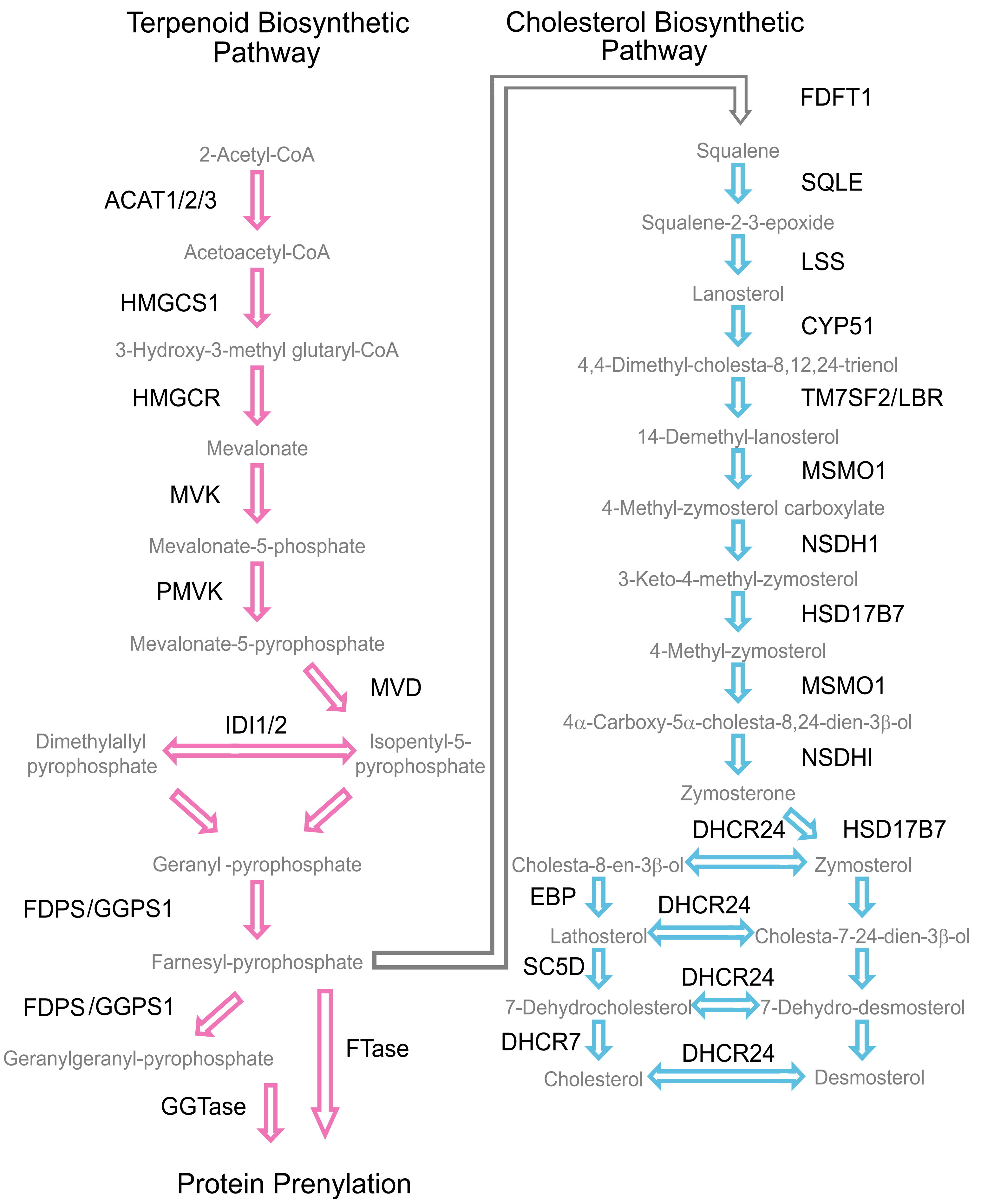

**Figure S7. Terpenoid and Cholesterol biosynthetic pathway. Related to Figure 5.**

Schematic depicting the terpenoid and cholesterol biosynthetic pathways.

### Supplementary Figure 8

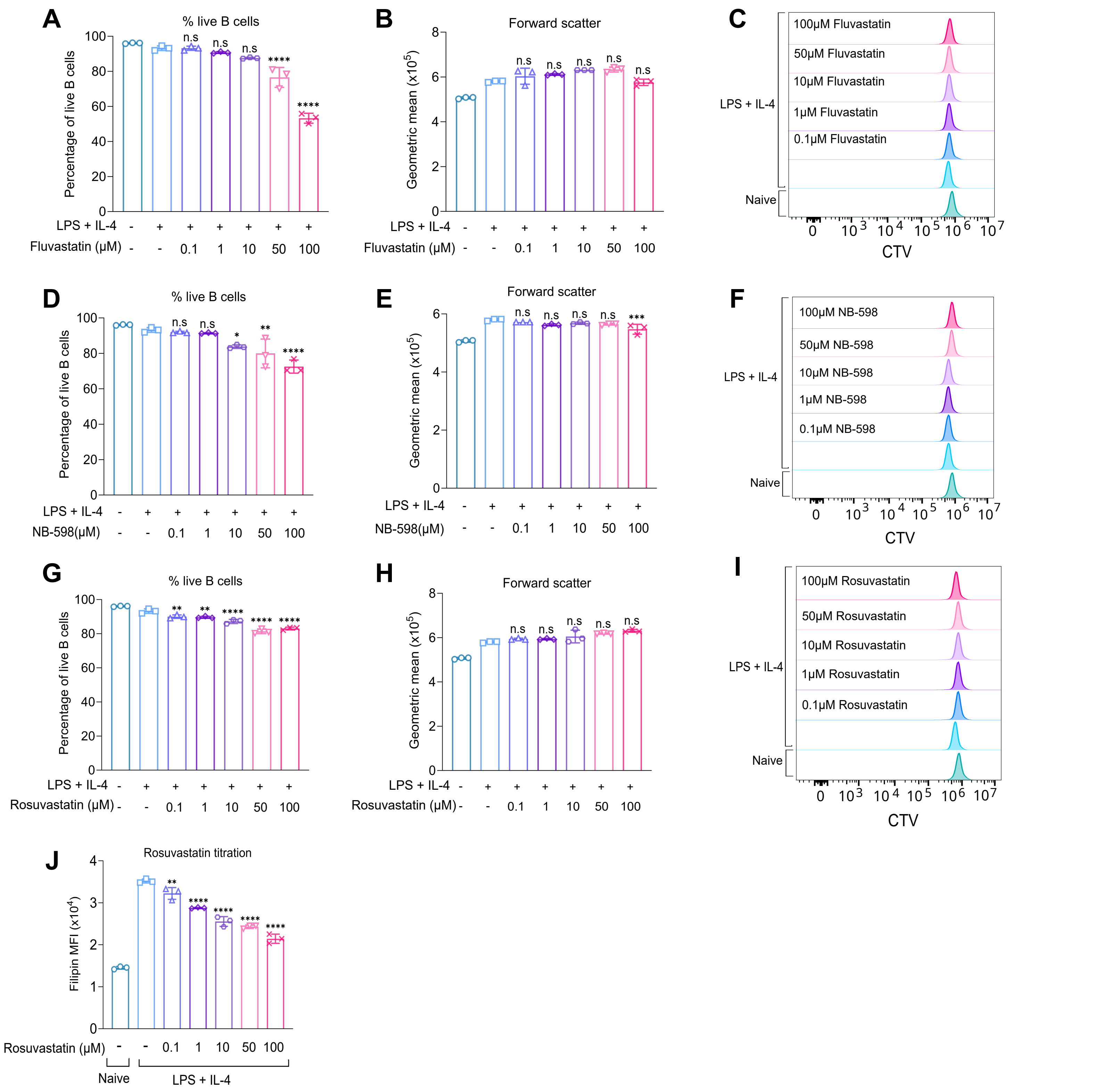

**Figure S8. Blocking rate-limiting enzymes in the cholesterol biosynthesis pathway reduces B cell growth, survival and proliferation. Related to Figure 5.**

B cells were purified from the spleens of WT mice and cultured in normal media. The cells were stained with CTV and pre-treated with DMSO as a vehicle control or varying concentrations of Fluvastatin, NB-598 or Rosuvastatin for 45 minutes before stimulation with LPS (20μg/ml) and IL-4 (10ng/ml) for 24 hours. For all panels, cells in the absence of LPS + IL-4 were naïve B cells analysed on the day of isolation.

**(A-C)** relates to the data in Figure 5G and shows the percentage of live (7AAD-ve) B cells **(A)**, forward scatter of B cells **(B)** and representative histograms for CTV staining **(C)** for Fluvastatin treatment.

**(D-F)** relates to the data in Figure 5H and shows the percentage of live (7AAD-ve) B cells **(D)**, forward scatter of B cells **(E)** and representative histograms for CTV staining **(F)** for NB-598 treatment.

**(G-I)** Graphs show the percentage of live (7AAD-ve) B cells **(G)**, forward scatter of B cells **(H)** and representative histograms for CTV staining **(I)** for Rosuvastatin treatment.

**(J)** Splenocytes from WT mice were plated in cholesterol-free media and pre-treated with DMSO as a vehicle control or varying concentrations of Rosuvastatin for 45 minutes before stimulation with LPS (20μg/ml) and IL-4 (10ng/ml). The cells were fixed after 24 hours and stained with filipin. **(J)** Filipin staining of Rosuvastatin titration.

Graphs show three technical replicates from cells isolated from one mouse and is representative of three independent experiments. Statistical power was determined using one-way ANOVA followed by multiple comparison testing via Dunnett's analysis, where $p < 0.05$ is indicated by *, $p < 0.01$ by **, $p < 0.001$ by *** and $p < 0.0001$ by **** for comparison to LPS + IL-4 condition.

### Supplementary Figure 9

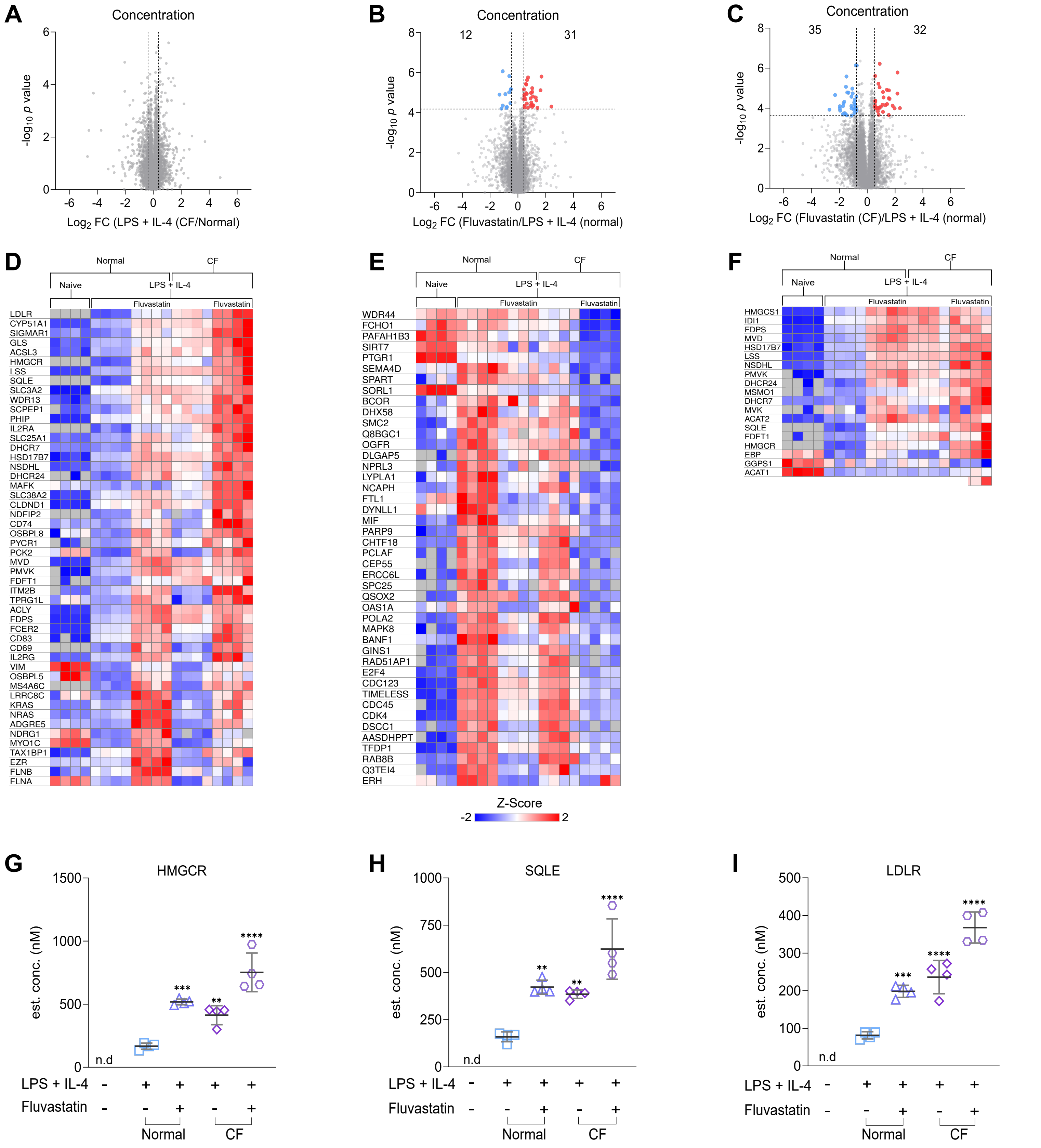

**Figure S9. Blocking cholesterol biosynthesis or uptake upregulates proteins involved in cholesterol metabolism in B cells. Related to Figure 6.**

B cells were purified from the spleens of C57BL6/J mice and cultured in normal or cholesterol-free (CF) media. The cells were then pre-treated with DMSO as a vehicle control or Fluvastatin (10µM), where indicated, for 45 minutes prior to stimulation with LPS (20µg/ml) and IL-4 (10µg/ml) for 24 hours. Alternatively, naïve B cells were lysed directly. Cells were lysed and analysed by proteomics as described in the methods. Samples from four mice for each condition were generated. **(A)** Volcano plot showing the estimated cellular protein concentration (µM) of LPS + IL-4 stimulated B cells cultured in normal or CF media. **(B)** Volcano plot showing the estimated cellular protein concentration (µM) of LPS + IL-4 stimulated B cells cultured in normal media +/- Fluvastatin. **(C)** Volcano plot showing the estimated cellular protein concentration (µM) of LPS + IL-4 stimulated B cells cultured in normal media or CF media + Fluvastatin. Horizontal dashed lines indicate *q*<0.05. Vertical dashed lines indicate log2 fold change of one standard deviation away from the median. Heat map showing the expression of proteins significantly upregulated **(D)** or downregulated **(E)** derived from the proteomic data. **(F)** Heatmap of all the enzymes involved in cholesterol biosynthesis. **(G-J)** Graphs depicting changes in cellular concentration (nM) of **(G)** HMGCR, **(H)** SQLE and **(I)** LDLR. Statistical power was determined using a one-way ANOVA followed by multiple comparison testing via Dunnett's analysis, where *p(adj)*<0.01 is indicated by **, *p(adj)*<0.001 by ***, *p(adj)*<0.0001 by ****, as determined from the FDR calculations on the proteomic dataset. Statistical power for **(J)** was determined using Welch's ANOVA to account for unequal variances followed by multiple comparison testing via Dunnett's T3 analysis. For comparison *p(adj)*<0.01 is indicated by **. ns by *p(adj)*>0.05 for comparisons to the LPS + IL-4 condition in normal media.

#### Supplementary Figure 10

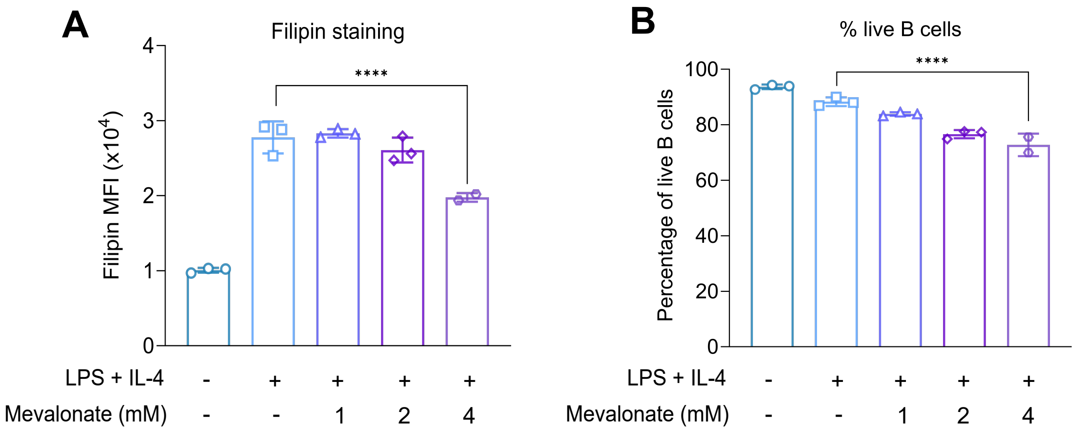

**Figure S10. High levels of exogenous mevalonate are toxic to B cells. Related to Figure 7.** Splenocytes were plated in normal media and pre-treated with HEPES as a vehicle control or mevalonate at the stated concentrations for 1 hour prior to LPS (20µg/ml) and IL-4 (10ng/ml) stimulation. The cells were cultured for 24 hours, then fixed and stained with filipin prior to acquisition. Gating strategy described in Figure S15A. **(A)** Filipin staining. **(B)** Percentage of live B cells (L/D-ve). The graph shows three technical replicates from cells isolated from one mouse and is representative of two independent experiments. Statistical power for was determined using one-way ANOVA followed by multiple comparison testing via Dunnett's analysis, where $p<0.0001$ is indicated by ****.

### Supplementary Figure 11

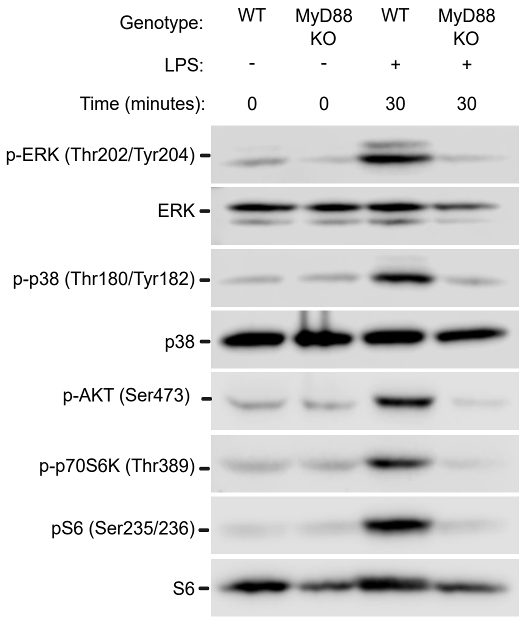

**Figure S11. MyD88 is required for signalling through TLR4. Related to Figure 9.**

B cells were purified from the spleens of WT and MyD88 KO mice then stimulated with LPS (20µg/ml). The cells were lysed after 60 minutes. Western blotting was used to probe samples with the appropriate antibody (Table 4).

### Supplementary Figure 12

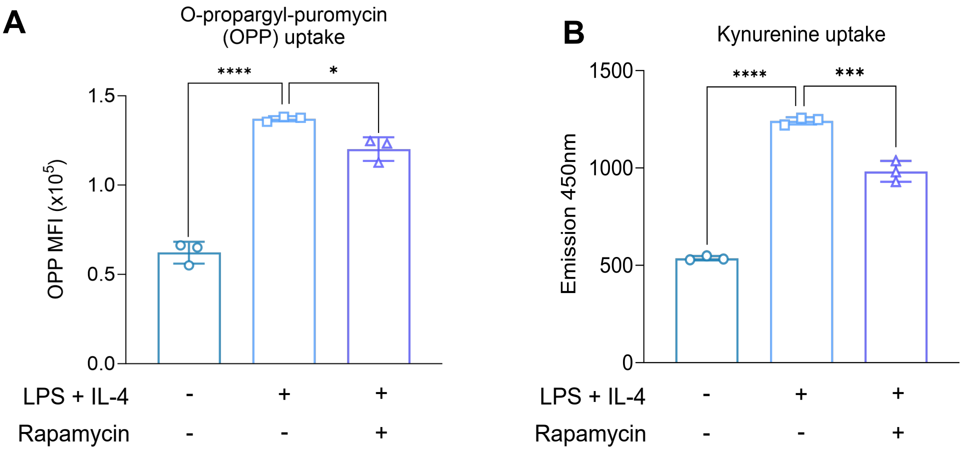

**Figure S12. Inhibition of mTOR impacts B cell function. Related to Figure 9.**

**(A)** Splenocytes from C57BL6/J mice were pre-treated with DMSO as a vehicle control or rapamycin (20nM), where indicated, for 45 minutes. The cells were stimulated with LPS (20µg/ml) and IL-4 (10µg/ml) for 24 hours before fixing and staining for the uptake of puromycin analog O-propargyl-puromycin (OPP) to measure protein synthesis. Gating strategy for OPP staining is shown in Figure S14C. **(A)** OPP uptake after rapamycin treatment. Data shows the results of three biological replicates.

**(B)** B cells were purified from the spleens of C57BL6/J mice, then pre-treated with DMSO as a vehicle control or rapamycin (20nM), where indicated, for 45 minutes. The cells were stimulated with LPS (20µg/ml) and IL-4 (10µg/ml) for 24 hours before fixing and staining for the uptake of kynurenine to measure amino acid uptake. Gating strategy for kynurenine uptake in Figure S14D. **(B)** Kynurenine uptake after rapamycin treatment.

Data shows the results of three biological replicates. Statistical power was determined by one-way ANOVA followed by multiple comparison testing via Dunnett's analysis, where $p<0.05$ is indicated by *, $p<0.001$ by *** and $p<0.0001$ by ****.

#### Supplementary Figure 13

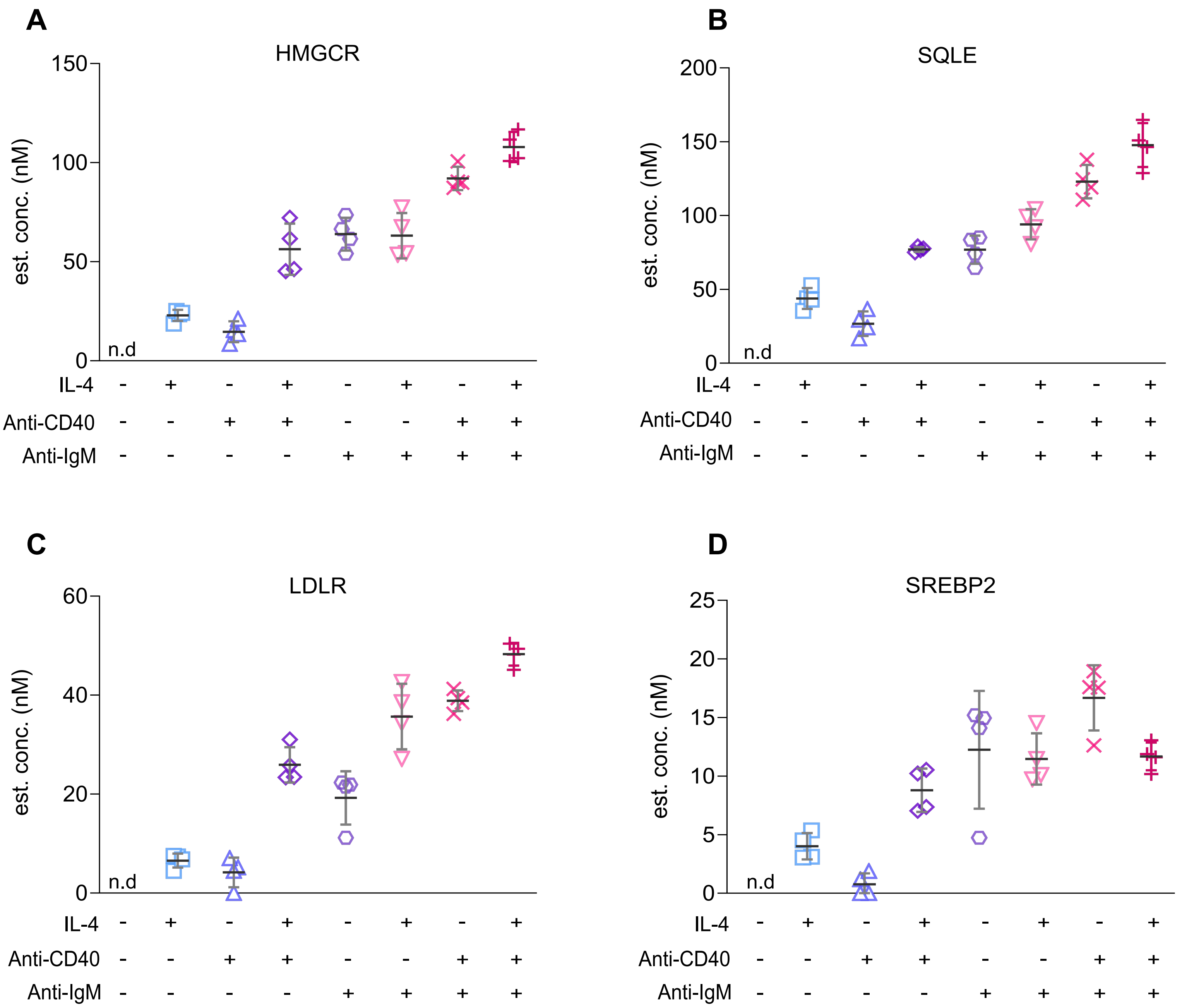

**Figure S13. IL-4, anti-IgM and anti-CD40 stimulation upregulates proteins involved in cholesterol metabolism in B cells. Related to Figure 10.**

Proteomic data comparing protein concentration (nM) between naïve, IL-4, anti-IgM or anti-CD40 stimulated B cells for 40 hours from wild-type C57BL/6 mice derived from (James et al, 2024). **(A-D)** Graphs depicting changes in cellular concentration (nM) of **(A)** HMGCR, **(B)** SQLE, **(C)** LDLR and **(D)** SREBP2.

### Supplementary Figure 14

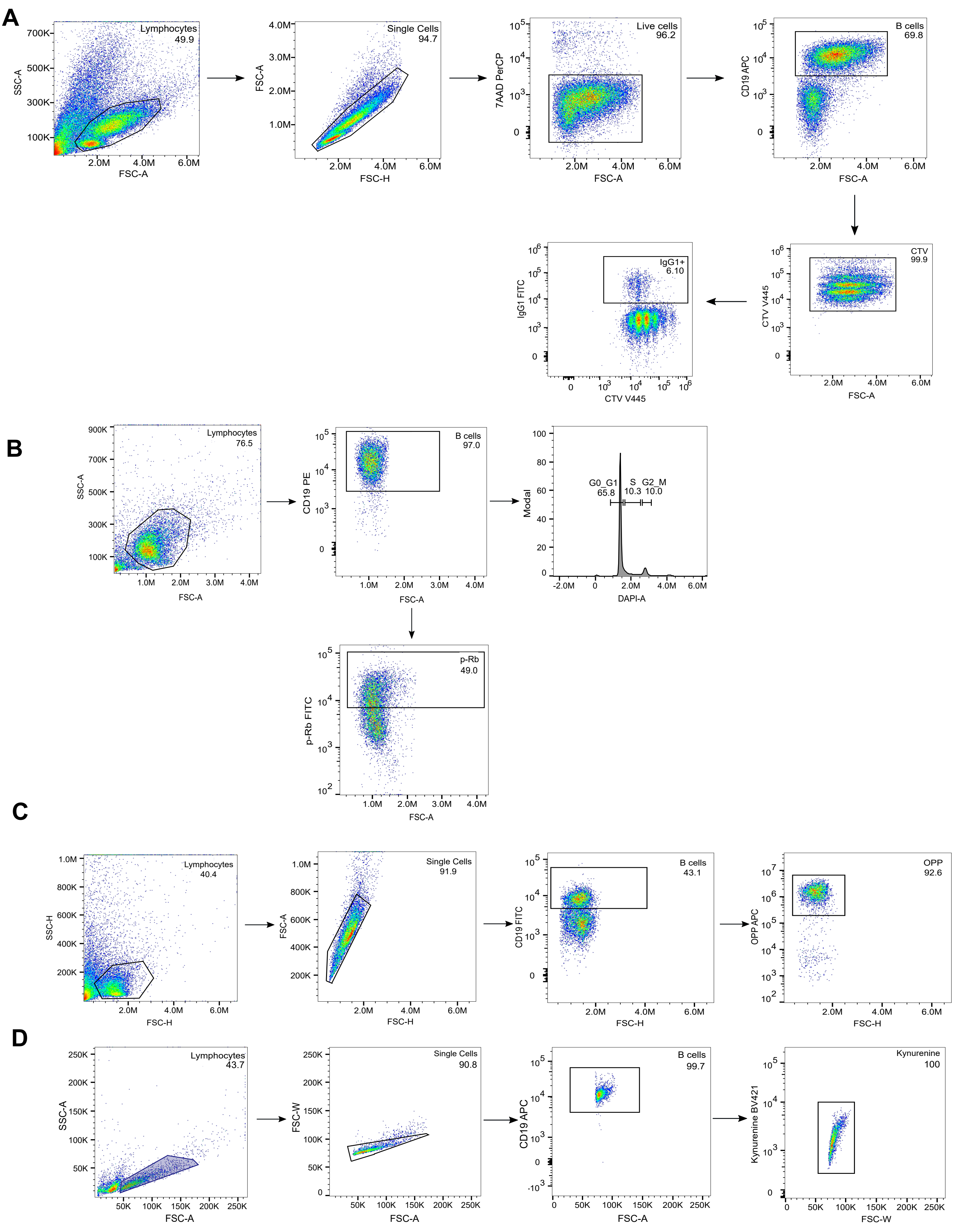

**Figure S14. Representative gating strategies**

**(A)** Gating strategy for lymph nodes and splenocytes cell size, proliferation, CTV, IgG1 (Figure 1A-D).
**(B)** Gating strategy for cell cycle analysis (Figure 2A, B, G, H).
**(C)** Gating strategy for OPP uptake (Figure 3H-I, Figure S4A-B, Figure S12A).
**(D)** Gating strategy for kynurenine uptake (Figure 4J-K, Figure S12B).

#### Supplementary Figure 15

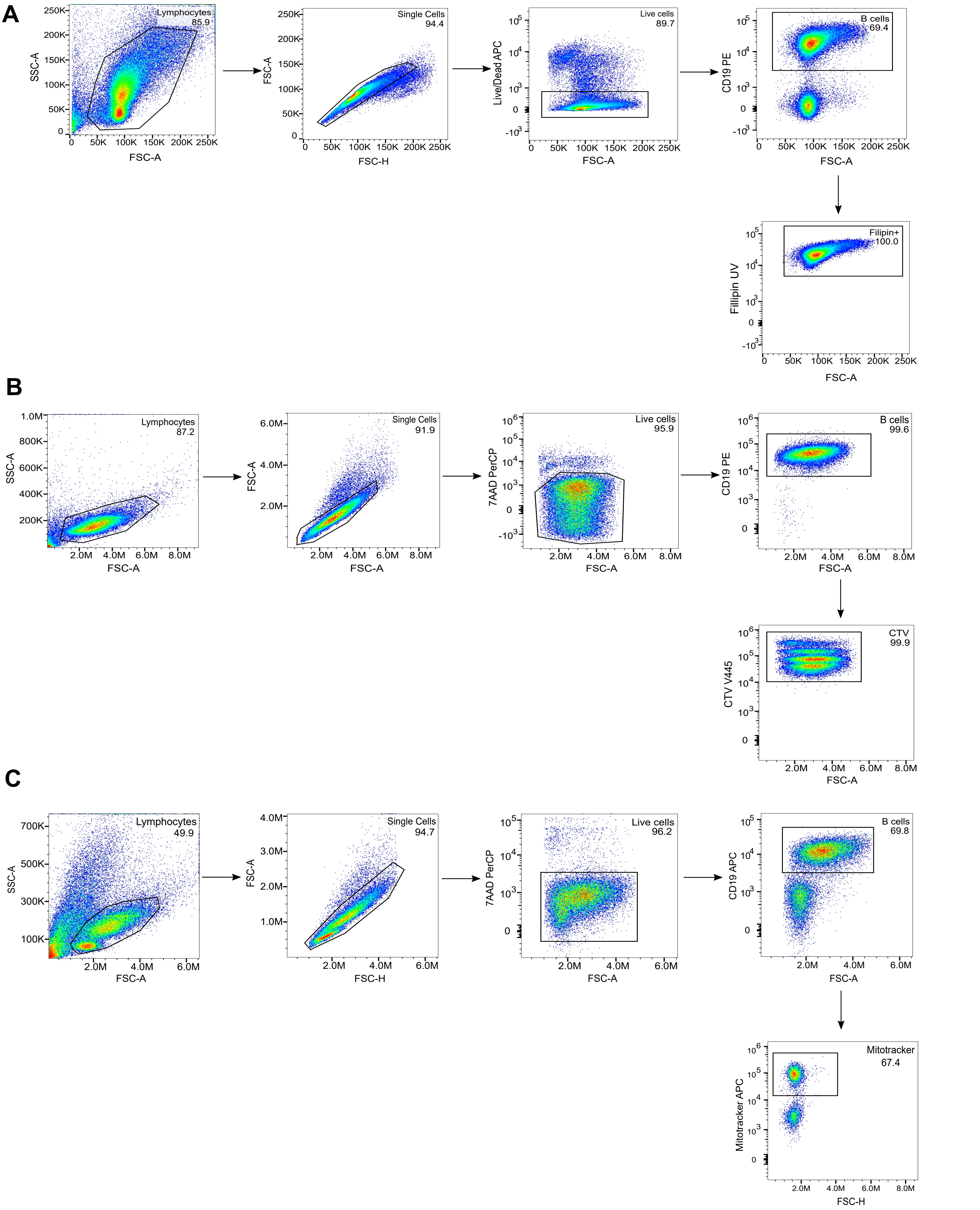

**Figure S15. Representative gating strategies**

**(A)** Gating strategy for filipin staining (Figue 5F-I, Figure 7A-B, Figure 8I-J, Figure 9F, Figure 10A-B, Figure S8J, Figure S10A-B .

**(B)** Gating strategy for purified B cells for cell size, proliferation and CTV (Figure 6A-P, Figure 7C-F, Figure 8A-H,Figure 9A-E, Figure 10C-E, Figure S8A-I).

**(C)** Gating strategy for MitoTracker (Figure S2G-H).

### Supplementary Figure 16

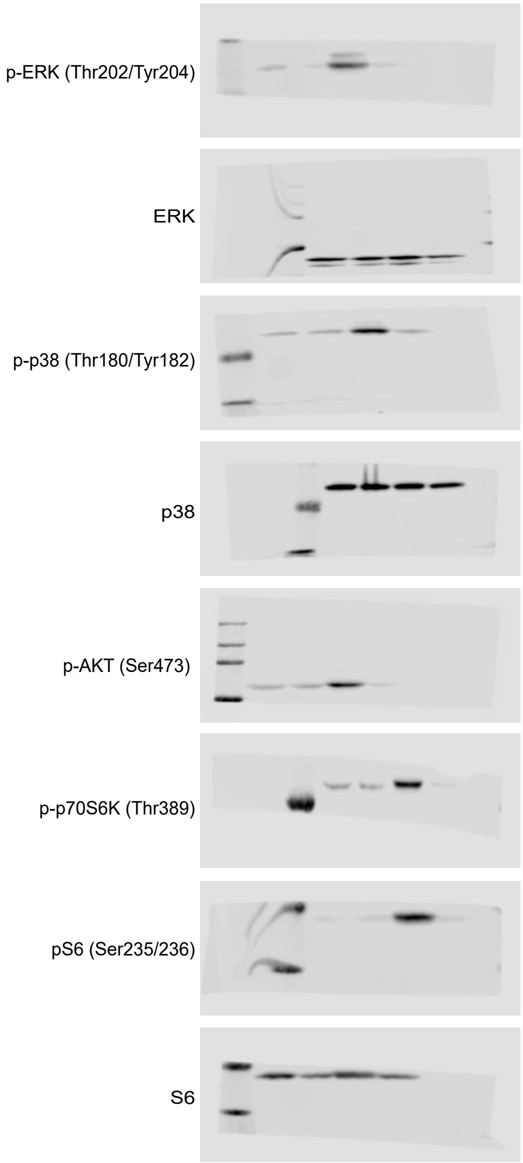

**Figure S16. MyD88 is required for signalling through TLR4. Related to Figure S11.**

B cells were purified from the spleens of WT and MyD88 KO mice then stimulated with LPS (20µg/ml). The cells were lysed after 60 minutes. Western blotting was used to probe samples with the appropriate antibody (Table 4).

**Table S2. Serum tests related to Figure 5**

This table presents the results from the Cholesterol/Lipid profile carried out by London Health Company, comparing differences between normal FBS and cholesterol-free FBS.

| **Normal FBS** | **Biochemistry tests** | | | |
|---|---|---|---|---|
| | Test | Result | Normal Range | Units |
| | **Cholesterol/Lipid Profile** | | | |
| | Triglycerides | 0.73 | <2.3 | mmol/L |
| | Cholesterol | 1 | Optimum <5.0 | mmol/L |
| | HDL Cholesterol | L 0.25 | 0.9-1.7 | mmol/L |
| | LDL Cholesterol | 0.59 | Up to 3.0 | mmol/L |

| **Cholesterol free FBS** | **Biochemistry tests** | | | |
|---|---|---|---|---|
| | Test | Result | Normal Range | Units |
| | **Cholesterol/Lipid Profile** | | | |
| | Triglycerides | 0.67 | <2.3 | mmol/L |
| | Cholesterol | 0.1 | Optimum <5.0 | mmol/L |
| | HDL Cholesterol | L 0 | 0.9-1.7 | mmol/L |
| | LDL Cholesterol | 0.02 | Up to 3.0 | mmol/L |

**Table S5. Statistics table related to all figures**

This table presents the full results of the ANOVA calculations

| Figure | Test | ANOVA<br>F value | ANOVA<br>P value | Test for multiple comparisons | Multiple comparisons (Adjusted P value) | Brown Forsythe test P value |
|---|---|---|---|---|---|---|
| 1C | One-way ANOVA | 1348 | <0.0001 | Sidak's multiple comparison test | Naïve vs IL-4: <0.0001<br><br>Naive vs LPS + IL-4: <0.0001<br><br>IL-4 vs LPS + IL-4 = <0.0001 | 0.2662 |
| 1D | One-way ANOVA | 299.2 | <0.0001 | Sidak's multiple comparison test | Naïve vs IL-4: 0.0002<br><br>Naïve vs LPS + IL-4: <0.0001<br><br>IL-4 vs LPS + IL-4 = <0.0001 | 0.3455 |

| Figure | Test | ANOVA<br>F value | ANOVA<br>P value | Test for multiple comparisons | Multiple comparisons (Adjusted P value) | Brown Forsythe test P value |
|---|---|---|---|---|---|---|
| 2B | Two-way ANOVA | Interaction: 152.6<br><br>Cell cycle phase: 3368<br><br>Stimulation: 9.313 | Interaction: <0.0001<br><br>Cell cycle phase: <0.0001<br><br>Stimulation: 0.0101 | Sidak's multiple comparison test | G0_G1: Naïve vs LPS + IL-4: <0.0001<br><br>S: Naïve vs LPS + IL-4: 0.0001<br><br>G2_M: Naïve vs LPS + IL-4: 0.0023 | |

| Figure | Test | ANOVA F value | ANOVA P value | Test for multiple comparisons | Multiple comparisons (Adjusted P value) | Brown Forsythe test P value |
|---|---|---|---|---|---|---|
| 4E | One-way ANOVA | 1561 | <0.0001 | Dunnett's multiple comparison's test | Naïve vs IL-4: 0.0004<br>Naïve vs LPS + IL-4: <0.0001 | 0.4645 |
| 4J | Two-way ANOVA | Interaction: 106.6<br><br>Condition: 340.1<br><br>Genotype: 1707 | Interaction: <0.0001<br><br>Condition: <0.0001<br><br>Genotype: <0.0001 | Sidak's multiple comparison test | Naïve WT vs LPS + IL-4 WT: <0.0001<br><br>Naïve SLC7A5 KO vs LPS + IL-4 SLC7A5 KO: <0.0001<br><br>WT vs SLC7A5 KO (Kynurenine): <0.0001<br><br>WT vs SLC7A5 KO (BCH): >0.9999<br><br>WT vs SLC7A5 KO (HBSS): <0.0001 | |
| 4K | Two-way ANOVA | Interaction: 108.1<br><br>Condition: 384.4<br><br>Genotype: 774.9 | Interaction: <0.0001<br><br>Condition: <0.0001<br><br>Genotype: <0.0001 | Sidak's multiple comparisons test | Naïve WT vs LPS WT: <0.0001<br><br>Naïve SLC7A5 KO vs LPS SLC7A5 KO: <0.0001<br><br>WT vs SLC7A5 KO (Kynurenine): <0.0001<br><br>WT vs SLC7A5 KO (BCH): 0.9992<br><br>WT vs SLC7A5 KO (HBSS): 0.8046 | |

| Figure | Test | ANOVA F value | ANOVA P value | Test for multiple comparisons | Multiple comparisons (Adjusted P value) | Brown Forsythe test P value |
|---|---|---|---|---|---|---|
| 5F | One-way ANOVA | 932.6 | <0.0001 | Dunnett's multiple comparison's test | Naïve vs 2 hours: 0.0970<br><br>Naïve vs 4 hours: 0.0181<br><br>Naïve vs 8 hours: <0.0001<br><br>Naive vs 16 hours: <0.0001<br><br>Naive vs 24 hours: <0.0001 | 0.3544 |
| 5G | One-way ANOVA | 166.1 | <0.0001 | Dunnett's multiple comparison's test | Naïve vs LPS + IL-4: <0.0001<br><br>LPS + IL-4 vs 0.1μM Fluvastatin: <0.0001<br><br>LPS + IL-4 vs 1μM Fluvastatin: <0.0001<br><br>LPS + IL-4 vs 10μM Fluvastatin: <0.0001<br><br>LPS + IL-4 vs 50μM Fluvastatin: <0.0001<br><br>LPS + IL-4 vs 100μM Fluvastatin: <0.0001 | 0.4365 |
| 5H | One-way ANOVA | 147.0 | <0.0001 | Dunnett's multiple comparison's test | Naïve vs LPS + IL-4: <0.0001<br><br>LPS + IL-4 vs 0.1μM NB-598: 0.0003<br><br>LPS + IL-4 vs 1μM NB-598: <0.0001<br><br>LPS + IL-4 vs 10μM NB-598: <0.0001<br><br>LPS + IL-4 vs 50μM NB-598: <0.0001 | 0.8640 |

| | | | | | |
|---|---|---|---|---|---|
| | | | | | LPS + IL-4 vs 100µM NB-598: <0.0001 |
| 5I | Two-way ANOVA | Interaction: 36.95<br><br>Media 332.8<br><br>Inhibitor: 488.8 | Interaction: <0.0001<br><br>Media: <0.0001<br><br>Inhibitor: <0.0001 | Tukey's multiple comparisons test | Normal<br><br>Naïve vs LPS + IL-4: <0.0001<br><br>LPS + IL-4 vs NB-598: <0.0001<br><br>LPS + IL-4 vs Fluvastatin: <0.0001<br><br>Normal vs Cholesterol-free<br>LPS + IL-4: <0.0001<br><br>Cholesterol-free<br><br>Naïve vs LPS + IL-4: <0.0001<br><br>LPS + IL-4 vs NB-598: <0.0001<br><br>LPS + IL-4 vs Fluvastatin: <0.0001 |

| Figure | Test | ANOVA F value | ANOVA P value | Test for multiple comparisons | Multiple comparisons (Adjusted P value) | Brown Forsythe test P value |
|---|---|---|---|---|---|---|
| 6B | Two-way ANOVA | Interaction: 8.996<br><br>Media: 9.466<br><br>Inhibitor: 728.5 | Interaction: 0.0010<br><br>Media: 0.0072 | Tukey's multiple comparisons test | Normal<br><br>Naïve vs LPS + IL-4: <0.0001<br><br>LPS + IL-4 vs NB-598: <0.0001<br><br>LPS + IL-4 vs Fluvastatin: <0.0001<br><br>Normal vs Cholesterol-free<br>LPS + IL-4: <0.0001<br><br>Cholesterol-free | |

| | | | | | |
|---|---|---|---|---|---|
| | | | Inhibitor: <0.0001 | | Naïve vs LPS + IL-4: <0.0001<br><br>LPS + IL-4 vs NB-598: <0.0001<br><br>LPS + IL-4 vs Fluvastatin: <0.0001 |
| 6C | Two-way ANOVA | Interaction: 4.956<br><br>Media: 7.811<br><br>Inhibitor: 645.7 | Interaction: 0.0128<br><br>Media: 0.0130<br><br>Inhibitor: <0.0001 | Tukey's multiple comparisons test | Normal<br><br>Naïve vs LPS + IL-4: 0.3922<br><br>LPS + IL-4 vs NB-598: <0.0001<br><br>LPS + IL-4 vs Fluvastatin: <0.0001<br><br>Normal vs Cholesterol-free LPS + IL-4: 0.6252<br><br>Cholesterol-free<br><br>Naïve vs LPS + IL-4: 0.0012<br><br>LPS + IL-4 vs NB-598: <0.0001<br><br>LPS + IL-4 vs Fluvastatin: <0.0001 |
| 6D | Two-way ANOVA | Interaction: 11.74<br><br>Media: 1.331<br><br>Inhibitor: 76.74 | Interaction: 0.0003<br><br>Media: 0.2655<br><br>Inhibitor: <0.0001 | Tukey's multiple comparisons test | Normal<br><br>Naïve vs LPS + IL-4: <0.0001<br><br>LPS + IL-4 vs NB-598: 0.0388<br><br>LPS + IL-4 vs Fluvastatin: 0.0004<br><br>Normal vs Cholesterol-free LPS + IL-4: 0.0003<br><br>Cholesterol-free<br><br>Naïve vs LPS + IL-4: <0.0001 |

| | | | | | |
|---|---|---|---|---|---|
| | | | | | LPS + IL-4 vs NB-598: <0.0001<br><br>LPS + IL-4 vs Fluvastatin: <0.0001 |
| 6F | Two-way ANOVA | Interaction: 14.47<br><br>Media: 10.13<br><br>Inhibitor: 716.6 | Interaction: 0.0006<br><br>Media: 0.0079<br><br>Inhibitor: <0.0001 | Tukey's multiple comparisons test | Normal<br><br>Naïve vs LPS + IL-4: <0.0001<br><br>LPS + IL-4 vs FGTI-2734: <0.0001<br><br>Normal vs Cholesterol-free<br>LPS + IL-4: <0.0001<br><br>Cholesterol-free<br><br>Naïve vs LPS + IL-4: <0.0001<br><br>LPS + IL-4 vs FGTI-2734: <0.0001 |
| 6G | Two-way ANOVA | Interaction: 108.8<br><br>Media: 297.7<br><br>Inhibitor: 6586 | Interaction: <0.0001<br><br>Media: <0.0001<br><br>Inhibitor: <0.0001 | Tukey's multiple comparisons test | Normal<br><br>Naïve vs LPS + IL-4: 0.6468<br><br>LPS + IL-4 vs FGTI-2734: <0.0001<br><br>Normal vs Cholesterol-free<br>LPS + IL-4: 0.0001<br><br>Cholesterol-free<br><br>Naïve vs LPS + IL-4: 0.0037<br><br>LPS + IL-4 vs FGTI-2734: <0.0001 |
| 6H | Two-way ANOVA | Interaction: 42.17<br><br>Media: 10.39 | Interaction: <0.0001<br><br>Media: 0.0073 | Tukey's multiple comparisons test | Normal<br><br>Naïve vs LPS + IL-4: <0.0001<br><br>LPS + IL-4 vs FGTI-2734: <0.0001 |

| | | | | | | |
|---|---|---|---|---|---|---|
| | | Inhibitor: 422.7 | Inhibitor: <0.0001 | | Normal vs Cholesterol-free<br>LPS + IL-4: <0.0001<br><br>Cholesterol-free<br><br>Naïve vs LPS + IL-4: <0.0001<br><br>LPS + IL-4 vs FGTI-2734: <0.0001 | |
| 6J | One-way ANOVA | 33.93 | <0.0001 | Dunnett's multiple comparison's test | Normal<br><br>Naïve vs LPS + IL-4: <0.0001<br><br>LPS + IL-4 vs 10μM FTI-277: <0.0001<br><br>LPS + IL-4 vs 30μM FTI-277: <0.0001<br><br>LPS + IL-4 vs 10μM GGTI-298: <0.0001<br><br>LPS + IL-4 vs 30μM GGTI-298: <0.0001 | 0.1288 |
| 6K | One-way ANOVA | 139.2 | <0.0001 | Dunnett's multiple comparison's test | Normal<br><br>Naïve vs LPS + IL-4: 0.9988<br><br>LPS + IL-4 vs 10μM FTI-277: 0.0021<br><br>LPS + IL-4 vs 30μM FTI-277: <0.0001<br><br>LPS + IL-4 vs 10μM GGTI-298: <0.0001<br><br>LPS + IL-4 vs 30μM GGTI-298: <0.0001 | 0.3614 |
| 6L | One-way ANOVA | 66.44 | <0.0001 | Dunnett's multiple comparison's test | Normal<br><br>Naïve vs LPS + IL-4: <0.0001<br><br>LPS + IL-4 vs 10μM FTI-277: 0.0010 | 0.47 |

| | | | | | | |
|---|---|---|---|---|---|---|
| | | | | | LPS + IL-4 vs 30μM FTI-277: <0.0001<br><br>LPS + IL-4 vs 10μM GGTI-298: <0.0001<br><br>LPS + IL-4 vs 30μM GGTI-298: <0.0001 | |
| 6N | One-way ANOVA | 142.3 | <0.0001 | Dunnett's multiple comparison's test | <u>CF</u><br><br>Naïve vs LPS + IL-4: <0.0001<br><br>LPS + IL-4 vs 10μM FTI-277: <0.0001<br><br>LPS + IL-4 vs 30μM FTI-277: <0.0001<br><br>LPS + IL-4 vs 10μM GGTI-298: <0.0001<br><br>LPS + IL-4 vs 30μM GGTI-298: <0.0001 | 0.3492 |
| 6O | One-way ANOVA | 685.5 | <0.0001 | Dunnett's multiple comparison's test | <u>CF</u><br><br>Naïve vs LPS + IL-4: 0.1736<br><br>LPS + IL-4 vs 10μM FTI-277: 0.0790<br><br>LPS + IL-4 vs 30μM FTI-277: <0.0001<br><br>LPS + IL-4 vs 10μM GGTI-298: <0.0001<br><br>LPS + IL-4 vs 30μM GGTI-298: <0.0001 | 0.6699 |
| 6P | One-way ANOVA | 1591 | <0.0001 | Dunnett's multiple | <u>CF</u><br><br>Naïve vs LPS + IL-4: <0.0001<br><br>LPS + IL-4 vs 10μM FTI-277: <0.0001 | 0.4074 |

| | | | | | |
|---|---|---|---|---|---|
| | | | | comparison's test | LPS + IL-4 vs 30µM FTI-277: <0.0001<br><br>LPS + IL-4 vs 10µM GGTI-298: <0.0001<br><br>LPS + IL-4 vs 30µM GGTI-298: <0.0001 |

| Figure | Test | ANOVA F value | ANOVA P value | Test for multiple comparisons | Multiple comparisons (Adjusted P value) | Brown Forsythe test P value |
|---|---|---|---|---|---|---|
| 7A | Two-way ANOVA | Interaction: 10.82<br><br>Time: 27.12<br><br>Treatment: 38.02 | Interaction: 0.0021<br><br>Time: 0.0002<br><br>Treatment: <0.0001 | Sidak's multiple comparisons test | <u>Normal</u><br><br>LPS + IL-4 vs Fluvastatin (24 hours): 0.0209<br><br>Fluvastatin vs MVA (24 hours): 0.8260<br><br>LPS + IL-4 vs Fluvastatin (48 hours): <0.0001<br><br>Fluvastatin vs MVA (48 hours): <0.0001 | |
| 7B | Two-way ANOVA | Interaction: 15.32<br><br>Time: 14.26<br><br>Treatment: 195.1 | Interaction: 0.0005<br><br>Time: 0.0026<br><br>Treatment: <0.0001 | Sidak's multiple comparisons test | <u>CF</u><br><br>LPS + IL-4 vs Fluvastatin (24 hours): <0.0001<br><br>Fluvastatin vs MVA (24 hours): 0.8756<br><br>LPS + IL-4 vs Fluvastatin (48 hours): <0.0001<br><br>Fluvastatin vs MVA (48 hours): 0.2388 | |
| 7C | Two-way ANOVA | Interaction: 302.4<br><br>Media: 270.4 | Interaction: <0.0001<br><br>Media: <0.0001 | Sidak's multiple comparisons test | <u>Normal</u><br><br>LPS + IL-4 vs Fluvastatin: <0.0001<br><br>Fluvastatin vs MVA: <0.0001 | |

| | | | | | |
|---|---|---|---|---|---|
| | | Treatment: 922.5 | Treatment: <0.0001 | | <u>Cholesterol free</u><br><br>LPS + IL-4 vs Fluvastatin: <0.0001<br><br>Fluvastatin vs MVA: 0.0023 |
| 7D | Two-way ANOVA | Interaction: 18.29<br><br>Media: 0.5922<br><br>Treatment: 20.81 | Interaction: 0.0002<br><br>Media: 0.4565<br><br>Treatment: 0.0001 | Sidak's multiple comparisons test | <u>Normal</u><br><br>LPS + IL-4 vs Fluvastatin: 0.0129<br><br>Fluvastatin vs MVA: 0.0009<br><br><u>Cholesterol free</u><br><br>LPS + IL-4 vs Fluvastatin: 0.0006<br><br>Fluvastatin vs MVA: 0.7214 |
| 7F | Two-way ANOVA | Interaction: 93.12<br><br>Media: 0.02330<br><br>Treatment: 2594 | Interaction: <0.0001<br><br>Media: 0.8812<br><br>Treatment: <0.0001 | Sidak's multiple comparisons test | <u>Normal</u><br><br>LPS + IL-4 vs Fluvastatin: <0.0001<br><br>Fluvastatin vs MVA: <0.0001<br><br><u>Cholesterol free</u><br><br>LPS + IL-4 vs Fluvastatin: <0.0001<br><br>Fluvastatin vs MVA: 0.9810 |

| Figure | Test | ANOVA F value | ANOVA P value | Test for multiple comparisons | Multiple comparisons (Adjusted P value) | Brown Forsythe test P value |
|---|---|---|---|---|---|---|
| 8B | One-way ANOVA on log transformed data | 37.49 | <0.0001 | Dunnett's multiple comparison's test | <u>Normal</u><br><br>Fluvastatin vs LPS + IL-4: <0.0001<br><br>Fluvastatin vs MVA: 0.0003<br><br>Fluvastatin vs GGPP: <0.0001<br><br>Fluvastatin vs MVA + GGPP: <0.0001 | 0.9972 |
| 8C | One-way ANOVA | 63.63 | <0.0001 | Dunnett's multiple comparison's test | <u>Normal</u><br><br>Fluvastatin vs LPS + IL-4: <0.0001<br><br>Fluvastatin vs MVA: <0.0001<br><br>Fluvastatin vs GGPP: <0.0001<br><br>Fluvastatin vs MVA + GGPP: <0.0001 | 0.7134 |
| 8D | One-way ANOVA | 52.03 | <0.0001 | Dunnett's multiple comparison's test | <u>Normal</u><br><br>Fluvastatin vs LPS + IL-4: <0.0001<br><br>Fluvastatin vs MVA: <0.0001<br><br>Fluvastatin vs GGPP: <0.0001<br><br>Fluvastatin vs MVA + GGPP: <0.0001 | 0.8877 |
| | | | | | <u>CF</u><br><br>Fluvastatin vs LPS + IL-4: <0.0001 | |

| | | | | | | |
|---|---|---|---|---|---|---|
| 8F | One-way ANOVA on log transformed data | 38.63 | <0.0001 | Dunnett's multiple comparison's test | Fluvastatin vs MVA: 0.248<br><br>Fluvastatin vs GGPP: 0.4568<br><br>Fluvastatin vs MVA + GGPP: 0.9404 | 0.9069 |
| 8G | One-way ANOVA | 3461 | <0.0001 | Dunnett's multiple comparison's test | <u>CF</u><br><br>Fluvastatin vs LPS + IL-4: <0.0001<br><br>Fluvastatin vs MVA: <0.0001<br><br>Fluvastatin vs GGPP: <0.0001<br><br>Fluvastatin vs MVA + GGPP: <0.0001 | 0.8872 |
| 8H | One-way ANOVA | 23.61 | <0.0001 | Dunnett's multiple comparison's test | <u>CF</u><br><br>Fluvastatin vs LPS + IL-4: 0.0005<br><br>Fluvastatin vs MVA: 0.0893<br><br>Fluvastatin vs GGPP: 0.1305<br><br>Fluvastatin vs MVA + GGPP: 0.9897 | 0.7828 |
| 8I | Two-way ANOVA | Interaction: 19.16<br><br>Media: 219.9<br><br>Inhibitor: 105.7 | Interaction: 0.0002<br><br>Media: <0.0001<br><br>Inhibitor: <0.0001 | Sidak's multiple comparisons test | <u>Normal</u><br><br>LPS + IL-4 vs Fluvastatin (24 hours): 0.0003<br><br>Fluvastatin vs GGPP (24 hours): 0.0042<br><br>LPS + IL-4 vs Fluvastatin (48 hours): <0.0001<br><br>Fluvastatin vs GGPP (48 hours): <0.0001 | |

| | | | | | |
|---|---|---|---|---|---|
| 8J | Two-way ANOVA | Interaction: 96.40<br><br>Media: 66.99<br><br>Inhibitor: 388.8 | Interaction: <0.0001<br><br>Media: <0.0001<br><br>Inhibitor: <0.0001 | Sidak's multiple comparisons test | CF<br><br>LPS + IL-4 vs Fluvastatin (24 hours): <0.0001<br><br>Fluvastatin vs GGPP (24 hours): >0.9999<br><br>LPS + IL-4 vs Fluvastatin (48 hours): <0.0001<br><br>Fluvastatin vs GGPP (48 hours): >0.9999 |

| Figure | Test | ANOVA F value | ANOVA P value | Test for multiple comparisons | Multiple comparisons (Adjusted P value) | Brown Forsythe test P value |
|---|---|---|---|---|---|---|
| 9B | Two-way ANOVA | Interaction: 601.8<br><br>Generation: 1207<br><br>Treatment: 2.930 | Interaction: <0.0001<br><br>Generation: <0.0001<br><br>Treatment: 0.0218 | Sidak's multiple comparisons test | Generation 1<br>Naïve vs LPS + IL-4: <0.0001<br>LPS + IL-4 vs PD18352: <0.0001<br>LPS + IL-4 vs VX745: 0.9950<br>LPS + IL-4 vs PD18352 + VX745: <0.0001<br>LPS + IL-4 vs Rapamycin: <0.0001<br><br>Generation 2<br>LPS + IL-4 vs PD18352: >0.9999<br>LPS + IL-4 vs VX745: 0.9238<br>LPS + IL-4 vs PD18352 + VX745: <0.0001<br>LPS + IL-4 vs Rapamycin: >0.9999<br><br>Generation 3<br>LPS + IL-4 vs PD18352: >0.9999<br>LPS + IL-4 vs VX745: 0.0134<br>LPS + IL-4 vs PD18352 + VX745: <0.0001<br>LPS + IL-4 vs Rapamycin: <0.0001<br><br>Generation 2 | |

| | | | | | | |
|---|---|---|---|---|---|---|
| | | | | | LPS + IL-4 vs PD18352: <0.0001<br>LPS + IL-4 vs VX745: <0.0001 | |
| 9C | One-way ANOVA | 64.25 | <0.0001 | Dunnett's multiple comparison's test | Naïve vs LPS + IL-4: <0.0001<br><br>LPS + IL-4 vs PD18352: <0.0001<br><br>LPS + IL-4 vs VX745: 0.0409<br><br>LPS + IL-4 vs PD18352 + VX745: <0.0001<br><br>LPS + IL-4 vs Rapamycin: <0.0001 | 0.4358 |
| 9D | One-way ANOVA | 123.7 | <0.0001 | Dunnett's multiple comparison's test | Naïve vs LPS + IL-4: 0.0034<br><br>LPS + IL-4 vs PD18352: <0.0001<br><br>LPS + IL-4 vs VX745: >0.9999<br><br>LPS + IL-4 vs PD18352 + VX745: <0.0001<br><br>LPS + IL-4 vs Rapamycin: <0.0001 | 0.2771 |
| 9E | One-way ANOVA | 300.9 | <0.0001 | Dunnett's multiple comparison's test | Naïve vs LPS + IL-4: <0.0001<br><br>LPS + IL-4 vs PD18352: <0.0001<br><br>LPS + IL-4 vs VX745: 0.5399<br><br>LPS + IL-4 vs PD18352 + VX745: <0.0001<br><br>LPS + IL-4 vs Rapamycin: <0.0001 | 0.5122 |

| | | | | | | |
|---|---|---|---|---|---|---|
| 9F | One-way ANOVA | 31.90 | <0.0001 | Dunnett's multiple comparison's test | Naïve vs LPS + IL-4: <0.0001<br><br>LPS + IL-4 vs PD18352: 0.0001<br><br>LPS + IL-4 vs VX745: 0.0013<br><br>LPS + IL-4 vs PD18352 + VX745: <0.0001<br><br>LPS + IL-4 vs Rapamycin: <0.0001 | 0.7256 |

| Figure | Test | ANOVA F value | ANOVA P value | Test for multiple comparisons | Multiple comparisons (Adjusted P value) | Brown Forsythe test P value |
|---|---|---|---|---|---|---|
| 10A | Two-way ANOVA | Interaction: 14.67<br><br>Media: 66.83<br><br>Inhibitor: 711.2 | Interaction: <0.0001<br><br>Media: <0.0001<br><br>Inhibitor: <0.0001 | Tukey's multiple comparisons test | <u>Normal</u><br><br>Naïve vs IL-4: <0.0001<br><br>Naïve vs LPS: <0.0001<br><br>Naïve vs LPS + IL-4: <0.0001<br><br><u>Cholesterol-free</u><br><br>Naïve vs IL-4: <0.0001<br><br>Naïve vs LPS: <0.0001<br><br>Naïve vs LPS + IL-4: <0.0001 | |
| 10B | One-way ANOVA | 284.4 | <0.0001 | Dunnett's multiple comparison's test | Naïve vs LPS: <0.0001<br><br>Naïve vs Resiquimod: <0.0001<br><br>Naïve vs CpG: <0.0001<br><br>Naïve vs Anti-IgM: <0.0001<br><br>Naïve vs CD40L: <0.0001 | 0.8282 |

| | | | | | |
|---|---|---|---|---|---|
| 10C | Two-way ANOVA | Interaction: 89.65<br><br>Inhibitor: 2014<br><br>Stimuli: 80.20 | Interaction: <0.0001<br><br>Inhibitor: <0.0001<br><br>Stimuli: <0.0001 | Tukey's multiple comparisons test | Naïve vs LPS: 0.0178<br><br>LPS vs Fluvastatin: <0.0001<br><br>Naïve vs Anti-IgM: 0.9997<br><br>Anti-IgM vs Fluvastatin: <0.0001<br><br>Naïve vs CD40L: >0.9999<br><br>CD40L vs Fluvastatin: <0.0001<br><br>Naïve vs Resiquimod: 0.9975<br><br>Resiquimod vs Fluvastatin: <0.0001<br><br>Naïve vs CpG: 0.9396<br><br>CpG vs Fluvastatin: <0.0001 |
| 10E | Two-way ANOVA on log transformed data | Interaction: 94.68<br><br>Inhibitor: 1776<br><br>Stimuli: 40.29 | Interaction: <0.0001<br><br>Inhibitor: <0.0001<br><br>Stimuli: <0.0001 | Tukey's multiple comparisons test | Naïve vs LPS: <0.0001<br><br>LPS vs Fluvastatin: <0.0001<br><br>Naïve vs Anti-IgM: 0.0535<br><br>Anti-IgM vs Fluvastatin: <0.0001<br><br>Naïve vs CD40L: >0.9999<br><br>CD40L vs Fluvastatin: <0.0001<br><br>Naïve vs Resiquimod: 0.0006<br><br>Resiquimod vs Fluvastatin: <0.0001<br><br>Naïve vs CpG: 0.0002<br><br>CpG vs Fluvastatin: <0.0001 |

| Figure | Test | ANOVA F value | ANOVA P value | Test for multiple comparisons | Multiple comparisons (Adjusted P value) | Brown Forsythe test P value |
|---|---|---|---|---|---|---|
| Figure S3C | One-way ANOVA | 71.82 | <0.0001 | Dunnett's multiple comparison's test | Naïve vs naïve + IFN-β: <0.0001<br><br>LPS vs LPS + IFN-β: <0.0001 | 0.3666 |

| Figure | Test | ANOVA F value | ANOVA P value | Test for multiple comparisons | Multiple comparisons (Adjusted P value) | Brown Forsythe test P value |
|---|---|---|---|---|---|---|
| Figure S4B | One-way ANOVA | 168.6 | <0.0001 | Dunnett's multiple comparison's test | Naïve vs LPS: <0.0001<br><br>Naïve vs Resiquimod: <0.0001<br><br>Naïve vs CpG: <0.0001<br><br>Naïve vs Anti-IgM: <0.0001<br><br>Naïve vs CD40L: <0.0001 | 0.7622 |

| Figure | Test | ANOVA F value | ANOVA P value | Test for multiple comparisons | Multiple comparisons (Adjusted P value) | Brown Forsythe test P value |
|---|---|---|---|---|---|---|
| Figure S6A | One-way ANOVA | 44.94 | <0.0001 | Dunnett's multiple comparison's test | Naïve vs LPS (2 hours): 0.0425<br><br>Naïve vs LPS (4 hours): <0.0001<br><br>Naïve vs LPS (8 hours): <0.0001 | 0.0846 |
| Figure S6B | One-way ANOVA | 14.89 | <0.0001 | Dunnett's multiple comparison's test | Naïve vs LPS (2 hours): 0.2765<br><br>Naïve vs LPS (4 hours): 0.0019<br><br>Naïve vs LPS (8 hours): 0.0053 | 0.2086 |

| | | | | | | |
|---|---|---|---|---|---|---|
| Figure S6C | One-way ANOVA | 55.65 | <0.0001 | Dunnett's multiple comparison's test | Naïve vs LPS (2 hours): <0.0001<br>Naïve vs LPS (4 hours): <0.0001<br>Naïve vs LPS (8 hours): 0.0004 | 0.2816 |
| Figure S6D | One-way ANOVA | 75.20 | <0.0001 | Dunnett's multiple comparison's test | Naïve vs LPS (2 hours): <0.0001<br>Naïve vs LPS (4 hours): <0.0001<br>Naïve vs LPS (8 hours): 0.0008 | 0.3293 |
| Figure S6E | One-way ANOVA | 46.20 | <0.0001 | Dunnett's multiple comparison's test | Naïve vs LPS (2 hours): <0.0001<br>Naïve vs LPS (4 hours): <0.0001<br>Naïve vs LPS (8 hours): 0.1394 | 0.5125 |
| Figure S6F | One-way ANOVA | 3.915 | 0.0319 | Dunnett's multiple comparison's test | Naïve vs LPS (2 hours): 0.2252<br>Naïve vs LPS (4 hours): 0.4463<br>Naïve vs LPS (8 hours): 0.3083 | 0.1257 |

| Figure | Test | ANOVA F value | ANOVA P value | Test for multiple comparisons | Multiple comparisons (Adjusted P value) | Brown Forsythe test P value |
|---|---|---|---|---|---|---|

| | | | | | | |
|---|---|---|---|---|---|---|
| Figure S8A | One-way ANOVA | 110.1 | <0.0001 | Dunnett's multiple comparison's test | LPS + IL-4 vs 0.1μM Fluvastatin: >0.9999<br><br>LPS + IL-4 vs 1μM Fluvastatin: 0.6755<br><br>LPS + IL-4 vs 10μM Fluvastatin: 0.0642<br><br>LPS + IL-4 vs 50μM Fluvastatin: <0.0001<br><br>LPS + IL-4 vs 100μM Fluvastatin: <0.0001 | 0.2767 |
| Figure S8B | One-way ANOVA | 24.24 | <0.0001 | Dunnett's multiple comparison's test | LPS + IL-4 vs 0.1μM Fluvastatin: 0.3002<br><br>LPS + IL-4 vs 1μM Fluvastatin: 0.1048<br><br>LPS + IL-4 vs 10μM Fluvastatin: 0.0055<br><br>LPS + IL-4 vs 50μM Fluvastatin: 0.0041<br><br>LPS + IL-4 vs 100μM Fluvastatin: 0.9990 | 0.5532 |
| Figure S8D | One-way ANOVA | 18.37 | <0.0001 | Dunnett's multiple comparison's test | LPS + IL-4 vs 0.1μM NB-598: 0.9921<br><br>LPS + IL-4 vs 1μM NB-598: 0.9641<br><br>LPS + IL-4 vs 10μM NB-598: 0.0215<br><br>LPS + IL-4 vs 50μM NB-598: 0.0016<br><br>LPS + IL-4 vs 100μM NB-598: <0.0001 | 0.1551 |
| | | | | Dunnett's multiple | LPS + IL-4 vs 0.1μM NB-598: 0.5711<br><br>LPS + IL-4 vs 1μM NB-598: 0. 0384 | |

| | | | | | | |
|---|---|---|---|---|---|---|
| Figure S8E | One-way ANOVA | 35.70 | <0.0001 | comparison's test | LPS + IL-4 vs 10μM NB-598: 0.2167<br><br>LPS + IL-4 vs 50μM NB-598: 0.0727<br><br>LPS + IL-4 vs 100μM NB-598: 0.0003 | 0.4192 |
| Figure S8G | One-way ANOVA | 80.33 | <0.0001 | Dunnett's multiple comparison's test | LPS + IL-4 vs 0.1μM Rosuvastatin: 0.0078<br><br>LPS + IL-4 vs 1μM Rosuvastatin: 0.0034<br><br>LPS + IL-4 vs 10μM Rosuvastatin: <0.0001<br><br>LPS + IL-4 vs 50μM Rosuvastatin: <0.0001<br><br>LPS + IL-4 vs 100μM Rosuvastatin: <0.0001 | 0.8511 |
| Figure S8H | One-way ANOVA | 37.33 | <0.0001 | Dunnett's multiple comparison's test | LPS + IL-4 vs 0.1μM Rosuvastatin: 0.4964<br><br>LPS + IL-4 vs 1μM Rosuvastatin: 0.5414<br><br>LPS + IL-4 vs 10μM Rosuvastatin: 0.0767<br><br>LPS + IL-4 vs 50μM Rosuvastatin: 0.0052<br><br>LPS + IL-4 vs 100μM Rosuvastatin: 0.0005 | 0.2395 |
| Figure S8J | One-way ANOVA | 197.4 | <0.0001 | Dunnett's multiple comparison's test | LPS + IL-4 vs 0.1μM Rosuvastatin: 0.0039<br><br>LPS + IL-4 vs 1μM Rosuvastatin: <0.0001<br><br>LPS + IL-4 vs 10μM Rosuvastatin: <0.0001<br><br>LPS + IL-4 vs 50μM Rosuvastatin: <0.0001 | 0.5381 |

| | | | | | |
|---|---|---|---|---|---|
| | | | | | LPS + IL-4 vs 100µM Rosuvastatin: <0.0001 |

| Figure | Test | ANOVA F value | ANOVA P value | Test for multiple comparisons | Multiple comparisons (Adjusted P value) | Brown Forsythe test P value |
|---|---|---|---|---|---|---|
| Figure S9G | One-way ANOVA | 31.18 | <0.0001 | Dunnett's multiple comparison's test | LPS + IL-4 vs Fluvastatin (normal): 0.0003<br><br>LPS + IL-4 vs LPS + IL-4 (normal/CF): 0.0047<br><br>LPS + IL-4 vs Fluvastatin (CF): <0.0001 | 0.2643 |
| Figure S9H | One-way ANOVA | 20.61 | <0.0001 | Dunnett's multiple comparison's test | LPS + IL-4 vs Fluvastatin (normal): 0.0022<br><br>LPS + IL-4 vs LPS + IL-4 (normal/CF): 0.0066<br><br>LPS + IL-4 vs Fluvastatin (CF): <0.0001 | 0.1823 |
| Figure S9I | One-way ANOVA | 55.50 | <0.0001 | Dunnett's multiple comparison's test | LPS + IL-4 vs Fluvastatin (normal): 0.0006<br><br>LPS + IL-4 vs LPS + IL-4 (normal/CF): <0.0001<br><br>LPS + IL-4 vs Fluvastatin (CF): <0.0001 | 0.1322 |

| Figure | Test | ANOVA F value | ANOVA P value | Test for multiple comparisons | Multiple comparisons (Adjusted P value) | Brown Forsythe test P value |
|---|---|---|---|---|---|---|
| | | | | | Naïve vs LPS + IL-4: <0.0001 | |

| | | | | | | |
|---|---|---|---|---|---|---|
| Figure S10A | One-way ANOVA | 109.2 | <0.0001 | Dunnett's multiple comparison's test | LPS + IL-4 vs 1mM MVA: 0.9879<br><br>LPS + IL-4 vs 2mM MVA: 0.2461<br><br>LPS + IL-4 vs 4mM MVA: <0.0001 | 0.6727 |
| Figure S10B | One-way ANOVA | 105.7 | <0.0001 | Dunnett's multiple comparison's test | Naïve vs LPS + IL-4: 0.0187<br><br>LPS + IL-4 vs 1mM MVA: 0.0625<br><br>LPS + IL-4 vs 2mM MVA: <0.0001<br><br>LPS + IL-4 vs 4mM MVA: <0.0001 | 0.3874 |

| Figure | Test | ANOVA F value | ANOVA P value | Test for multiple comparisons | Multiple comparisons (Adjusted P value) | Brown Forsythe test P value |
|---|---|---|---|---|---|---|
| Figure S12A | One-way ANOVA | 165.6 | <0.0001 | Dunnett's multiple comparison's test | Naïve vs LPS + IL-4: <0.0001<br><br>LPS + IL-4 vs rapamycin: 0.0137 | 0.6914 |
| Figure S12B | One-way ANOVA | 345.0 | <0.0001 | Dunnett's multiple comparison's test | Naïve vs LPS + IL-4: <0.0001<br><br>LPS + IL-4 vs rapamycin: 0.0001 | 0.2597 |
